# Regulatory architectures optimized for rapid evolution of gene expression

**DOI:** 10.1101/2025.06.10.658850

**Authors:** Réka Borbély, Gašper Tkačik

## Abstract

Cis-regulatory elements (CREs), such as enhancers and promoters, control gene expression by binding regulatory proteins. In contrast to their bacterial counterparts, eukaryotic CREs typically bind transcription factors with short recognition motifs across multiple functional yet often weak binding sites. The evolutionary origin of this genetic architecture remains unclear. Here we study adaptive evolution of entire CREs under selection for controllable gene expression. In a biophysical toy model that recapitulates the essential non-linearities of eukaryotic regulation, a regulatory phenotype requires a gene to be active when its CRE binds cognate TFs, yet inactive in the presence of noncognate TFs that can cause deleterious crosstalk. We explore CRE evolutionary outcomes utilizing “optimize-to-adapt” approach suggested by the theory presented in a companion paper. In this approach, CRE evolution is simulated explicitly at the sequence level, while the parameters of the biophysical model itself – i.e., the properties of the replicated genotype-phenotype map – are numerically optimized for evolvability of CREs. In the optimal regime, selection navigates the tradeoff between slowly evolving strong and long binding sites (which guarantee low crosstalk), and rapidly evolving multiple short and weak binding sites (which necessitate diffuse selection against too much noncognate binding across the entire CRE). When we further explore various scenarios for cooperative regulation, we find that the optimal regime predicts a diversity of strong and weak, yet always short, binding sites and favors “synergistic activation” of transcription, as reported empirically for eukaryotes. These results showcase how information theory can link evolutionary dynamics with biophysical constraints to rationalize – and possibly even predict – optimal regulatory architectures.

---

Regulatory sequences in the DNA, such as promoters and enhancers—often referred to generically as “cis-regulatory elements” (CREs)—specify gene expression programs essential to the life of simple prokaryotic cells as well as complex multi-cellular organisms. For housekeeping genes, these sequences might encode a preferred constant gene expression level; for developmental genes, in contrast, they often encode elaborate phenotypes that condition target gene expression on developmental time, position within the embryo, tissue type, or external chemical cues. Either way, these sequences, which typically comprise several hundred base pairs, are read out by transcription factors (TFs), proteins that recognize ∼6 − 20 bp sequence motifs and bind to closely matching transcription factor binding sites (BSs) embedded within the CREs. The biophysical basis for this molecular recognition is generally well-understood [1]. The resulting control of gene expression has been worked out in detail for prokaryotes [2–4], whereas for eukaryotes and especially metazoans important mechanistic questions remain open. That is why, even though we posses a detailed list of relevant molecular players and interactions, most quantitative models for eukaryotic regulation are either purely statistical [5], or biophysically-inspired yet of a highly simplified variety [6].

Despite the complexity of the mechanisms involved, extensive genomic evidence suggests that the evolution of regulatory sequences is pervasive, plastic, and rapid. BSs can turn over quickly within and across populations [7, 8]; mutations in BSs are a way to rewire the regulatory networks [9]; TF gene duplications followed by divergence of their motifs offer an accessible route to functional specialization [10]; entirely random sequences may be one or only a few mutations away from functional promoters [11, 12]; individual point mutations in eukaryotic enhancers can quantitatively change gene expression phenotypes [13–15]; and genome-wide association studies often reveal a high fraction of significant hits in regulatory sequences for various traits of interest [16, 17]. Taken together, regulatory sequences appear to be a suitable substrate for rapid evolutionary adaptation across different kingdoms of life [18–20].

A fundamental question asks which conditions are necessary and sufficient for regulatory sequences to be so evolvable. Ultimately, this is a question about the molecular regulatory machinery that reads out CREs and maps them into controllable gene expression phenotypes, i.e., expression levels conditional on environmental or cell-internal states [21]. All CREs in an organism, together with their shared regulatory machinery, can be seen as evolving on a single, unchanging, global genotype-phenotype map for the entire regulatory system. A theoretical framework presented in the companion paper [22] suggests that we can view this same process from a local viewpoint: as a genotype-phenotype map for a single CRE that we focus on and whose properties themselves can evolve, under the condition that we self-consistently apply these “replicated GP map” properties to all CREs in the organism. We show that evolution will tune the properties of such “replicated genotype-phenotype maps” so as to maximize the number of fit genotypes (thereby typically making these maps most evolvable) under quite general conditions. Here, we reformulate this theoretical result that strictly holds only in the evolutionary steady state, into a hypothesis about evolutionary dynamics: namely, that evolution will have tuned the biophysical parameters of the shared regulatory machinery reading out CREs such that the adaptation of these CREs is most rapid. In what follows, we will state this optimality hypothesis precisely and explore its predictions using extensive numerical simulations in a newly-proposed “optimize-to-adapt” approach.

Before proceeding, we note that the attractive picture of pervasive, plastic, and rapid regulatory sequence evolution is not without unresolved puzzles and apparent paradoxes. These emerge when genomic findings, biophysical constraints, and evolutionary theory are brought into contact. **First**, population genetics predicts long waiting times for BSs to emerge from random sequence, which appears difficult to reconcile with genomic observations except at exceedingly high selection strength [23, 24]. **Second**, eukaryotic TFs recognize surprisingly short (6− 10 bp) motifs despite drastically larger genomic background relative to prokaryotes (that utilize 10− 20 bp long motifs) [25], thus incurring a larger risk of regulatory crosstalk due to noncognate binding [6, 26–28]. While the broad range of 6 − 20 bp motif lengths has been identified as evolutionarily favored [29], the reasons for systematic and counterintuitive pro-vs eukaryotic differences within this range remain unclear. **Third**, in contrast to the established paradigm that views CREs as nearly random sequences which contain well-delineated, strong and often conserved BSs, weak binding throughout eukaryotic CREs seems to be widespread and functionally important in natural and synthetic settings [5, 30, 31]. **Fourth**, while theoretical arguments clarify that mismatches between the preferred motif and individual BSs can be expected [32], they also suggest that selection would be most effective when regulatory information is spread across very long binding sites (unlike what we see) so that each individual base pair only conveys weak specificity and thus undergoes nearly neutral evolution [33, 34].

To work out the consequences of our optimality hypothesis and shed light on the unresolved puzzles listed above, we sought a modeling framework that could capture the essential regulatory biophysics of eukaryotes. Such models must be heavily stylized versions of reality in order to be interpretable and amenable to direct simulation as well as theoretical reasoning. Nevertheless, we insisted that the model satisfy the following minimal requirements: **(i)** describe the evolution of entire CREs at the sequence level; **(ii)** enable multiple BSs of various strength for different TFs to emerge and affect, adaptively or deleteriously, the “regulatory phenotype” under selection; **(iii)** permit the incorporation of various modes of cooperative regulation; **(iv)** be specified by a small number of physical parameters whose typical values and ranges are either known or could be set by the postulated optimality principle.

In the “optimize-to-adapt” approach, motivated by our recent theory [22] and deployed here, CRE adaptation is simulated explicitly at the sequence level, while the parameters of the shared regulatory machinery reading out the CREs are numerically optimized to speed up the CRE adaptation. To the best of our knowledge, such an approach has not been attempted before. Optimization of regulatory parameters can be interpreted as a (much slower) adaptation of the replicated genotype-phenotype map itself, in order to make CREs most evolvable [22, 35]. Importantly, in this approach, evolvability is not treated only as a qualitative descriptor or a *post-hoc* statistic, but rather as a predictive mathematical optimization principle that will determine the properties of the replicated genotype-phenotype map, thereby simultaneously (re)shaping all evolved cis-regulatory elements.

## A biophysically-inspired model for CRE evolution

Here we briefly introduce our minimal model of gene regulation that satisfies the four requirements posed in the Introduction (for details, please see SI Appendix Section 1A).

We consider the evolution of a representative cis-regulatory element **s**, where *s*_*i*_ ∈ {*A, C, G, T*} for *i* = 1, …, *L*, in which multiple, possibly overlapping, binding sites of length *ℓ* ≪ *L* for *n* + 1 different transcription factors can emerge through point mutations. As we clarify later, one of these TFs will be denoted as “cognate” and the remaining *n* as “noncognate.” In the simplest model version, we consider TF binding as non-interacting (i.e., there is no intrinsic cooperativity, synergistic activation, or steric occlusion effects) and limit ourselves to activators only. While we relax some of these restrictions later in the paper, we note that even the simplest model version shows rich phenomenology while remaining interpretable.

Each TF *j* is characterized by its preferred sequence motif (or consensus sequence) **m**_*j*_. We adopt the mismatch energy model to calculate the equilibrium binding probability *ρ*_*ij*_ of a TF to each possible *ℓ*-base-pair window in the CRE **s**, via the “binding nonlinearity” as follows: *ρ*_*ij*_ = *c*_*j*_*/* (*c*_*j*_ + exp(*εk*_*i,j*_)) (Fig. 1A) [23, 24]. Here, *c*_*j*_ is the concentration of the TF *j* in units where the dissociation constant to the strongest possible binding site would be 1, and *k*_*i,j*_ is the integer number of mismatches between the motif **m**_*j*_ and the actual CRE sequence starting at location *i* (i.e., {*s*_*i*:*i*+*ℓ−*1_}). Thus, *k* = 0, the minimal possible mismatch, indicates a perfect BS that exactly matches the TF’s preferred motif and is most strongly bound; *k* = *ℓ*, the maximal possible mismatch, will lead to weakest binding. Parameter *ε*, known as the “mismatch energy,” is confined to the range 3 in natural Boltzmann (*k*_*B*_*T*) units, set by the energy scale of hydrogen bonds between the TF’s DNA binding domain and the sequence.

**Figure 1.**
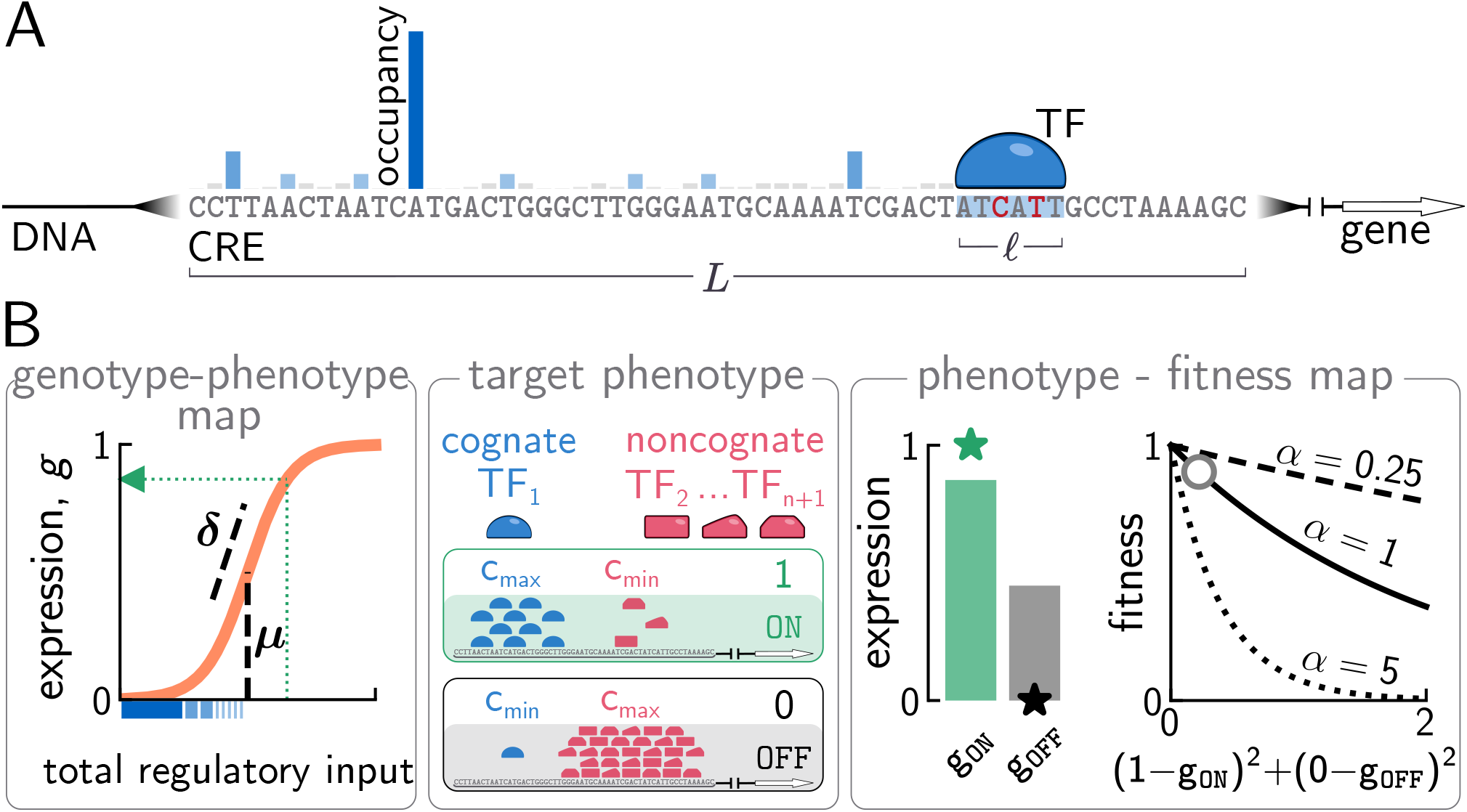
A toy model for eukaryotic gene regulation as a nonlinear multi-layer genotype-phenotype-fitness map. **(A)** The occupancy of each possible binding site (blue vertical bar) within the CRE of length *L* bp is determined by the number of mismatches it harbors in a BS of *ℓ* bp compared to the consensus sequence of the TF (blue half-oval), the mismatch energy *ε*, and the TF concentration. Total regulatory input is the sum over all site occupancies. **(B)** Total regulatory inputs for cognate (blue) or noncognate (red) factors determine gene expression via a GP map, parameterized by the slope *δ* and threshold *µ*. Fitness, Eq. (3), favors high expression (*g*_ON_, green bar) in ON environments where cognate TF is present at high concentration, and low expression (*g*_OFF_, gray bar) in OFF environments where cognate TF is not present but noncognate TFs may be, by penalizing deviations of expression (bars) from its desired target (stars, *G*_*e*_ = 0, 1 for *e* = {OFF, ON}, respectively) with tunable strength *α* (solid, dashed, dotted curves; see text).

The assumption of independent TF binding that we start with leads us to define the total “regulatory input” of TF *j* as a summation of binding probabilities over all locations within the CRE:

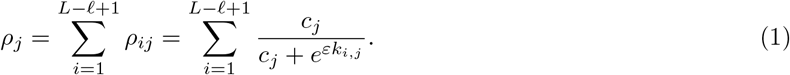

This standard, thermodynamically-motivated binding nonlinearity [23] with additive contributions over the entire sequence considered, as in Eq (1), has been used before to model gene expression from bacterial promoters (e.g., with *ρ* being interpreted as normalized gene expression where *j* = 1 stood for a single activator TF or the RNA polymerase itself) [12, 24]. Realistic models introduce a differential impact of TFs binding at various positions or orientations within the CRE, which is supported by quantitative data, but would not materially affect our evolutionary outcomes.

To move beyond independent TF binding and challenge the additivity assumption, one can modify Eq (1) so as to recapitulate various proposed cooperativity mechanisms (SI Appendix Section 1B). One such mechanism is the “homodimer cooperativity” characteristic of prokaryotes, where the binding at individual site *i* is modeled by Hill functions with steepness coefficient larger than one. Another is “synergistic activation” reported for eukaryotes, where the entire regulatory input *ρ*_*j*_ is raised to some power *η >* 1 before it is fed into the model for gene expression, Eq (2). We return to the evolutionary impact of these mechanisms in the Results section.

In contrast to previous modeling work, our approach combines regulatory inputs from different (cognate and noncognate) TFs and passes them through the second sigmoid “activation nonlinearity” in order to yield the resulting gene expression of the regulated gene. We introduce this nonlinearity phenomenologically, but note that it could result from different plausible biophysical mechanisms [6, 36, 37]. We denote gene expression level by *g* (normalized to 1 at maximum) and write it as a function of *n* + 1 TF concentrations **c** = {*c*_*j*_}:

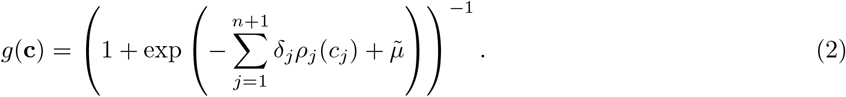

Here, parameters *δ*_*j*_ set the scale of various regulatory inputs (i.e., *δ*_*j*_ is the maximal contribution that a single perfect BS for factor *j* could exert); *δ*_*j*_ *>* 0 for activators, and *δ*_*j*_ *<* 0 could be used to model repressor TFs, which we leave open for future work. For simplicity, we assume in the following that all TFs contribute with the same strength, i.e., *δ*_*j*_ = *δ >* 0 for all *j*. With this choice, we rewrite *g*(**c**) = (1 + exp(− *δ*(*ρ* − *µ*))^*−*1^, where *ρ* now stands for the sum of contributions across all (*n* + 1) TFs.

Figure 1B sketches the essential role of the activation nonlinearity: it allows the maximal gene expression level to be achieved by different combinations of weak and strong BSs in the CRE, depending on the threshold *µ* that controls how much total regulatory input is needed to switch the gene ON.

Equation (2) specifies our regulatory phenotype encoded by the CRE **s**, i.e., the gene expression level *g* as a function of TF concentrations **c**. To model regulation, we now consider different cellular environments *e*: in each environment, a subset of TFs is “present,” i.e., their concentration is high, *c*_*j,e*_ = *c*_max_, while the other TFs are “absent,” i.e., their concentrations are low *c*_*j,e*_ = *c*_min_ ≪ *c*_max_. These environments can be interpreted broadly: they can be thought of as presence / absence of external nutrients, stressors, or other agents that cause gene expression response change for, e.g., unicellular organisms; alternatively, these could be external morphogenetic signals during development, or different tissue types in multicellular organisms. For our framework, what matters is that cells need to respond to these different environments *e* appropriately, by up-or down-regulating genes according to programs encoded in those genes’ CREs. For simplicity we assume that gene expression levels adjust instantaneously to environmental changes (as in previous work [10, 27, 28]) and that environments occur with equal probability (SI Appendix Section 1C).

Fitness is maximized when the gene is ON in the environments *e* where the cognate TF, enumerated as *j* = 1, is present (we refer to these environments as “ON environments” for short); the gene must also remain OFF in the “OFF environments” *e* where the cognate TF is absent, irrespective of the state of the other *n* noncognate TFs. In other words, it is not enough for the gene to be indiscriminately ON (which could trivially be achieved by making *µ* very small irrespective of the sequence); the gene must be expressed selectively and specifically only in response to its cognate signal. Any gene expression that is uncorrelated with the cognate signal is, in our model, considered to be a deleterious result of “regulatory crosstalk” – an interference due to the binding of noncognate TFs [27, 28].

Evolutionary selection for a reliable regulatory phenotype is naturally captured by the following fitness function:

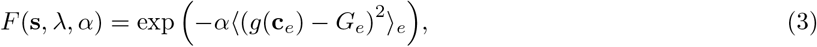

where we used the standard squared fitness cost for deviations of actual gene expression *g* in environment *e* from the desired (target) expression, *G*_*e*_ (which is either 0 or 1, depending on the presence or absence of cognate TF in each *e*), averaged over the possible environments *e* (SI Appendix Section 1C). While we assumed equal selection against both kinds of deviations for simplicity – i.e., expression in environments where there shouldn’t be any; no expression in environments that require it – this symmetry could easily be removed.

In the presented setup, the parameters naturally cluster into two groups. On the one hand, we have the population genetic parameters. Population size is *N*, which we assume does not change in time, while *α* implicitly controls the selection strength. We call this control “implicit” because evolution on our strongly nonlinear GP map changes fitness effects as it unfolds, so that the selection coefficients have nontrivial dynamics themselves (SI Fig. S1), as studied previously in simple [38] or realistic evolutionary models [21]. Nevertheless, we will see that our results will be largely independent of *α* in the initial steps of our analysis, while later higher *α* will correlate with stronger effective selection strength. On the other hand, the “regulatory parameter” vector *λ* = {*δ, µ, ε, c*_max_, *R* = log10 (*c*_max_*/c*_min_), *η*, **m**_*j*_, *L*} collects together biophysical parameters that specify the replicated genotype-phenotype map. Jointly, these two groups of parameters define the fitness *F* for every CRE, as given by Eqs (1,2,3), and govern the evolutionary dynamics of CREs.

Our parameterization makes the correspondence with the theory laid out in the companion paper explicit [22]. When regulatory parameters *λ* control replicated GP maps for multiple CREs in the same organism – among which we here choose to track the evolution of a single, typical CRE **s** – selection for regulatory function will implicitly yet strongly favor values of *λ* that maximize the number of fit genotypes. In the evolutionary steady-state with many CREs in the weak-mutation limit (i.e., under “the fixed states approximation”), this theoretical optimization principle for replicated GP maps holds exactly. Outside of that formal limit, it provides a strong expectation anchor for all our explorations below.

We study the evolutionary dynamics of CREs in the weak-mutation limit, since we are dealing with relatively short sequences of *L* ∼ 100 bp; per-locus mutation rates are low and recombination at this scale should play a minor role. The haploid population is fixed for a single genotype most of the time. Mutations arise rarely at rate *u* per bp per individual per generation and either rapidly die out or fix according to Kimura’s fixation probability (SI Appendix Section 1C) in a population of *N* individuals, with selection coefficients computed from Eq (3). Evolutionary dynamics for the entire population under the resulting fixed states approximation is then a Markov chain, which can be solved using linear algebra when the state spaces are small (e.g., for individual binding sites [10, 24]), or simulated efficiently using variants of the stochastic simulation algorithm [39], which is the route we will take.

## Results

### Evolution of CREs in the sharp threshold, low concentration (STLC) limit

We start with a theoretically interesting limiting case where the activation nonlinearity is a sharp threshold (i.e., a step) function at *µ*, which occurs when *δ* → ∞. We simultaneously unfold the binding nonlinearity by selecting *c*_max_ ≪ 1, so that no BS – even if it matches the TF motif exactly – can be close to saturated occupancy. In this STLC limit, *δ* is no longer a free parameter, *c*_max_ can be absorbed into a choice of units, and the threshold *µ* becomes the focal regulatory parameter shaping the replicated GP map and governing the evolution of CREs on it (SI Appendix Section 1B and 3A).

In the STLC limit, stochastic simulations of CRE evolution for a regulatory architecture typical of eukaryotes (CRE length *L* = 256bp, binding site length *ℓ* = 8bp, *ε* = 3; SI Appendix Section 1D) show a large variability in fitness trajectories at a fixed value of *µ*, consistent with previous reports [12, 24]. Consequently, we plot and quantify the evolutionary rates for emergence of function (“adaptation rates”) as averages over replicate simulation runs on a logarithmic time axis. Despite this variability, we observe an intriguingly strong non-monotonic dependence of the adaptation rate on *µ* (Fig. 2A), where intermediate values of *µ* lead to the fastest emergence of functional CREs. The choice of *µ* also strongly influences the structure of the evolved CREs. High *µ* values necessitate multiple close-to-consensus (with *k* = 0 or 1 mismatches) binding sites, evidenced by their strong over-representation in evolved sequences relative to the random background (Fig. 2B, green), as summarized in Fig. 2C. In contrast, low *µ* values trigger gene expression too readily, so much so that even random initial sequences may express irrespective of their cognate activating factors. This is heavily penalized by our fitness function, requiring selection to eliminate noncognate binding sites in the CRE, which results in their significant under-representation (Fig. 2B, violet).

**Figure 2.**
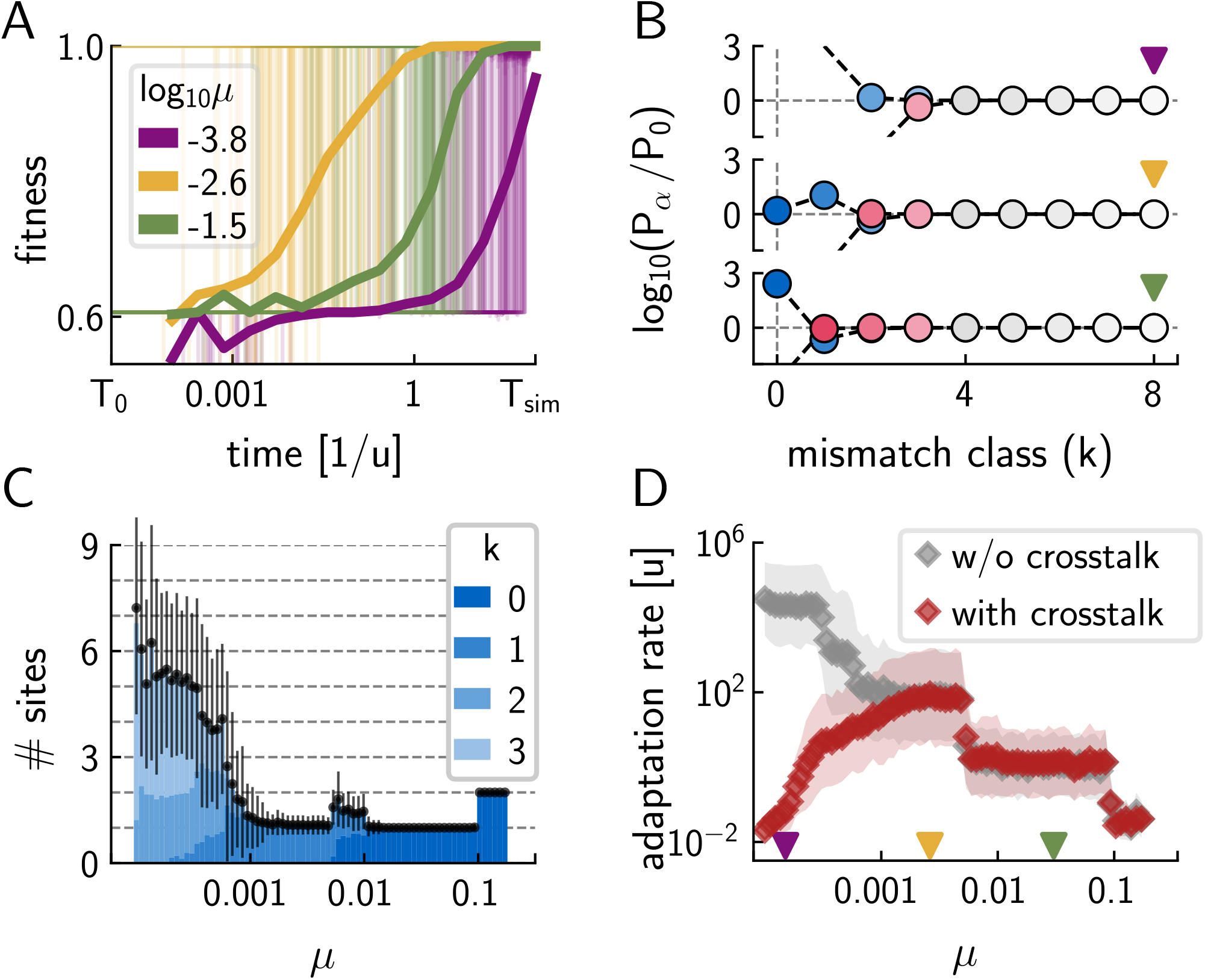
CRE evolutionary outcomes in the sharp-threshold low-concentration (STLC) limit. **(A)** Replicate simulation fitness trajectories (thin lines, 100 replicates per *µ*) and their averages (thick lines) at three activation threshold values *µ* (legend). Adaptation is fastest at intermediate *µ* (yellow). Simulations terminate at 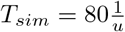. **(B)** Ratio between adapted (*P*_*α*_) and neutral (*P*_0_) mismatch distribution across the entire CRE for different *µ* (triangle color as in A), for cognate (blue) and noncognate (red) TFs. Positive (negative) values indicate enrichment (depletion) of the corresponding BSs in evolved CREs. Missing markers at small *k* indicate that no corresponding CRE sequence evolved across 1000 simulated replicates. Statistics are extracted when a replicate simulation run first exceeds fitness *F >* 0.99. **(C)** BS composition of evolved CREs as a function of *µ*. Color (legend) indicates the average mismatch composition of BS that together contribute at least 90% of the total cognate regulatory input; shown are average ± SD over replicates. At high *µ*, 1-2 BSs fully matching the cognate TF motif contribute; at low *µ*, the gene is activated by many weakly-contributing BSs. **(D)** CRE adaptation rate as a function of *µ*. Adaptation rate is defined by Eq. (4) (red markers and shade = average ± SD over 1000 replicate simulations; gray = same for the control scenario without crosstalk, *n* = 0). Colored triangles = *µ* values used in A and B. Parameters: *L* = 256, *ℓ* = 8, *n* = 3, *R* = 2, *c*_max_ = 0.1, *ε* = 3, *α* = 1, *N* = 100.

Between the two extremes, at intermediate *µ* values (Fig. 2B, yellow), evolving a single *ℓ* = 8bp locus into a close-to-consensus cognate binding site is sufficient to controllably activate expression (Fig. 2C) and CRE adaptation proceeds most rapidly (SI Appendix Section 3). Fig. 2D systematically explores the dependence of the adaptation rate *a* (in units of mutation rate *u*) on *µ*, to confirm the existence of an optimal *µ*^*\**^ value:

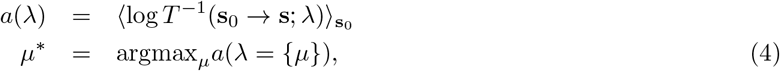

where *T* is the “completion time” taken to evolve from a random starting sequence **s**_0_ to a functional CRE **s** whose fitness first exceeds a high threshold value (here *F* = 0.99), and ⟨·⟩ indicates averaging over starting sequences **s**_0_ and multiple stochastic realizations of the evolutionary process.

Numerical optimization over explicitly simulated evolutionary trajectories, encapsulated by Eq (4) and depicted in Fig. 2D, is the first instantiation of our “optimize-to-adapt” approach. In the STLC limit, we reduce this to the optimization over a single component (*µ*) of the regulatory parameter vector (*λ*): at the optimal value *µ*^*\**^, functional CREs emerge up to ∼ 4 orders of magnitude faster than at alternative values – even though high fitness solutions do exist at many of those alternative values and could, in principle, be reached and maintained at the mutation-selection-drift balance.

The existence of an optimum in *µ* is non-trivial. Why does this optimum emerge only in the presence of non-cognate TFs (Fig. 2D, red vs gray; see SI Fig. S2 for a detailed analysis of the control case)? Intuitively, in the STLC limit, the fitness landscape is a piecewise constant function of gene expression and the governing evolutionary force is drift. On the one hand, at low *µ*, drift is very ineffective in suppressing erroneous “crosstalk” expression due to noncognate TF binding. One may be tempted to disregard this crosstalk entirely, which would trivially solve the problem for an isolated CRE. Yet the isolated CRE that we simulate is only one representative CRE on a replicated GP map for gene regulation that shares its regulatory machinery with all the other CREs in an organism [22]: and for those CREs, the roles of cognate and non-cognate TFs will be exchanged. Thus, crosstalk must be self-consistently included in calculations for any representative CRE [27, 28]; when included, it emerges as a major evolutionary constraint at low *µ*. This is consistent with the theoretical study reported in the companion paper [22], which demonstrates that the structure of optimal GP maps is strongly determined by common, not rare, phenotypes at regulatory loci, e.g., the prevalent selection “against non-cognate TF binding.” On the other hand, at high *µ*, drift is similarly ineffective in creating multiple close-to-consensus sites for cognate TFs which are required for gene activation. Here, drift amounts to a random evolutionary search in the absence of a selection gradient, leading to a slow, unguided exploration over an exponentially large genotypic space [24].

The tradeoff between these two extremes gives rise to an intermediate *µ*^*\**^ that maximizes the CRE adaptation rates. In the STLC limit of our setup, this regime typically yields one single strong cognate TF binding site in an evolved CRE (SI Appendix Section 3B). In the next section, we use information-theoretic arguments to make this reasoning precise; to show how to predict optimal *µ*^*\**^ without resorting to stochastic evolutionary simulations; and, as a result, to explore CRE adaptation rates in different regulatory architectures (by varying *ε, ℓ, L*, number of crosstalking TFs *n*, etc.) that were hitherto held fixed. These foundations will subsequently allow us to go beyond the STLC limit and consider more realistic regimes of CRE evolution.

### Information theory predicts when functional CREs emerge most rapidly in the STLC limit

Our simulation results can be rationalized using an information-theoretic approach for characterizing the cost of selection [22, 34]. We start by considering genotypes **s** in a sequence space *S* of size |*S*|, distributed according to some neutral distribution *φ*(**s**). Selection acts to enrich the sequences with higher fitness, which can be captured by a different distribution *ψ*(**s**). We can then speak of “genetic information” that selection has accumulated in the genomes, defined as a Kullback-Leibler divergence, in bits, between the neutral and selected distributions:

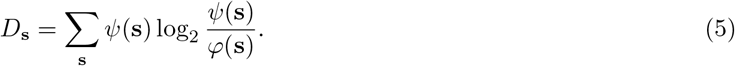

This quantity is very general but difficult to compute or estimate, because it entails manipulating distributions defined over very high-dimensional sequence spaces. Nevertheless, we can conveniently lower-bound it by “phenotypic information” [22, 34], which can be further simplified using additional assumptions into a form introduced previously by Wagner [40]. We will start with this simple and intuitive formulation, and return to the more general framework later.

We first assume that the neutral distribution is uniform over the sequence space, which would be expected if sequences were generated by point mutations with no mutational bias; this assumption is true for models that we explore here. The entropy of the neutral distribution is then *H*(*S*) = log_2_(|*S*|) bits. Let selection act on these random sequences by maintaining only the functional ones, **s**_*ρ*_ ∈ *S*_*ρ*_, whose phenotypes *ρ* satisfy some predefined criteria for function (e.g., high fitness), and discarding the sequences that do not match this criteria. Specifically, selection cleanly partitions all sequences into two disjoint categories, functional and non-functional, yet does not distinguish between sequences in each category. Then we say that the evolutionary process has accumulated “phenotypic information” of *𝒟* bits:

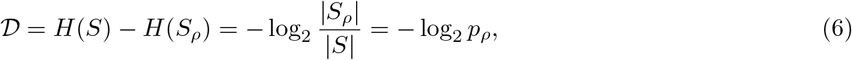

where *p*_*ρ*_ = |*S*_*ρ*_| */* |*S*| is a function of the GP map that determines the fraction of functional sequences in the genotype space *S*. This simple counting approach, which compares the number of functional to the number of all genotypes, is a special case of a more general framework based on Eq. (5) that holds for non-uniform neutral distributions, smooth or probabilistic genotype-phenotype-fitness maps, and various criteria for what is “functional” (specified in terms of population states, genotypes, multi-dimensional phenotypes, average fitness, etc.) [22, 34]. In the STLC limit, however, information *𝒟* defined either via the simple Eq. (6) or within the more powerful and generic IT framework, yields identical results, and lower bounds the genetic information, such that *𝒟* ≤ *𝒟*_**s**_ (SI Appendix Section 2A,B).

Applied to the case of regulatory sequences at hand, whether or not a CRE **s** is functional depends entirely on the its 2D phenotype *ρ* = (*ρ*_ON_, *ρ*_OFF_): i.e., on the total regulatory input in the ON vs the OFF environment. While this phenotype is trivially computable for any given sequence **s** and regulatory parameters *λ*, an average over various functions of this phenotype over the entire space *S* is not trivial. Nevertheless, *p*_*ρ*_ and thus can be estimated via Monte Carlo sampling: we randomly pick ∼ 10^7^ sequences of length *L* = 256bp and project them onto the 2D phenotype plane (Fig. 3A,B; SI Appendix Section 3A). Here, “being functional,” and thus contributing positively to *p*_*ρ*_, implies that a sequence resides in the lower right corner of the phenotype plane, where the total regulatory input is above (resp., below) the activation threshold *µ* in the ON (resp., OFF) environment. As a function of *µ*, the curve *𝒟* (*µ*) exhibits a distinct minimum at 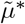 where functional sequences are statistically “minimally displaced” from random ones (SI Appendix Section 2A,B; we use the ∼symbol to denote that this threshold value is obtained via information-theoretic optimality). According to our theory [22], selection for regulatory function on the replicated GP map for CREs will automatically favor such IT-optimal 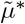 in evolutionary equilibrium, no matter how this regulatory parameter is encoded in the genome. According to our hypothesis, this is also the regime where the CRE sequences will evolve fastest (Fig. 3A).

**Figure 3.**
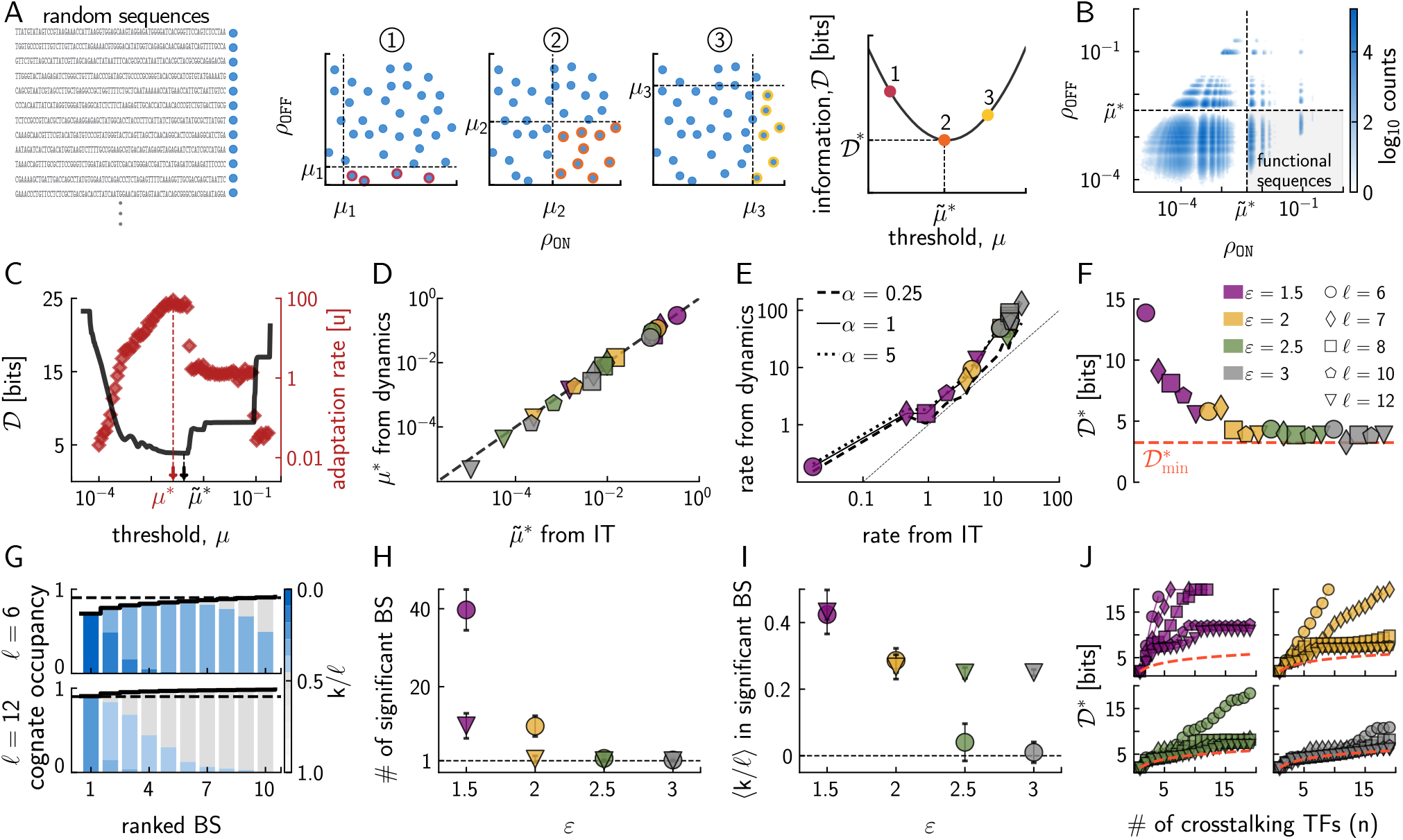
Information theory predicts optimal regulatory parameters in the STLC limit. **(A)** A schematic of the information-theoretic argument in the STLC limit. Random sequences (left) map into 2D phenotypes *ρ* = (*ρ*_ON_, *ρ*_OFF_) (see text) depicted on a plane (each blue dot = one sequence). For any given activation threshold *µ* (dashed lines; circled numerals 1,2,3 depict three example values), functional sequences (outlined dots) lie in the lower right corner of the plane (middle), and their fraction defines information *𝒟* via Eq. (6). IT-optimal activation threshold 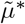 minimizes *𝒟* (right). **(B)** Monte Carlo sampling of random sequences depicted in the (*ρ*_ON_, *ρ*_OFF_) plane (SI Appendix Section 3A). **(C)** Overlay of 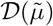 (black, left axis, Monte Carlo over random sequences) and adaptation rate estimated via “optimize-to-adapt” approach (red, right axis, evolutionary dynamics simulations). Activation threshold 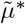 that minimizes *𝒟* (black arrow) nearly coincides with activation threshold *µ*^*\**^ that maximizes adaptation rate (red arrow). **(D)** Activation thresholds from “optimize-to-adapt” simulations match IT-optimal thresholds that minimize *𝒟* across different mismatch energy *ε* and binding site length *ℓ* values (legend in F). **(E)** Adaptation rates from “optimize-to-adapt” simulations recapitulate rates derived (see text) at IT-optimal activation thresholds (legend in F), independently of population size and selection strength *α* (legend). **(F)** As *ε* and *ℓ* increase, optimal information D*\** approaches the IT bound (dashed red, Eq. (7)). **(G)** Fractional contributions to cognate regulatory input, *ρ*_*s*_, at IT-optimal activation threshold, for short (*ℓ* = 6bp, top) or long (*ℓ* = 12bp, bottom) binding sites. Evolved BSs are ranked by contribution (x-axis), color (bar at right) shows the normalized match to TF consensus (dark blue = full match, gray = complete mismatch), thick black line = cumulative input, dashed black = ≥ 90% threshold. Multiple sites contribute at *ℓ* = 6 in contrast to *ℓ* = 12. **(H)** Number of significant BSs that contribute 90% of cognate regulatory input, shown as a function of mismatch energy (x-axis, color), for short (*ℓ* = 6, circles) and long (*ℓ* = 12, triangles) binding sites. At low *ε* irrespective of *ℓ*, significant contribution comes from multiple BSs; at high *ε*, evolved CREs harbor only one significant BS. **(I)** Fraction of mismatches in significant BSs, plotting conventions as in I. At low *ε*, multiple BS with mismatches contribute significantly to cognate regulatory input; at high *ε*, evolved CREs harbor only one significant BS perfectly matching TF consensus for *ℓ* = 6 while several mismatches are permitted for *ℓ* = 12. **(J)** As the number of noncognate TFs *n* (x-axis) increases, IT-efficient regulatory architectures utilizing optimal threshold *µ*^*\**^, which match the 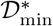 bound (dashed red line), exist only if binding sites are sufficiently long (plot symbols for *ℓ* as in F) and mismatch energy is sufficiently high (color as in F). Otherwise, evolved CREs contain multiple significant BSs and cannot be IT-efficient due to overspecification. Baseline parameters: *L* = 256, *n* = 3, *R* = 2, *c*_*max*_ = 0.1, *N* = 100, *α* = 1; in B-C *ℓ* = 8, *ε* = 3; in G *ε* = 2. Statistics in G,J are averages and in H-I average ± SD over 100 replicate simulations. Reported adaptation rates in C, and E are averages over 1000 replicates (same as in Fig. 2).

In Fig. 3B-E we verify this hypothesis and its match to IT predictions using a large-scale simulation study. For a representative pick of the mismatch energy *ε* = 3 and binding site length *ℓ* = 8, the IT-optimal value 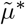 is very close to the value *µ*^*\**^ that maximizes adaptation rate of Eq. (4), obtained by the “optimize-to-adapt” approach. Fig. 3D shows that this correspondence is tight and robust across a range of values for *ε* and *ℓ*. When evolution is mostly neutral (as in the STLC limit) and thus well-described by a random search in sequence space, adaptation rate is ≈ 2^*−𝒟*^*Lu*, i.e., the product of *p*_*ρ*_ = 2^*−𝒟*^, the fraction of functional genotypes; the CRE length *L*; and the mutation rate *u*, independently of the population size *N* and selection strength *α*. This simple approximation captures the adaptation rate surprisingly well for an extended range of regulatory architectures (Fig. 3E, SI Appendix Section 3C, SI Fig. S3).

For our setup, information theory puts a lower bound on information *𝒟* as a function of the number of non-cognate factors *n*:

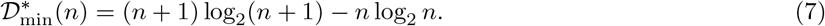

This combinatorial bound does not rely on any postulated mechanism for TF-DNA sequence recognition and is thus surprisingly general (SI Appendix Section 2B); regulatory architectures that reach it are “IT-efficient” and therefore the best possible, in an absolute sense [22]. For example, with *n* = 3 noncognate TFs, and IT-optimal activation threshold in the STLC limit, Fig. 3F shows that IT-efficient architectures exist for *ℓ >* 6 and *ε* ≥ 2.5. In these architectures, a single binding site in the CRE for the cognate TF contributes almost the entire required regulatory input *ρ*_*s*_, consistent with Fig. 2. If *ℓ* and *ε* increase even further at constant *n* (Fig. 3G,H), entropy considerations dictate that this single binding site develops mismatches [32, 41]. In contrast, for *ℓ* and/or *ε* that are too low, multiple sites for the cognate TF must evolve to turn the gene on, even using IT-optimal activation threshold (Fig. 3I). Among these sites, some are pushed strongly towards the consensus while others harbor more mismatches (Fig. 3G,H) – yet despite this flexibility, the regulatory architecture is not IT-efficient, as it does not saturate the 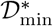 bound. This is our first example of “overspecification” (a term introduced already by Berg and von Hippel [1] and recently given an information-theoretic interpretation [22]): the inability of biological mechanisms to fully implement IT-efficient regulation, which in turn necessitates the selection to accumulate extra genetic information in the CREs.

Fig. 3J summarizes how the number of noncognate TFs, *n*, controls the transition between IT-efficient and inefficient regimes, and the transition’s dependence on the regulatory architecture (explored by changing *ε* and *ℓ*). While it is difficult to estimate the biologically relevant value for *n*, a quick divergence of *𝒟*^*\**^from the IT-efficiency bound even for small *n* ∼ 10 seen in Fig. 3J suggests typical values for the mismatch energy and binding site length should be at least *ε* ≥ 2.5 and *ℓ* ≥ 6, as observed empirically.

In sum, fundamental theory as well as simulation-based “optimize-to-adapt” approach consistently predict a biophysically plausible, IT-efficient, and most adaptable regime, where a single cognate binding site within a CRE can appropriately activate a desired gene in the STLC limit with optimally-chosen activation threshold *µ*. In this limit, it is always beneficial to utilize larger *ℓ*, i.e., to evolve towards prokaryotic-like architectures that favor one sufficiently long binding site (perhaps even long enough to harbor mismatches). This is in contrast with extensive evidence that eukaryotic CREs contain multiple short BSs of diverse specificity. In the next section, we progressively relax the STLC limit and examine how the corresponding optimal predictions change.

### Beyond the STLC limit, CREs are enriched in multiple weaker binding sites

We first relax the low-concentration assumption while keeping the activation threshold sharp (*δ* → ∞). Following the Monte Carlo steps used to estimate *𝒟* as a function of the single regulatory parameter *µ* in Fig. 3C, we now analogously estimate *𝒟* as a two-dimensional function of a pair of regulatory parameters: the activation threshold *µ* and the maximal TF concentration *c*_max_. Fig. 4A shows a trough of IT-optimal solutions minimizing *𝒟* in the (*µ, c*_max_) plane that gets progressively deeper as both parameters are jointly taken towards zero; “optimize-to-adapt” simulations in Fig. 4B (left) mirror these IT results almost precisely. Thus, even if high TF concentrations are permitted and binding sites could be fully occupied, we nevertheless recover the STLC limit as optimal, with IT-efficient architectures still favoring a single strong cognate BS.

**Figure 4.**
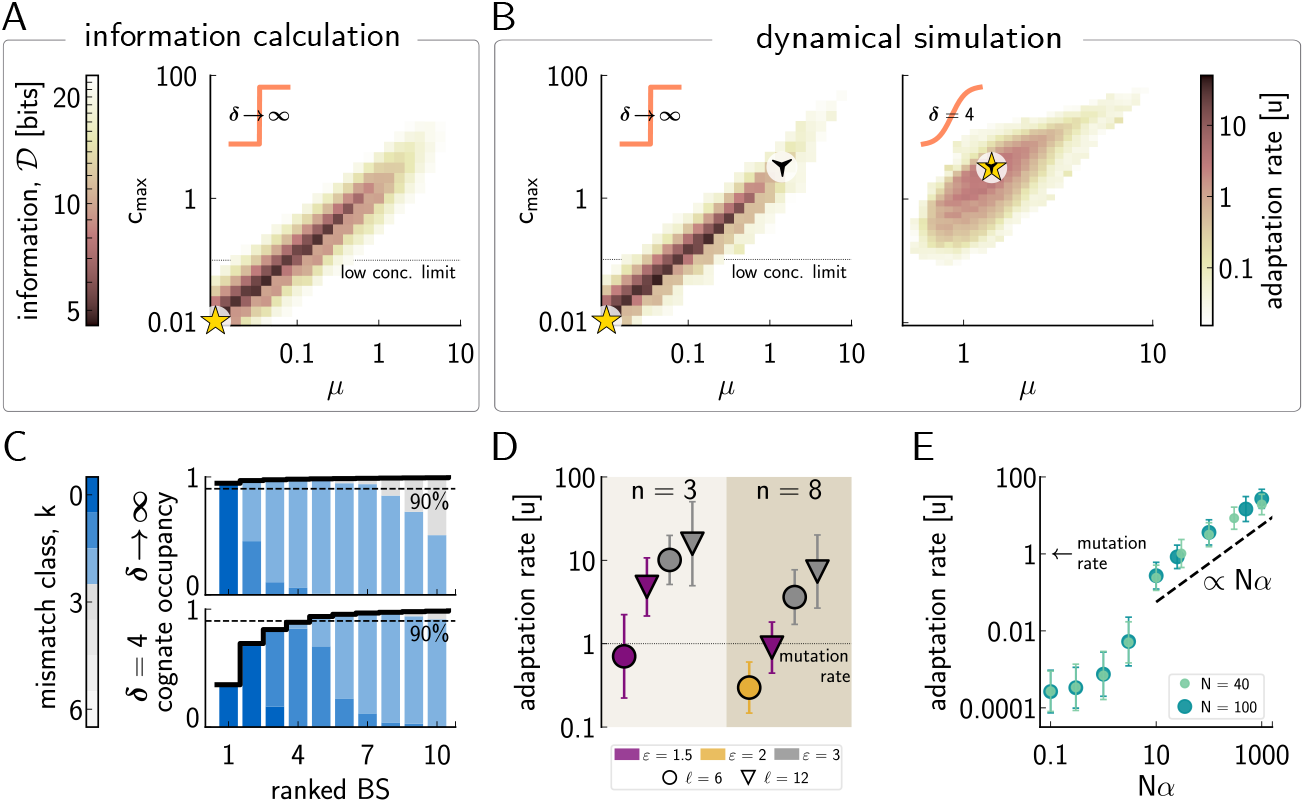
CRE evolutionary outcomes beyond the sharp-threshold low-concentration (STLC) limit. **(A)** Information (colorbar) in the sharp activation threshold (*δ*→ ∞) limit, as a function of regulatory parameters *µ* and *c*_max_, estimated using Monte Carlo sampling over 10^7^ random sequences. Dotted line = low concentration limit of Fig. 2; star = IT-optimal parameters that minimize *𝒟*. **(B)** CRE adaptation rate (colorbar) in “optimize-to-adapt” dynamical simulations, for sharp (left) vs smooth (right, *δ* = 4) activation nonlinearity. Stars = parameters that maximize adaptation rate. A non-degenerate optimum appears for the smooth activation nonlinearity (right) at *µ*^*\**^ = 1.407,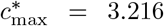. **(C)** Fractional contributions to cognate regulatory input, *ρ*_*s*_, at optimal regulatory parameters (stars in B) with a sharp (top) and smooth (bottom) activation nonlinearity. In the latter case, a larger number of BSs with broader range of specificity contribute to activate the gene. Plotting conventions as in Fig. 3G. **(D)** Adaptation rate for different regulatory architectures (binding site lengths *ℓ*, mismatch energy *ε*, and number of noncognate TFs *n*; see legend). Regulatory parameters (*µ, c*_max_) are separately optimized for each case. **(E)** Adaptation rate at optimal regulatory parameters, as a function of population size *N* (legend) and selection strength *α*. For low *Nα*, drift dominates; for *Nα* ≫ 1, selection dominates in the scaling regime (dashed line). Baseline parameters: *L* = 256, *ℓ* = 6, *n* = 8, *R* = 2, *ε* = 3, *α* = 1. Statistics are average ± SD over 100 replicate simulations.

These conclusions change dramatically if *δ* is finite and the activation nonlinearity is therefore not a sharp threshold but rather a smooth sigmoid function, as would be expected for biologically realistic systems. The results of “optimize-to-adapt” dynamical simulations at *δ* = 4 are shown in Fig. 4b (right). In absence of unambiguous empirical data, we chose *δ* = 4 arbitrarily, but insisted it be of order of unity to provide a strong contrast with the sharp threshold (see SI Appendix Section 3 for a detailed analysis covering different choices for *δ*). At finite *δ*, CRE adaptation is fastest at a single non-degenerate optimal combination (*µ*^*\**^,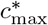). Whereas in the STLC limit an individual consensus BS contributed nearly all cognate regulatory input (Fig. 3G), at *δ* = 4 evolution with optimal regulatory parameters favors a broader spectrum of BSs within each CRE. For example, in Fig. 4C, four 6bp binding sites emerge to activate the gene: two, with zero mismatches, reach 50% occupancy at optimal concentration and contribute ∼ 77% of regulatory input, and two, with one mismatch each, that contribute a large share of what remains. This diversity of contributions is larger for longer binding sites (SI Figs. S4, S5).

Despite the lifting of degeneracy in optimal regulatory parameters (Fig. 4B) and clear differences in evolved CREs (Fig. 4C), Fig. 4D shows that the dependence of CRE adaptation rate on mismatch energy *ε*, binding site length *ℓ*, and the number of noncognate TFs *n*, remains qualitatively unchanged from the STLC case. Adaptation is slower with more crosstalk (higher *n*), and it is always accelerated by using longer BSs that can be more specific. In other words, while going beyond the STLC limit explains the increase in the diversity and multiplicity of evolved BSs in functional CREs for fixed *ℓ*, it still does not explain if and why shorter binding sites should be evolutionarily favored compared to longer sites. We will return to this question at the end of the Results section.

Beyond the STLC limit, adaptation rate is no longer independent of the selection strength set by *α* (Fig. 4E). For small population size *N* or small *α*, the governing evolutionary force indeed remains drift, but for sufficiently high *Nα* ≫ 1 we enter the “scaling regime” where adaptation rate is proportional to *Nα* (SI Appendix Section 4A,B). Here, the smooth activation nonlinearity permits the population to respond to the selection gradient rather than perform a random search through the sequence space as before, which should speed up the emergence of fit CREs. Yet, this is not what we observe – the adaptation rate is slower (not faster) with a smooth nonlinearity, even at optimal regulatory parameters (cf. adaptation rates at yellow stars in Fig. 4B)!

To explain this conundrum, we turn to the powerful new information-theoretic framework for the cost of selection [22, 34]. In this framework, various information measures *D*_*X*_ are defined as Kullback-Leibler divergences between selected-for vs neutral distributions for *X*, where *X* can stand for population states, genotypes, phenotypes, or functions thereof. For certain *X*, these measures can be tractably estimated or bounded.

Here we focus on two such measures, *D*_mm_ and *D*_fit_. The first, *D*_mm_, measures the “work” that selection has exerted to displace stationary mismatch distributions for cognate and non-cognate TFs away from neutral (random sequence) expectation in order to make CREs have high fitness (Fig. 5A), quantifying in IT terms the enrichment (or depletion) of BSs in evolved CREs reported in Fig. 2B. A more principled measure would have been *D*_**s**_ introduced in Eq. (5), the divergence of entire *L* = 256 bp evolved sequence distribution from the neutral one; unfortunately, due to the curse of dimensionality, these distributions can be estimated neither empirically nor in simulations. Nevertheless, *D*_**s**_ can typically be bounded from below as *D*_mm_ ≤ *D*_**s**_ (SI Appendix Sections 2C,D), since mismatch counts are statistics of the entire sequence distribution. The gap between the two information measures is due to the neglect of site-to-site correlations in *D*_mm_ (which arise because evolving one BS releases pressure on other sites, as well as from convolutional overlaps of between various BSs); *D*_mm_ treats all *ℓ* bp windows within a CRE of length *L* as independent, whereas in fact they are not. On the other hand, this measure can easily be estimated from CREs (actual or simulated) without any knowledge of the GP map beyond the TF motifs or energy matrices. Also, it is a straightforward generalization of the “sequence logo” information [42] and related estimates [25, 43], while taking noncognate binding into account.

**Figure 5.**
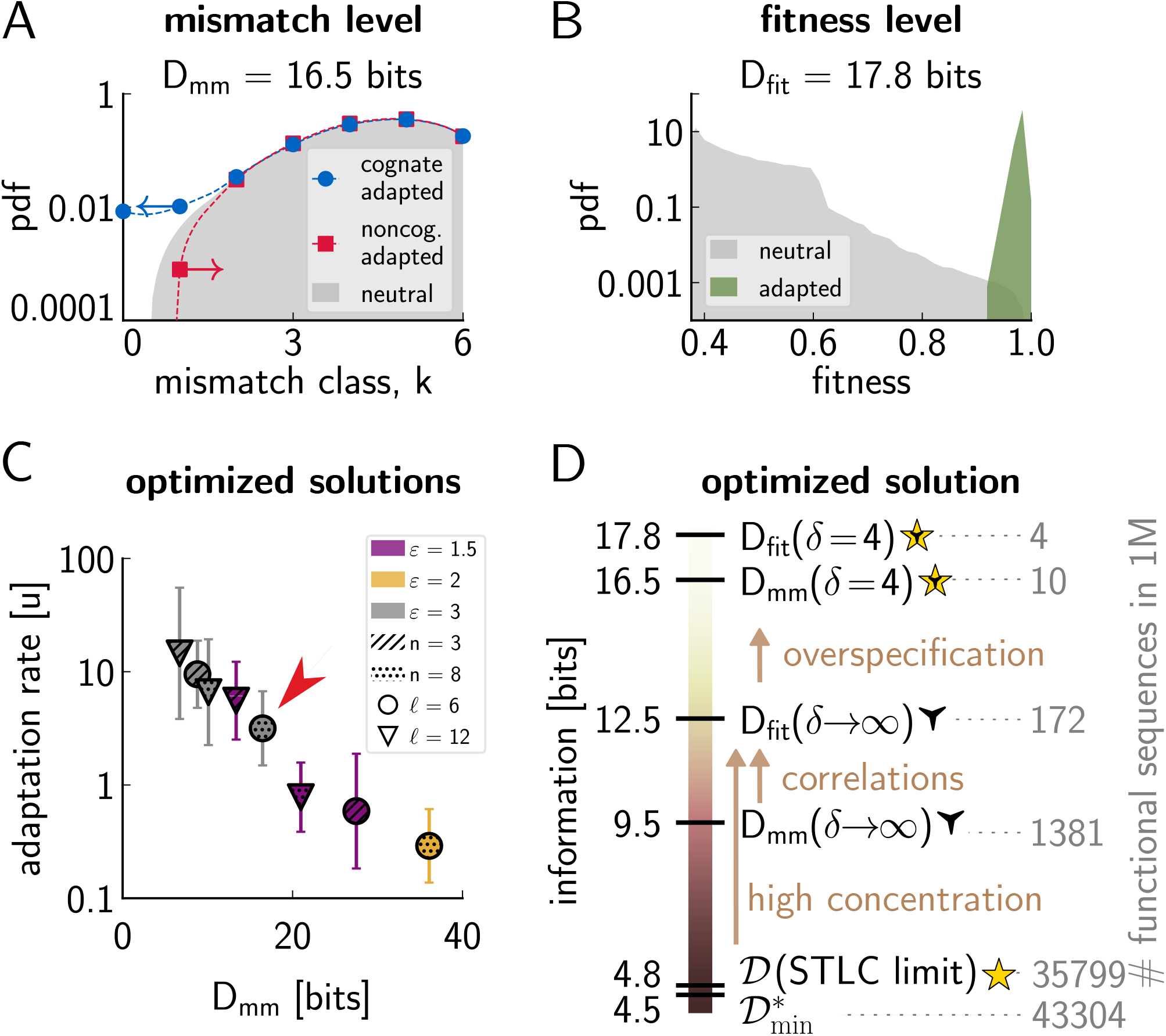
Information-theoretic analysis of CRE adaptation beyond the STLC limit. **(A)** Adapted cognate (blue) and noncognate (red) mismatch distributions for functional CREs. For random sequences, the mismatch distribution is binomial (gray). *D*_mm_ quantifies the divergence, in bits, of adapted distributions from the neutral expectation, here amounting to 16.5 bits for *n* = 8 noncognate TFs, *L* = 256 bp, *ℓ* = 6 bp, and *ε* = 3, smooth activation nonlinearity with *δ* = 4, and an optimal choice of *µ* and *c*_max_. **(B)** Adapted (green) and neutral (gray) fitness distributions. Neutral distributon is estimated over 10^8^ random sequences; parameters as in (A). **(C)** CRE adaptation rate decreases exponentially with mismatch information across a range of regulatory parameters (legend). **(D)** Mismatch and fitness information, *D*_mm_ and *D*_fit_ (left vertical axis), respectively, accumulated by selection in functional CREs, for smooth (*δ* = 4) and sharp (*δ* → ∞) activation thresholds at regulatory parameters (*µ, c*_max_) (indicated by markers) as in Fig. 4B. Regulatory parameters are as in (A-B), and denoted by a red arrow in (C). Numbers in gray (right vertical axis) interpret the corresponding information measures as the equivalent number of functional CREs in a pool of 10^6^ random sequences.

In addition to *D*_mm_, which lower-bounds genotype information *D*_**s**_, we can also directly estimate *D*_fit_, the information that selection has accumulated on the fitness level, by comparing the distributions of *F* given by Eq. (3) over random vs evolved CREs (Fig. 5B). Since sequences fully determine fitness in our model, theory tells us that *D*_fit_ ≤ *D*_**s**_ [34]. For the example case shown in Fig. 5A and B, comparison between *D*_mm_ = 16.5 bits and *D*_fit_ = 17.8 bits suggests that the mismatch information is a reasonable approximation for the information on the fitness level; extensive simulations confirm this across a wide range of parameters (SI Fig. S6). Moreover, *D*_fit_ generalizes D introduced in Eq. (6) – in the STLC limit, the two measures are identical (SI Appendix Section 2A), but *D*_fit_ remains well-defined even when genotypes cannot be uniquely partitioned into two disjoint sets of functional and non-functional CREs.

The relationship between the two information measures is both interesting and important: *D*_mm_ reports a displacement that selection has generated at the level of BS mismatches in CRE sequences, while *D*_fit_ tells us that selection exerted similar work to separate functional from non-functional sequences at the fitness level. By definition, adapted CREs are those that exceed a high threshold value of *F*, so their “adapted” fitness distributions (Fig. 5B, green) must be very stereotyped; hence, *D*_fit_ depends on the regulatory parameters primarily via the neutral fitness distribution (Fig. 5B, gray). Perhaps surprisingly, it is this neutral distribution that carries a real imprint of the genotype-phenotype map: evolvable maps are those that maximize the fraction of functional (high fitness) random sequences and thus maximize the overlap (equivalently, minimize *D*_fit_) between neutral and adapted fitness distributions [22]. In contrast, at the mismatch level, the neutral distribution (Fig. 5A, gray) is always trivial for our toy model – it is simply a binomial distribution, Bin(*ℓ, p* = 3*/*4). Selection reshapes its tails (Fig. 5A, color), and evolvable GP maps correspond to minimal displacements in those tails required to achieve CREs with high fitness.

Mismatch information predicts CRE adaptation rate very well (Fig. 5C, SI Fig. S7), as does *D*_fit_ (SI Fig. S6), extending the results from the STLC limit. For architectures with optimized regulatory parameters (*µ*^*\**^,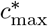), linear increases in sequence information that selection must accumulate lead to exponential slowdowns in the CRE adaptation rate, and this relationship is steeper at lower effective selection strength (SI Fig. S8); yet in any case, more information in total must be accumulated with a smooth nonlinearity to reach high fitness. Detailed information analysis for a particular choice of *ε* and *ℓ* in Fig. 5D (and more generally, see SI Fig. S6) again points to “overspecification” as the root cause – this is our second example where, due to biophysical constraints (i.e., a limit to activation nonlinearity slope), selection must accumulate extra information to ensure proper function. Here, to ensure high fitness with a smooth activation nonlinearity, cognate regulatory input must not only be slightly above the sharp threshold (as in STLC limit), but substantially so, requiring higher number of significant cognate binding sites (and vice versa for selection against noncognate binding).

Taken together, a biophysically-realistic smooth activation nonlinearity affects CRE adaptation rates in two opposing ways. On the one hand, it speeds up adaptation by providing non-zero selection gradient even when the sequence is far from being fully functional – this effect is particular to the nature of evolutionary dynamics and is not captured directly by our IT calculations that apply in evolutionary steady state [22]. On the other hand, smooth activation nonlinearity slows adaptation down by requiring more information to be accumulated and maintained at steady state due to overspecification – this effect is generic and pertinent regardless of which adaptation dynamics (or learning rule) would be used. How these two effects balance out at a given selection strength determines the optimal regulatory architecture and thus the most evolvable GP map for gene regulation (SI Fig. S8). More generally, an intuitive way to think about these constraints, and about the cost of selection to evolve function, is to interpret various information measures *D*_*X*_ as probabilities,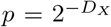, of drawing a functional sequence with high fitness by chance from a pool of entirely random sequences. For example, Fig. 5D suggests that in a pool of 1 million random sequences, ∼ 200 would be functional if the activation threshold were sharp, but this would drop by about 40-fold due to overspecification for a smooth nonlinearity.

### CREs harboring short, cooperatively-acting binding sites of diverse specificity are evolutionarily optimal

Finally, we return to the evolutionary origins of eukaryotic regulatory architectures that utilize short (*ℓ* ∼ 6 − 8 bp) binding sites. In Fig. 4 we argued that if maximal TF concentrations, *c*_max_, can be adjusted freely, then longer *ℓ* are always preferred and would be, for certain combinations of parameters *L, n*, and *ε*, also IT-efficient in the STLC limit (Fig. 3). If maximal TF concentrations were instead limited but activation nonlinearity remained smooth, then longer BSs would be hampered instead of benefiting entropically, compared to short BSs. To maintain sufficient occupancy and contribute significantly to regulatory input even at limited *c*_max_, short BSs would need to fix less mutations than long BSs. In other words, selection would need to accumulate less sequence information, making short BS architecture more evolvable. One avenue towards explaining short BSs would therefore invoke a regime of low crosstalk and a strong physical constraint on *c*_max_. While possible in principle, this regime seems implausible for eukaryotes in practice. Evolution has tuned intracellular TF concentrations to be higher in eukaryotes; if anything, longer BSs should be favored. We therefore wondered if other known regulatory mechanisms, hitherto not taken into account in our model, could significantly change the optimality predictions, even without explicit resource limits on *c*_max_.

Cooperative interactions in TF binding are one such promising avenue to explore. They enable efficient gene regulation at lower TF concentrations while also mitigating some of the detrimental effects of crosstalk due to noncognate TFs [27].

In prokaryotes, a broadly utilized mechanism is “homodimer cooperativity” [2], which we model as follows. We consider the evolutionary emergence of “compound BSs” of length *ℓ* bp that can bind two cognate TF molecules with a motif length of *ℓ/*2 each. If the joint occupancy of the two half-sites by the two TF molecules is energetically strongly favored (here, by a cooperative binding energy *E*_*c*_ *<* 0), we can replace regulatory contributions at location *i* for TF *j* in Eq. (1) with contributions from the two half-sites (SI Appendix Section 1B):

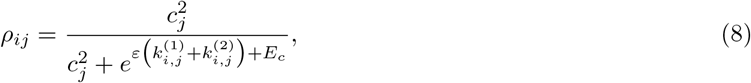

where 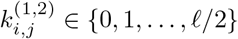 are the mismatches of the motif of *j*-th TF at location *i* in the CRE with the first (or second, respectively) half-site. The previously analyzed case of non-cooperative binding to a single BS of length *ℓ* provides a natural comparison to the homodimer cooperativity model with two half-sites.

Additionally, we also consider the “synergistic activation” implicated in eukaryotic gene regulation [44, 45]. While different molecular mechanisms could lead to the reported synergistic effect, they all share a common feature whereby the joint binding of multiple TFs contributes more than the sum of individual binding effects towards turning the gene on. Moreover, BSs are geometrically much less constrained in their placement or orientation within the CRE, compared to the prokaryotic case where the two half-sites typically need to maintain a rigid arrangement in close proximity.

The simplest model for such synergism introduces an extra parameter *η*≥ 1 that controls the degree of nonlinear summation of individual binding contributions to the total regulatory input (SI Appendix Section 1B):

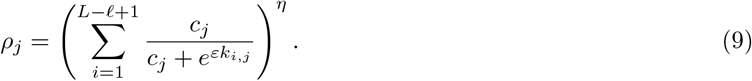

We performed “optimize-to-adapt” simulations of CRE evolution with homodimer cooperativity (*E*_*c*_ = − 4) and synergistic activation (*η* = 3), and compared them to the *ℓ* = 6 and *ℓ* = 12 noncooperative regulatory architectures in Fig. 6 (details and heterodimer cooperativity in SI Figs. S9, S10). As before, for each architecture the regulatory parameters (*µ, c*_max_) have been separately optimized, using a smooth activation nonlinearity (*δ* = 4) throughout. Fig. 6A reveals a surprising result of these simulations: with both models of cooperativity, the most evolvable architecture now utilizes shorter, 6bp-long BSs, instead of longer ones – even if evolution can tune maximal TF concentrations freely. Synergistic activation, in particular, delivers large speedups, likely because it relaxes co-localization constraints relevant for homodimer cooperativity, and is therefore entropicaly favored.

**Figure 6.**
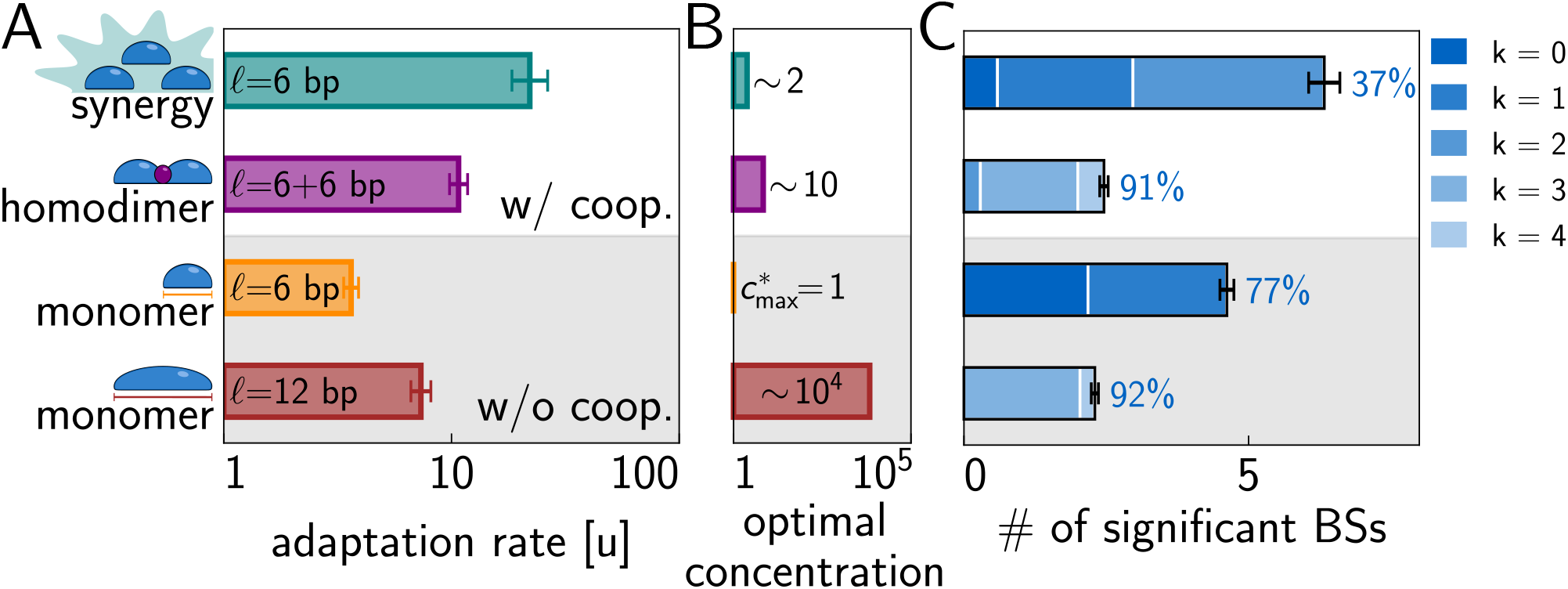
Functional CREs emerge fastest in a regulatory architecture with short, synergistically-acting binding sites of diverse specificity. **(A)** CRE adaptation rate in two regulatory architectures with cooperativity (top two rows: synergistic activation with *η* = 3, homodimer cooperativity with *E*_*c*_ = −4) and two architectures without (bottom rows: BSs of *ℓ* = 6 or 12bp in length), with regulatory parameters (*µ, c*_max_) optimized separately for each architecture. **(B)** Optimal maximal TF concentrations, 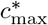 normalized to the reference case of *ℓ* = 6 without cooperativity. **(C)** Structure details of evolved CREs across different regulatory architectures. Bars show the number of BSs contributing significantly to total regulatory input, broken down by the number of mismatches (bluish hue; legend). Percentages show the fraction of total regulatory input accounted for by BSs with occupancy *>* 0.5 at optimally chosen *c*_max_. Synergistic activation utilizes multiple short binding sites of diverse specificity (due to a spectrum of mismatches) in each CRE, with many contributing BSs not close to saturated occupancy. Parameters: *L* = 256, *n* = 8, *R* = 2, *δ* = 4, *ε* = 3, *α* = 1. Statistics are average ± SD over 100 replicate simulations.

Cooperativity also enables resource-efficient evolutionary outcomes: the optimal TF concentration,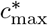, for synergistic activation is only two-fold higher compared to the optimum for baseline non-cooperative *ℓ* = 6 architecture, and is ten-fold higher for homodimer cooperativity (Fig. 6b). In contrast, for 12bp long BSs without cooperativity, optimal 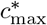 needs to be unreasonably high, as this architecture enjoys adaptation benefits only when BSs can reach high occupancy with many mismatches. This is consistent with previous work that demonstrated an exponential slow-down in the BS evolutionary emergence rates with increasing *ℓ*, in case where *c*_max_ cannot be simultaneously adjusted [24]. Lastly, Fig. 6C suggests that synergistic activation, much more so than homodimer cooperativity, favors architectures where multiple short sites of diverse specificity – e.g., harboring 0, 1, or 2 mismatches in 6bp long BSs – contribute jointly to activate a gene, and that a large fraction of these contributions comes from BSs that, at optimal 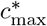, are occupied less than half of the time. These predictions are qualitatively consistent with known features of eukaryotic CREs that have, to date, remained unexplained [30].

## Discussion

We motivated this work by four unresolved puzzles about eukaryotic gene regulatory architecture. Central to these puzzles is the observation that in eukaryotes, regulation is effected by TFs binding to short binding sites of “diverse specificity,” i.e., featuring both strong and weak yet functional sites [30]; and that regulatory information appears to be distributed densely across entire CREs [13]. These observations seem counterintuitive and at odds with what a well-designed system might look like.

What should a well-designed system look like? The absolute benchmark must be an IT-efficient system, which reaches the 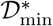 bound (for our example setup, this is given by Eq. (7)). No matter how information is stored in or read out from the sequence, Shannon’s theory excludes the existence of systems that could achieve the desired regulatory function with lower information [22]. While this bound can be approached in certain limits that make theoretical sense – e.g., in the STLC limit with long BSs and high mismatch energy – biologically realistic systems might operate far from that kind of efficiency. For instance, the physical constraints of smooth activation nonlinearity, and a limited *ϵ* that underpins molecular recognition, require substantial “overspecification” – i.e., an additional information-theoretic cost to selection – to evolve functional CREs. Larger overspecification, in bits, translates into the need to fix more alleles at regulatory loci, which results in slower CRE adaptation. Regulatory architectures that minimize this overspecification as much as possible within allowed constraints are IT-optimal (even if not IT-efficient), which should lead to more evolvable genotype-phenotype maps and thus to faster CRE adaptation. More precisely, we hypothesized that replicated genotype-phenotype maps whose regulatory parameters will have evolved towards their free fitness maximum (and thus information minimum) studied at evolutionary equilibrium in the companion paper [22], will be the same maps that are most evolvable in the dynamic sense, i.e., maps on which CREs adapt most rapidly.

Informed by these considerations, we used the “optimize-to-adapt” approach to compare CRE evolvability across regulatory architectures that feature BSs of different length, specificity, and TF cooperativity, each time adjusting the remaining regulatory parameters (activation thresholds, TF concentrations) to maximize the CRE adaptation rates. This computational process mimics the outcomes of long-term evolutionary adaptation which would similarly drive the regulatory parameters of replicated GP maps towards their optimal values, as we have shown in the companion paper [22]. The “optimize-to-adapt” simulation results robustly support our hypothesis.

Within a flexible model class able to capture some of the essential aspects of eukaryotic regulation, we find that CRE architectures featuring short binding sites of diverse specificity are most evolvable. This optimality robustly emerges in the presence of crosstalk, with realistic activation nonlinearities, and even when TF concentrations and gene activation thresholds can be adjusted to give every regulatory architecture its “best shot.” Importantly, cooperative (synergistic) action of multiple BSs turns out to be necessary to access this optimal regime of CRE sequence evolution. In other words, cooperativity of molecular interactions is a real game changer, not only for systems biology where its implications have been thoroughly investigated in the past, but also for evolution of gene regulation, where its central role has, to date, not been fully appreciated.

The presence of weaker binding sites appearing side-by-side with strong ones in a CRE densely packed with regulatory information is thus not necessarily a signature of limited selection strength, of adaptation transients, or of selection for robustness; nor of the intrinsic “noisiness” of biological pathways or a mechanism for fine-tuning of gene expression. Instead, this architecture emerges simply as a steady-state evolutionary outcome in a molecular-recognition-based regulatory system with sufficiently rich nonlinear structure. Such architectures have regulatory parameters that coevolve together with CREs so that, over the long run, selection will tend to maximize the number of degenerate high-fitness solutions. This route to evolvability emphasizes the positive role for an explosion of high fitness peaks or ridges, much as deep learning relies on a similar structure of loss landscapes in the overparameterized regime [46]. While other postulated benefits of the observed architecture are entirely plausible, they are not necessary for the minimal explanation. Allowing for some speculation, one could view our application of evolutionary and information theory as an *ab initio* prediction (or perhaps a post-diction?) for eukaryotic regulatory architecture, which emerges as soon as the molecular machinery allows for sufficiently flexible and complex nonlinear sequence readout. As we argue in the companion paper, this extra complexity of the molecular machinery can be supported if its own encoding cost is smaller than the information savings it affords across all regulatory sequences in the organism’s genome [22].

Our results call into question the current understanding of regulatory sequence evolution. Instead of focusing on the function, conservation, and evolution of individual binding sites embedded into an assumed “random sequence background” of the CRE, it may be more appropriate to think of entire CREs as individual units of selection. In contrast, studying the evolution of individual BSs in isolation might be misleading, since one has to *a priori* commit to a criterion for what a “functional” binding site is – yet this criterion is itself evolvable within biophysical constraints, as a genetically-encoded trait of the regulatory machinery. If one focuses on CREs as units of selection instead, one finds that evolved CREs are displaced from neutral expectation so that every binding location contributes towards or against the regulatory input from individual TFs: some localized contributions within the CRE are strong but rare, aligning with the textbook notion of a standard binding site; while other contributions might be weak but numerous, yet altogether still accruing to a significant fraction of the total regulatory input. Simultaneously, there may be pervasive selection against noncognate binding across the entire CRE locus that is not, however, localized, so that selection on any individual basepair is weak and would not be evident from genomic data, as assessed by alignment or conservation scores.

We recognize that our conclusions may be limited by the many simplifications we chose to make to bridge the gap between pure theory and some dose of biophysical realism. A key mechanism for eukaryotic gene control that we neglected uses chromatin accessibility, which enables entirely new modes of regulation [22, 28]. Furthermore, it is unclear which of the recently studied regulatory mechanisms such as loop extrusion [47], higher-order chromatin structure control [48–50], transcriptional condensates [51], etc., can be at least approximately brought under the universal umbrella of a two-layer, non-linear, cooperative/synergy-enabling genotype-phenotype map that we studied here. Lessons learned while studying prokaryotic regulation are instructive in this regard: it is almost impossible to guess up front which molecular or physical regulatory mechanisms that can be inferred from experimental data are, in fact, consequential for evolutionary dynamics [12, 21]. Our assumptions about the nature of such dynamics are restrictive as well: the fitness function selects only for zero or full expression in each environment, rather than for precise intermediate gene expression levels; we assume that the fitness function does not change in time and it is unclear whether our results carry over to quasi-stationary regimes where selection itself fluctuates; and we postulate low mutation rates, so that selection does not need to accumulate extra sequence information to maintain mutationally robust solutions in polymorphic populations [22, 34].

Despite these simplifications, we believe in the value of the normative (optimization) approach to understand and rationalize the observed biological complexity. Optimality for a tractably and conveniently chosen function proxy (e.g., a precise regulatory phenotype that we used in this paper) serves as a postulate from which one hopes to mathematically derive quantitative predictions about system’s function that go above and beyond fitting mathematical models to data [52], even when that function proxy is, from an evolutionary standpoint, not very rigorously motivated. And yet, such predictions can be non-trivial and sometimes surprisingly successful when compared to empirical data [53]. In the companion paper [22], we show for the first time that non-trivial optimization over regulatory parameters of the replicated genotype-phenotype map can emerge as an automatic consequence of evolutionary dynamics even when those are not in the *Ns*→ ∞ limit; in the simplified models explored here, we can observe the explanatory power and the non-trivial consequences of this emergent optimization for gene regulation. The theoretical clarity afforded by our approach will hopefully complement the modern, large-scale, data-driven and simulation-based approaches to understand the evolution of gene regulation [12, 14].

## Acknowledgments

Michal Hledik (Institue of Science and Technology Austria) made key contributions to this paper and should thus be listed as a co-author. Since we could not reach him to get his consent for co-authorship, biorxiv policy requires us to list his contribution in the Acknowledgments instead. RB, MH, GT acknowledge the support of the Human Frontiers Science Program Grant RGP0034/2018. We thank Nicholas H Barton for many insightful discussions and substantial contributions to this work, and Fyodor Kondrashov for several key comments about genotype-phenotype maps.

## Notes

### Competing Interest Statement

The authors have declared no competing interest.

## References

1. von Hippel, P. H. & Berg, O. G. On the Specificity of DNA-protein Interactions. Proceedings of the National Academy of Sciences 83 (1986).

2. Bintu, L. et al. Transcriptional regulation by the numbers: models. en. Current Opinion in Genetics & Development. Chromosomes and expression mechanisms 15, 116–124 (Apr. 2005).

3. Kinney, J. B., Murugan, A., Callan, C. G. & Cox, E. C. Using deep sequencing to characterize the biophysical mechanism of a transcriptional regulatory sequence. Proceedings of the National Academy of Sciences 107. Publisher: Proceedings of the National Academy of Sciences, 9158–9163 (May 2010).

4. Ireland, W. T. et al. Deciphering the regulatory genome of Escherichia coli, one hundred promoters at a time. Elife 9, e55308 (2020).

5. De Boer, C. G. et al. Deciphering eukaryotic gene-regulatory logic with 100 million random promoters. en. Nat Biotechnol 38, 56–65 (Jan. 2020).

6. Grah, R., Zoller, B., Tkačik, G., et al. Nonequilibrium models of optimal enhancer function. Proceedings of the National Academy of Sciences 117, 31614–31622 (2020).

7. Villar, D., Flicek, P. & Odom, D. T. Evolution of transcription factor binding in metazoans—mechanisms and functional implications. Nature Reviews Genetics 15, 221–233 (2014).

8. Zheng, W., Gianoulis, T. A., Karczewski, K. J., Zhao, H. & Snyder, M. Regulatory variation within and between species. Annual review of genomics and human genetics 12, 327–346 (2011).

9. Igler, C., Lagator, M., Tkačik, G., Bollback, J. P. & Guet, C. C. Evolutionary potential of transcription factors for gene regulatory rewiring. Nature Ecology & Evolution 2, 1633–1643 (2018).

10. Friedlander, T., Prizak, R., Barton, N. H. & Tkačik, G. Evolution of new regulatory functions on biophysically realistic fitness landscapes. en. Nat Commun 8, 216 (Aug. 2017).

11. Yona, A. H., Alm, E. J. & Gore, J. Random sequences rapidly evolve into de novo promoters. en. Nat Commun 9, 1530 (Dec. 2018).

12. Lagator, M. et al. Predicting bacterial promoter function and evolution from random sequences. eLife 11 (eds Krishna, S., Walczak, A. M., van Nimwegen, E. & Einav, T.) Publisher: eLife Sciences Publications, Ltd, e64543 (Jan. 2022).

13. Fuqua, T. et al. Dense and pleiotropic regulatory information in a developmental enhancer. en. Nature 587. Number: 7833 Publisher: Nature Publishing Group, 235–239 (Nov. 2020).

14. Vaishnav, E. D. et al. The evolution, evolvability and engineering of gene regulatory DNA. Nature 603, 455–463 (2022).

15. Galupa, R. et al. Enhancer architecture and chromatin accessibility constrain phenotypic space during Drosophila development. Developmental Cell 58, 51–62 (2023).

16. Boyle, E. A., Li, Y. I. & Pritchard, J. K. An expanded view of complex traits: from polygenic to omnigenic. Cell 169, 1177–1186 (2017).

17. Ružičková, N., Hledik, M. & Tkačik, G. Quantitative omnigenic model discovers interpretable genome-wide associations. Proceedings of the National Academy of Sciences 121, e2402340121 (2024).

18. Arnold, C. D. et al. Quantitative genome-wide enhancer activity maps for five Drosophila species show functional enhancer conservation and turnover during cis-regulatory evolution. en. Nat Genet 46. Publisher: Nature Publishing Group, 685–692 (July 2014).

19. Ludwig, M. Z., Bergman, C., Patel, N. H. & Kreitman, M. Evidence for stabilizing selection in a eukaryotic enhancer element. en. Nature 403. Publisher: Nature Publishing Group, 564–567 (Feb. 2000).

20. Li, X. C., Fuqua, T., van Breugel, M. E. & Crocker, J. Mutational scans reveal differential evolvability of Drosophila promoters and enhancers. Philosophical Transactions of the Royal Society B: Biological Sciences 378. Publisher: Royal Society, 20220054 (Apr. 2023).

21. Grah, R., Guet, C. C., Tkacik, G. & Lagator, M. Linking Molecular Mechanisms to their Evolutionary Consequences: a primer. bioRxiv, 2024–07 (2024).

22. Hledík, M., Barton, N. H. & Tkačik, G. Evolution and information content of optimal gene regulatory architectures 2025.

23. Berg, J., Willmann, S. & Lässig, M. Adaptive evolution of transcription factor binding sites. BMC Evolutionary Biology 4, 42 (Oct. 2004).

24. Tuğrul, M., Paixão, T., Barton, N. H. & Tkačik, G. Dynamics of Transcription Factor Binding Site Evolution. en. PLOS Genetics 11. Publisher: Public Library of Science, e1005639 (Nov. 2015).

25. Wunderlich, Z. & Mirny, L. A. Different gene regulation strategies revealed by analysis of binding motifs. en. Trends in Genetics 25, 434–440 (Oct. 2009).

26. Cepeda-Humerez, S. A., Rieckh, G. & Tkačik, G. Stochastic proofreading mechanism alleviates crosstalk in transcriptional regulation. Physical review letters 115, 248101 (2015).

27. Friedlander, T., Prizak, R., Guet, C. C., Barton, N. H. & Tkačik, G. Intrinsic limits to gene regulation by global crosstalk. en. Nat Commun 7. Number: 1 Publisher: Nature Publishing Group, 12307 (Aug. 2016).

28. Perkins, M. L., Crocker, J. & Tkačik, G. Chromatin enables precise and scalable gene regulation with factors of limited specificity. Proceedings of the National Academy of Sciences 122, e2411887121 (2025).

29. Stewart, A. J., Hannenhalli, S. & Plotkin, J. B. Why Transcription Factor Binding Sites Are Ten Nucleotides Long. Genetics 192, 973–985 (Nov. 2012).

30. Kribelbauer, J. F., Rastogi, C., Bussemaker, H. J. & Mann, R. S. Low-Affinity Binding Sites and the Transcription Factor Specificity Paradox in Eukaryotes. Annual Review of Cell and Developmental Biology 35. eprint: 10.1146/annurev-cellbio-100617-062719, 357–379 (2019).

31. Shahein, A. et al. Systematic analysis of low-affinity transcription factor binding site clusters in vitro and in vivo establishes their functional relevance. en. Nat Commun 13. Number: 1 Publisher: Nature Publishing Group, 5273 (Sept. 2022).

32. Lynch, M. & Hagner, K. Evolutionary meandering of intermolecular interactions along the drift barrier. en. PNAS 112. Publisher: National Academy of Sciences Section: PNAS Plus, E30–E38 (Jan. 2015).

33. Röschinger, T., Tovar, R. M., Pompei, S. & Lässig, M. Adaptive ratchets and the evolution of molecular complexity en. preprint (Biophysics, Nov. 2021).

34. Hledík, M., Barton, N. & Tkačik, G. Accumulation and maintenance of information in evolution. Proceedings of the National Academy of Sciences 119. Publisher: Proceedings of the National Academy of Sciences, e2123152119 (Sept. 2022).

35. Payne, J. L. & Wagner, A. The causes of evolvability and their evolution. Nature Reviews Genetics 20, 24–38 (2019).

36. Mirny, L. A. Nucleosome-mediated cooperativity between transcription factors. Proceedings of the National Academy of Sciences 107, 22534–22539 (2010).

37. Lleres, D., Gizzi, A. C. & Nollmann, M. Redefining enhancer action: Insights from structural, genomic, and single-molecule perspectives. Current Opinion in Cell Biology 95, 102527 (2025).

38. Kryazhimskiy, S., Tkačik, G. & Plotkin, J. B. The dynamics of adaptation on correlated fitness landscapes. Proceedings of the National Academy of Sciences 106, 18638–18643 (2009).

39. Gillespie, D. T. Exact stochastic simulation of coupled chemical reactions. The journal of physical chemistry 81, 2340–2361 (1977).

40. Wagner, A. Information theory, evolutionary innovations and evolvability. Philosophical Transactions of the Royal Society B: Biological Sciences 372. Publisher: Royal Society, 20160416 (Oct. 2017).

41. Sella, G. & Hirsh, A. E. The application of statistical physics to evolutionary biology. Proceedings of the National Academy of Sciences 102, 9541–9546 (2005).

42. Schneider, T. D. & Stephens, R. M. Sequence logos: a new way to display consensus sequences. Nucleic acids research 18, 6097–6100 (1990).

43. Mustonen, V., Kinney, J., Callan, C. G. & Lässig, M. Energy-dependent fitness: A quantitative model for the evolution of yeast transcription factor binding sites. en. PNAS 105. Publisher: National Academy of Sciences Section: Biological Sciences, 12376–12381 (Aug. 2008).

44. He, X., Samee, M. A. H., Blatti, C. & Sinha, S. Thermodynamics-Based Models of Transcriptional Regulation by Enhancers: The Roles of Synergistic Activation, Cooperative Binding and Short-Range Repression. en. PLOS Computational Biology 6. Publisher: Public Library of Science, e1000935 (Sept. 2010).

45. Reiter, F., Wienerroither, S. & Stark, A. Combinatorial function of transcription factors and cofactors. Current opinion in genetics & development 43, 73–81 (2017).

46. Li, H., Xu, Z., Taylor, G., Studer, C. & Goldstein, T. Visualizing the loss landscape of neural nets. Advances in neural information processing systems 31 (2018).

47. Gabriele, M. et al. Dynamics of CTCF-and cohesin-mediated chromatin looping revealed by live-cell imaging. Science 376, 496–501 (2022).

48. Mach, P. et al. Cohesin and CTCF control the dynamics of chromosome folding. Nature genetics 54, 1907–1918 (2022).

49. Zuin, J. et al. Nonlinear control of transcription through enhancer–promoter interactions. Nature 604, 571–577 (2022).

50. Brückner, D. B., Chen, H., Barinov, L., Zoller, B. & Gregor, T. Stochastic motion and transcriptional dynamics of pairs of distal DNA loci on a compacted chromosome. Science 380, 1357–1362 (2023).

51. Pei, G., Lyons, H., Li, P. & Sabari, B. R. Transcription regulation by biomolecular condensates. Nature Reviews Molecular Cell Biology, 1–24 (2024).

52. Bialek, W. Biophysics: searching for principles (Princeton University Press, 2012).

53. Sokolowski, T. R., Gregor, T., Bialek, W. & Tkačik, G. Deriving a genetic regulatory network from an optimization principle. Proceedings of the National Academy of Sciences 122, e2402925121 (2025).

