## Supplementary material for "Regulatory architectures optimized for rapid evolution of gene expression": SI Appendix, Methods, SI Figures

### Supporting Information Text

**5 Supplementary figures** 22

**6 List of tables** 32

### 1. A simple model of regulatory sequence evolution

We study how the gene regulatory architecture affects the evolvability of cis-regulatory elements (CREs). By regulatory architecture we refer to the components of the regulatory system outside the focal CRE DNA sequence, such as the (hyper)parameters of the genotype-phenotype map (the “regulatory parameters”), the chromatin structure of the DNA, properties of the TFs involved, or the wiring of the gene regulatory network, etc. Our work specifically focuses on one question: What are optimal regulatory parameters for the CRE genotype-phenotype maps that give rise to most rapid CRE adaptation?

We search for an answer in a toy model of gene regulation. Our model has two main building blocks: (i) a biophysically-inspired model of TF-DNA interactions and how these give rise to gene expression, which describes how genetic information translates into expression levels on the timescale at which proteins turn around, and (ii) long-term evolutionary dynamics, rooted in population genetics, which describes how the regulatory sequence itself changes. We start by introducing the biophysically-inspired model of gene regulation.

**A. Biophysically-inspired model of gene regulation.** Transcription factors (TFs) recognize short segments of the DNA, called binding sites (BSs). In metazoans, these sites are typically $6-10$ base pairs long and occur within cis-regulatory elements (CREs) – promoter and enhancer regions – on the DNA.

The interaction strength or binding energy between a TF and a given site depends on how well the site matches the consensus sequence of the TF. The consensus sequence is the DNA sequence that binds the TF the strongest, and by convention, we choose the energy scale such that the binding energy of a consensus site is 0. Consequently, binding to any other site comes at an energy cost. The cost is determined by the number of mismatches in the site compared to the consensus sequence. We assume that each base pair within the binding site contributes independently to the binding and each mismatch carries an energy penalty, $\varepsilon$. The physiological range for $\varepsilon$ is around $1-3\,k_BT$ (1, 2). This mismatch model has been extensively described in the literature (1, 3, 4).

We assume that the functional form of TF binding to the CRE mimics the expressions that can be derived in thermodynamic equilibrium (5). In prokaryotic systems, equilibrium models of transcriptional regulation have been quite successful in their quantitative predictions (6, 7). Whether or not eukaryotic regulation is at equilibrium is still debated and is not a topic of this work; non-equilibrium processes have, for instance, been suggested to provide functional advantages over equilibrium models (e.g., higher sensitivity or specificity) (8, 9). For our purposes, using an equilibrium form for TF-DNA interactions provides tremendous modeling simplicity compared to non-equilibrium models, which tend to have a vast parameter space. But as we will see next, beyond the TF-DNA interaction step, our model also includes nonlinearities which could require non-equilibrium operation. In short, we assume certain functional forms for TF-DNA interaction, cooperativity, and gene activation without committing to whether they do or do not occur at equilibrium, noting that equilibrium models may be sufficient to reach some, but possibly not all (e.g., steep activation nonlinearity), regimes of our model's operation.

We model the occupancy of a site $i$ on the DNA by TF $j$ via its binding energy and the concentration of that TF:

$$\rho_{ij}(c_j) = \frac{c_j}{c_j + e^{\varepsilon k_{i,j}}}, \qquad [1]$$

where $k_{i,j}$ is the number of mismatches in site $i$ with the consensus sequence of TF $j$, and concentration is measured in units such that at $c = 1$ the consensus binding site is half occupied.

In metazoans, high specificity – the ability to express genes at the correct time and in the correct cell – is thought to be achieved through the activation of an entire enhancer rather than specific binding sites within an enhancer (8). Therefore, we consider TF binding throughout the whole CRE, and the overall effect of the binding is "integrated" across all sites within the CRE.

We imagine a sliding window for each TF being run over the entire CRE, whereby a binding site can start at every position along the CRE. The total occupancy of the CRE comes from the cumulative contribution of these overlapping sites and all TFs present on the CRE:

$$\rho(\mathbf{c}) = \sum_{j=1}^{n+1} \sum_{i=1}^{L-\ell+1} \rho_{ij}(c_j), \qquad [2]$$

where we have $n+1$ different types of TF present, as explained in the main text. $L$ is the length of the CRE, and $\ell$ is the length of the TF motifs; $\ell$ is assumed to be the same for all TFs in our model, but this assumption could easily be relaxed.

Gene expression is a noisy and complex process mediated by transcriptional, translational, and post-translational steps. The binding of TFs on CREs is one of the first steps in the regulatory cascade. As we aim for a simple model, we assume the multiple steps following TF binding can be coarse grained into a single nonlinear (sigmoidal) function that determines gene expression. We further only consider activator TFs, which, for simplicity, we assume to act with the same strength, $\delta > 0$ (as explained in the main text; repressors could easily be modeled by having $\delta_j$ separate and not equal for each TF $j$, and requiring $\delta_j < 0$ for repressors). Then we can write gene expression as:

$$g(\mathbf{c}) = \left[1 + e^{-\delta(\rho(\mathbf{c}) - \mu)}\right]^{-1}, \qquad [3]$$

where $\mathbf{c} = \{c_j\}$ encompasses $n+1$ TF concentrations, $\mu$ is the activation threshold of the gene, and $\delta > 0$ controls how smooth this threshold is. $\mu$ and $\delta$ are thus effective parameters describing the complex biology that happens from the binding of the TFs to the actual production of proteins that the gene encodes.

Gene expression conditional on the TF concentrations is the phenotype of our CRE and regulatory system, on which selection can act; therefore Eqs. (1) to (3) constitute the genotype-phenotype map in our model. We also refer to this GP map as a "doubly-nonlinear map": the first, "binding" nonlinearity is the thermodynamically-inspired Eq. (1); the second, "activation" nonlinearity is the sigmoid of Eq. (3).

**B. Types and parameters of the genotype-phenotype maps.** Our main goal is to examine which genotype-phenotype maps allow for fast adaptation of CREs. We do this by comparing between different kinds of genotype-phenotype maps, and by scanning through the regulatory parameter values that determine each kind of map. On each of the maps examined, we play out the CRE evolution at the sequence level, while keeping the GP map constant throughout the simulation. This way, we can scan through various GP maps and look at the evolutionary outcomes.

We looked at 3 different kinds of genotype-phenotype maps depending on how TFs interact with the sequence or expression: (i) activation through monomer binding; (ii) activation through dimer binding; and (iii) activation through a synergistic cooperative effect of bound TFs on expression.

***B.1. Regulation via TF monomer binding.*** In the doubly-nonlinear GP map of Eqs. (1) to (3), activation of the CRE happens through independent TF binding (which we refer to here as "monomer binding") without any direct interactions among TFs, either of the same chemical species or across different species. In this model, biophysical parameters $\varepsilon, c_{\max}, \delta, \mu$ and the combinatorial parameters $\ell, L, n$ influence the GP map and thus have an effect on adaptation times. We keep $L = 256$ bp fixed as this is a reasonable enhancer length, as well as $R = \log_{10}(c_{\max}/c_{\min}) = 2$, which is a typical dynamic range of tunable TF concentrations, and vary the other parameters systematically in two regimes. First, we explore a theoretically interesting limit of low concentration, $c_{\max} \ll 1$ and sharp activation threshold, $\delta \to \infty$, (STLC limit). In this limit, the binding nonlinearity per site $c/(c+K)$ of Eq. (1), where $K = \exp(\epsilon k)$, can be approximated with $c/K$, after which the concentration scale (e.g., $c_{\max}$) can be pulled out of the sum over binding sites in Eq. (2). Then, it can be seen that the concentration scale simply multiplies $\delta$ and rescales $\mu$; the first parameter, $\delta$, is later taken to infinity to generate a sharp threshold, whereas the regulatory parameter $\mu$ is taken to be a free parameter to be optimized over. STLC limit is therefore well defined independently of the precise value of $c_{\max}$, which we also confirm numerically. Following the STLC analysis, we also look into a more realistic regime with a smooth threshold at $\delta = 4$ and optimize over both activation threshold, $\mu$ and concentration, $c_{\max}$.

***B.2. Regulation via TF dimer binding.*** We also add TF–TF cooperativity into our model. One way of doing so is to consider dimerization interactions between TFs on the DNA. In this setup, two TFs bind right next to each other within the CRE, occupying a $\ell = 6 + 6$ bp-long binding site. We consider activation only through dimerization; therefore, binding only one TF is not sufficient to contribute to gene activation. If the binding TFs have different consensus sequences, we call the complex a heterodimer, whereas two TFs with the same consensus sequence form a homodimer. The occupancy of this double-site (made of two shorter half-sites next to each other) is:

$$\rho_{ij} = \frac{c_j^2 e^{-\left[\varepsilon(k_{i_1}+k_{i_2})+E_c\right]}}{1 + c_j e^{-\varepsilon k_{i_1}} + c_j e^{-\varepsilon k_{i_2}} + c_j^2 e^{-\left[\varepsilon(k_{i_1}+k_{i_2})+E_c\right]}}, \qquad [4]$$

where $k_{i_1}, k_{i_2}$ denote the mismatch class of the first and second half-site, respectively, within the long binding site, starting at position $i$. $E_c \leq 0$ is the cooperative energy between the two TFs, in units of $k_BT$. If dimerization of the TFs occurs already in the free solution, $E_c = 0$, while dimerization on the CRE is described by $E_c < 0$. In this latter case, the binding of the first TF makes it energetically favorable for a second TF to bind both the CRE and interact with the first TF. While this is a typical narrative in the biology literature, we note that in equilibrium – if that is indeed relevant – the order of binding events of course cannot matter, and so each scenario will have a matched and balanced time-reversed scenario.

Dimerization occurs both for cognate and noncognate TF pairs. $c_j$ corresponds to the monomer concentration of TF $j$. The rest of the genotype-phenotype map is the same as before, set by Eqs. (2) and (3). Technically, in the simulations, we use a single motif to encode the consensus sequence of the dimerized TF pair comprising two-half sites, which constitutes an identical input as if we were to use a long monomer to bind. However, importantly, the underlying GP map does change via the expression for the occupancy, Eq. (5).

We can use an approximation of this model in the limit of strong cooperativity, i.e., $E_c \ll \ln(c) - \varepsilon k_1$ and $E_c \ll \ln(c) - \varepsilon k_2$, which results in the following occupancy:

$$\rho_{ij} = \frac{c_j^2 e^{-\left[\varepsilon(k_{i_1}+k_{i_2})+E_c\right]}}{1 + c_j^2 e^{-\left[\varepsilon(k_{i_1}+k_{i_2})+E_c\right]}}. \qquad [5]$$

This approximation has the advantage of reducing computing times because mismatch numbers appear only as a sum in Eq. (5). By scanning the consensus motif over the CRE, we can count how many times the mismatch number $k = k_1 + k_2$ appears along the CRE and sum over the contribution according to:

$$\rho = \sum_{k=0}^{\ell=\ell_1+\ell_2} m_k \frac{c_j^2 e^{-(\varepsilon k + E_c)}}{1 + c_j^2 e^{-(\varepsilon k + E_c)}}, \qquad [6]$$

where $\ell = \ell_1 + \ell_2$ is the total length of the site; we use $\ell_1 = \ell_2 = 6$, and $m_k$ is the number of times in the CRE that a mismatch $k$ occurs.

We carried out simulations both for the approximation and the full model (results not shown), and used cooperative energy values $E_c = 0$ or $-4\,k_BT$. As expected, the approximation works better with $E_c = -4\,k_BT$. The outcomes of the full model and the approximation are qualitatively the same. Although the optimal adaptation rates in the approximation model slightly overestimate those of the full model, the estimates remain within $\pm\sigma$ for $E_c = -4$ and within $\pm 1.5\sigma$ for $E_c = 0$, indicating statistical consistency; $\sigma$ indicates the standard deviation over replicate simulations. Throughout the main paper and in the SI figures, we report on the results from the approximation model.

***B.3. Regulation via TF synergy.*** We consider a GP map that features synergistic interactions among TFs, meaning that the cumulative binding strength of the same type of TF is larger than the sum of the individual effects. We calculate binding to an individual site according to Eq. (1), but the total occupancy of the CRE includes an enhanced effect from each type of TF:

$$\rho = \sum_{j=1}^{n+1} \left( \sum_{i=1}^{L-\ell+1} \rho_{ij} \right)^{\eta}, \qquad [7]$$

where $\eta > 1$ is the synergy coefficient. An important assumption, whose relaxation would provide an interesting avenue for future research, is that we assume the synergy to only occur between TFs of the same kind, i.e., for the same $j$. As a more general alternative, the synergy coefficients can be thought of as being arranged into a matrix $\eta_{jk}$ between TFs $j$ and $k$, and this matrix could itself be evolutionarily tuned for evolvability. For simplicity, our analysis takes $\eta_{jk}$ to be a diagonal matrix, with the same value along the diagonal. Biologically, such synergy could be implemented via protein-protein interactions between TFs and the intermediate molecular machinery, such as the Mediator complex, or directly via contacts with the RNAPol complex. Apart from the modified Eq. (7), the rest of the genotype-phenotype map is unchanged: we use Eq. (3) to calculate gene expression.

**C. Evolutionary dynamics.** We now turn to the second part of our model, the evolutionary dynamics. A reasonable assumption is that evolution happens over much longer timescales than the regulatory dynamics, therefore, we only consider gene expressions at steady state and do not model their dynamics. We further ignore the noise in gene regulation and assume that evolution acts on mean gene expression levels, which we calculate according to Eq. (3). The first step in setting up our evolutionary model is to define a target phenotype towards which the CRE will evolve.

***C.1. Target phenotype and fitness function.*** We argue that it is crucial to incorporate crosstalk into our model, for the following reasons: (i) crosstalk may be prevalent in eukaryotic gene regulation given the estimates of intrinsic limit at which it could exert systemic effects (10); (ii) crosstalk is needed to self-consistently deal with *replicated genotype-phenotype maps* (see main text), in an approach where we focus on the evolution of a single CRE but think of the same genotype-phenotype map determining also the response properties at other CREs in the cell. Thus, noncognate TFs for our focal CRE might act as cognate TFs for other CREs elsewhere in the genome, and vice versa. Without including crosstalk, many "reasonable solutions" (CRE sequences) for regulation could exist for CREs considered in isolation, yet they might not be functional in the cellular context of other CREs and noncognate TFs.

We introduce crosstalk by considering $n$ noncognate TFs plus 1 cognate TF in our model. The concentrations of these TFs define specific discrete environments, $e$. For simplicity, we restrict TF concentrations such that a particular TF is either "present" in the cell with high concentration, $c_{j,e} = c_{\max}$, or "absent" i.e., its concentration is low $c_{j,e} = c_{\min} \ll c_{\max}$. Our target phenotype is a particular gene expression pattern for the considered gene over environments, that is, a gene expression as a function of the environments, which can be thought of as a vector of values indexed by possible discrete environments $e$. In the simplest scenario we consider, a gene should be expressed only if the cognate TF is present in a given environment. We note that it would be possible to consider models where intermediate expression levels of target genes are desired, and the TF concentrations can be, correspondingly, adjusted on a continuum, as we have done in a separate, non-evolutionary work (11); embedding such a model into an evolutionary simulation is, however, technically and interpretationally challenging, so we opted for a simpler setup here.

As long as we keep the number and type of TFs fixed and weigh the environments equally, it is sufficient to consider only two extreme environments to gain basic insights, i.e., the "best" and the "worst" conservative case scenario. In the "best" (`ON`) environment, the cognate TF is present at high concentration, $c_{\max}$, whereas noncognate or crosstalking TFs have a significantly lower concentration, $c_{\min} = 10^{-R} c_{\max}$, where $R$ is the dynamic range, or maximal fold-change in base 10, of the TF concentration (our typical choice is $R = 2$). In the `ON` environment, the cognate TF should activate the gene, and the required phenotype is the absolute highest gene expression, $G_{\texttt{ON}} = 1$. In contrast, in the "worst" (`OFF`) environment, the cognate TF is present at low concentrations, $c_{\min}$, and the gene should remain silent ($G_{\texttt{OFF}} = 0$) despite the binding of noncognate TFs all present at high concentrations, $c_{\max}$. If the CRE successfully reaches this ($G_{\texttt{ON}} = 1, G_{\texttt{OFF}} = 0$) target phenotype, it would also function in any other environment $e$, set by a combination of $n + 1$ TF concentrations. Imagine an intermediate environment in which only a fraction of the noncognate TFs are present. If the CRE can withstand the crosstalk pressure from all $n$ noncognate TFs, the regulatory input from fewer noncognate TF won't activate the gene. Correspondingly, if the cognate TF is present together with noncognate TFs, the CRE – adapted to the extreme environments – will turn on the gene, as the regulatory input from the cognate TF alone is sufficient. (As long as the model is restricted to only two discrete levels of concentrations.)

After these preliminaries, we can write the total occupancy of the CRE in the `ON` and `OFF` environments as:

$$\rho_{\texttt{ON}} = \sum_{i=1}^{L-\ell+1} \frac{c_{\max}}{c_{\max} + e^{\varepsilon k_{i,1}}} + \sum_{j=2}^{n+1} \sum_{i=1}^{L-\ell+1} \frac{c_{\min}}{c_{\min} + e^{\varepsilon k_{i,j}}},$$

$$\rho_{\texttt{OFF}} = \sum_{i=1}^{L-\ell+1} \frac{c_{\min}}{c_{\min} + e^{\varepsilon k_{i,1}}} + \sum_{j=2}^{n+1} \sum_{i=1}^{L-\ell+1} \frac{c_{\max}}{c_{\max} + e^{\varepsilon k_{i,j}}}.$$

We used index 1 for the cognate TF and indices $2, \ldots, n+1$ for the noncognate TFs. Total occupancies then translate to gene expression in each environment via Eq. (3):

$$g_{\texttt{ON}} = \left[1 + e^{-\delta(\rho_{\texttt{ON}} - \mu)}\right]^{-1},$$
$$g_{\texttt{OFF}} = \left[1 + e^{-\delta(\rho_{\texttt{OFF}} - \mu)}\right]^{-1}.$$

Our fitness function describes selection towards this target phenotype by penalizing the squared distance from it:

$$F = e^{-\frac{\alpha}{2}\left[(G_{\texttt{ON}} - g_{\texttt{ON}})^2 + (G_{\texttt{OFF}} - g_{\texttt{OFF}})^2\right]}, \qquad [8]$$

where $\alpha$ sets the scale of selection strength, and the target expression in the `ON` environment is $G_{\texttt{ON}} = 1$, and in the `OFF` environment is $G_{\texttt{OFF}} = 0$. We average over the environments, hence the division by 2. While Eq. (8) takes the functional form for stabilizing selection, our specific choice of $G_{\texttt{ON,OFF}}$ values is forcing the gene expression phenotypes to their theoretical bounds, making the effective action of selection rather mimic directional selection models. Simulating selection for intermediate expression values could open up new evolutionary solutions and dynamics, but is beyond the scope of this work.

**C.2. Population genetics.** We consider CRE evolution according to a population genetics model in the regime of low mutation rates and strong selection. In this limit, we can describe the population with one effective genotype because each mutation fixes or gets lost in the population before the next one arises; this is known as "fixed states population" assumption. The fixation probability of a mutation depends on the effective population size, $N$, and its selective advantage, $s$, which is the relative fitness advantage of the new genotype over the old one, $s = \frac{F_{\text{new}} - F_{\text{old}}}{F_{\text{old}}}$. The fixation probability for a haploid population introduced by Kimura (12) is:

$$P_{\text{fix}} = \frac{1 - e^{-2s}}{1 - e^{-2Ns}}. \qquad [9]$$

The strong selection limit means that $Ns \gg 1$ and in our model this approximately implies $N\alpha \gg 1$, as $s \propto \alpha$:

$$s = \frac{F_{\text{new}}}{F_{\text{old}}} - 1 \approx \ln\left(\frac{F_{\text{new}}}{F_{\text{old}}}\right) = \ln(F_{\text{new}}) - \ln(F_{\text{old}}) \propto \alpha.$$

High $N\alpha$ results in a steep fitness landscape, which makes small expression differences matter, thereby decreasing fluctuations in the evolutionary trajectory. In contrast, low $N\alpha$ promotes neutral evolution on a flat fitness landscape, where the governing evolutionary force is drift. Parameter $\alpha$ sets the lowest possible range for fitness values. Given that gene expression $g$ only changes within the interval $(0, 1)$, fitness only varies over $(e^{-\alpha}, 1)$.

After defining the genotype-phenotype-fitness map, our next step is to introduce mutations into the evolutionary model. We treat the evolutionary process as a continuous time discrete space Markov Chain, where mutations arise at a constant rate $u$ per individual per basepair per generation. We describe mutations as a Poisson process with a constant rate, and we assume that they are independent. The time between successive events (mutations) in a Poisson process is exponentially distributed with mean $1/(NLu)$, where $NLu$ is the incoming number of mutations in the population per generation. We only consider point mutations along the CRE, and we use Gillespie Stochastic Simulation Algorithm (13) to simulate the mutational events. Every timestep, a mutation arises and we evaluate the fitness of the CRE through the genotype-phenotype-fitness map, then we either accept or reject the mutation based on Eq. (9); if the mutation is accepted, the CRE sequence updates, modeling the jump of the entire population to a new genotype. A time step until next mutation is drawn from an exponential distribution with scale parameter $1/(NL)$, since we measure time in inverse mutation rate, $1/u$. Simulations start at $T_0 = 10^{-5}\,\frac{1}{u}$ and run until a fit solution is reached or until $T_{\text{sim}} = 100\,\frac{1}{u}$ with baseline parameters $N = 100, \alpha = 1, R = 2$. Simulation time, $T_{\text{sim}}$, greatly varies with $N\alpha$, see Section 4.B.

**D. Simulation setup.** We start each evolutionary trajectory from a randomly generated CRE and a set of $(n + 1)$ consensus sequences. Consensus sequences are randomly and independently generated per replicate. Nucleotide frequencies are assumed to be uniform, with probability $f = 0.25$ for each letter in the DNA alphabet {`A, T, C, G`}. The only constraint we impose is that consensus sequences of $(n+1)$ TFs should not be too similar, therefore, we make sure that within each set of TF consensus sequences, each pair of consensus sequences differs in at least 3 positions (regardless of the length, $\ell$) (10). In a more complete framework, we could also treat consensus sequences as regulatory parameters – and this is, actually, how we formally denote them in the main paper. In such a framework, we could scan or optimize over the discrete consensus sequence values to find those sets that would maximize adaptation rates. This discrete optimization problem is extremely time consuming on the one hand; on the other, previous work actually suggests the optimal "placement" of consensus sequences in the sequence space. In our case, the cognate TF consensus sequence should be as different from the non-cognate ones, while non-cognate ones should be as similar to each other as possible. This solution, however, cannot be admissible for a replicated GP map, where the roles of cognate and non-cognate TFs are exchanged at other CREs: if, for our focal CRE, the non-cognate TFs are made very similar to each other in their consensus sequence, these same TFs will have a large crosstalk on other CREs where one of them will be cognate and others non-cognate. The only admissible solution is thus to space all TF consensus sequences as far apart from each other in the sequence space as possible to minimize crosstalk (10). Following this argument, we consider TF consensus sequences that differ sufficiently from each other. This arrangement has been arrived at also in previous work that looked at evolutionary simulations to study the co-evolution of TFBS and TF consensus sequences (14).

We typically run at least 100 replicate simulations starting from such sequence sets of one CRE of length $L = 256$ and $n + 1$ consensus sequences of length $\ell$. For parameter optimization over $c_{\max}$ and $\mu$, we always use the same set of 100 starting sequences — as long as parameters $(L, \ell, n)$ do not change; this reduces the variance and eliminates possible biases for side-by-side comparisons of evolutionary outcomes and adaptation rates at different regulatory parameters.

We typically run evolutionary simulations until a fit phenotype is reached, but no longer than $T_{\text{sim}} = 100\,\frac{1}{u}$ at $N = 100$ and $\alpha = 1$, which means on average a maximum of $100/(NL)^{-1} = 2.56$ million proposed mutations. Simulation time strongly depends on $N\alpha$, and with low $N\alpha$, $(N\alpha \ll 1)$ we go up to $T_{\text{sim}} = 10000\,\frac{1}{u}$.

Parameter ranges that were explored in our simulations are summarized in Table S1.

### 2. Information theoretic quantities and analyses

The Shannon entropy of a random variable $X$ quantifies the uncertainty or unpredictability associated with the variable's possible states, and it is calculated as:

$$H(X) = -\sum_{x \in X} P(x) \log_2 P(x). \qquad [10]$$

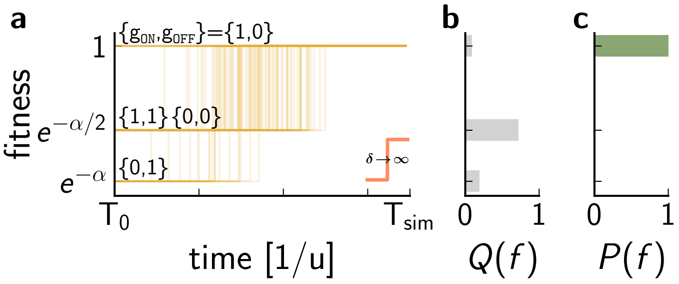

**Fig. F1. Phenotype and fitness states in the STLC limit.** **(a)** Example fitness trajectories in the STLC limit, using parameters that guarantee an IT-efficient solution (see main text). Neutral sequences start at 3 different fitness values that are determined by possible gene expression states in the two environments, marked by $\{g_{\texttt{ON}}, g_{\texttt{OFF}}\}$. **(b)** Neutral fitness distribution, $Q(f)$ at $\mathrm{T}_0$. The most populated state, at this particular setting of the regulatory parameters, is one in which the CRE satisfies the target phenotype in one of the two environments. **(c)** Adapted fitness distribution, $P(f)$ at $\mathrm{T}_{\mathrm{sim}}$. In this IT-efficient example, all trajectories reach a high fitness solution ($f = 1$) within the simulated time. Parameters: $L = 256, n = 3, \varepsilon = 3, \ell = 8, R = 2, c_{\max} = 0.1, \mu = \mu^*, N = 100, \alpha = 1$.

We measure information in *bits*, hence the base 2 in the logarithm. $P(x)$ is the probability of encountering state $x$ of the random variable $X$. Shannon entropy is maximized if the probability of states is uniform; for a discrete random variable defined over $|X|$ states, the entropy of the uniform distribution will be $\log_2 |X|$ bits.

The Kullback-Leibler divergence (KL divergence), or relative entropy between distributions $P$ and $Q$, measures how different the two distributions are,

$$D_{KL}(P||Q) = \sum_x P(x) \log_2 \frac{P(x)}{Q(x)}. \quad [11]$$

It is zero when $P(x) = Q(x)$ for all $x \in X$, and is positive otherwise. We can use KL divergence in an evolutionary setting by associating $Q$ and $P$ with neutral and adapted distributions (e.g., of population states, genotypes, phenotypes, or fitness), in which case the KL divergence estimates how much "work" selection has done by displacing the selected-for from the neutral distribution. Universal, information-theoretic bounds may relate such measures to each other, depending on the choice of $X$ (15).

**A. Fitness-level information in the sharp threshold low concentration (STLC) limit.** In the limit of sharp activation threshold ($\delta \to \infty$), we argue below that the fitness level information defined as a KL divergence between neutral and adapted fitness distributions over all genotypes, equals the phenotypic information defined by Wagner (16). In the sharp threshold limit, genotype-phenotype maps reduce to a step-function, making only two gene expression or phenotype states available: 0 and 1. These states map onto 3 discrete fitness values, as shown in Fig. F1. As a small technicality: when counting the possible states of the system, we omit cases where the total regulatory input would be exactly equal to the activation threshold – resulting in $g = 1/2$ – these outcomes are formally possible as in the simulations we use a finite value for $\delta$, $\delta = 10^7$, but we never observe them in practice. The neutral fitness distribution depends on the particular choice of activation threshold $\mu$: in the STLC limit, $\mu$ is the sole regulatory parameter determining the GP map and hence governing the distribution over the 3 discrete fitness states. Adapted fitness distributions, on the other hand, always consist of a single peak at $f = 1$, as a consequence of (by definition) a sharp GP map in the STLC limit. An example is shown in Fig. F1. Functional sequences map onto fitness $f = 1$, while non-functional sequences have fitness $f < 1$. We can use the KL divergence to quantify by how much adaptation under selection has reshaped the fitness distribution from neutrality:

$$\begin{aligned} \mathcal{D}(\mu) = D_{KL}(P||Q)|_\mu &= \sum_{f \in F} P(f) \log_2 \frac{P(f)}{Q(f,\mu)} \\ &= \sum_{f<1} P(f) \log_2 \frac{P(f)}{Q(f,\mu)} + \sum_{f=1} P(f) \log_2 \frac{P(f)}{Q(f,\mu)} \\ &= 0 + \sum_{f=1} P(f) \log_2 \frac{P(f)}{Q(f,\mu)} \\ &= P(f=1) \left[ \log_2 P(f=1) - \log_2 Q(f=1,\mu) \right] \\ &= -\log_2 Q(f=1,\mu), \end{aligned}$$

where we first sum over non-functional and functional sequences separately. The sharp threshold makes sure that in the adapted distribution, fitness values below one have a probability of zero, e.g. $P(f < 1) = 0$ and consequently $P(f = 1) = 1$. Therefore,

KL divergence in this limit can be estimated from the neutral fitness distribution alone, as it only depends on $Q(f=1)$, the probability of finding a functional sequence in the neutral distribution. In a finite sample we can approximate $Q(f=1)$ with the fraction of functional sequences, $p_\rho$ via

$$Q(f=1,\mu) = \frac{\#\text{ functional sequences at }\mu}{\text{total sample size}} = p_\rho(\mu).$$

Therefore, in the STLC limit, this information estimate, derived from the KL divergence between fitness distributions (where in the derivations above we have made the dependence on the regulatory parameter $\mu$ explicit) quantifies how "difficult" it is for selection to find fit solutions:

$$\mathcal{D}(\mu) = -\log_2 p_\rho(\mu). \quad [12]$$

Clearly, Eq. (12) also equals the phenotypic information proposed by Wagner (16).

***A.1. Information estimates in the STLC limit.*** Functional sequences are ones that generate high expression in the `ON` environment and low expression in the `OFF` environment. In the $\delta \to \infty$ limit, gene expression is defined by the activation threshold, $\mu$, and the total binding occupancy in the environment (Eqs. (3) and (22)). We denote the total binding occupancy in the `ON`, (`OFF`) environment by $\rho_{\texttt{ON}}, (\rho_{\texttt{OFF}})$. We get the desired expression pattern whenever

$$\rho_{\texttt{ON}} \geq \mu \quad \text{and} \quad \rho_{\texttt{OFF}} < \mu \quad [13]$$

are true simultaneously. In this limit, a CRE has high fitness ($f=1$) if and only if the sequence satisfies Eq. (13); therefore, counting functional sequences in the intermediate phenotype space $(\rho_{\texttt{ON}}, \rho_{\texttt{OFF}})$ is equivalent to counting high fitness sequences. This saves us two steps in the calculation – explicit calculation of expression and fitness values – yet results in the same information estimate.

Whenever we report $\mathcal{D}(\mu)$, we randomly sample $10^7$ sets of random CRE plus $(n+1)$ consensus sequences, calculate $\rho_{\texttt{ON}}$ and $\rho_{\texttt{OFF}}$ for each generated set, and count how many of these fulfill the requirement set by Eq. (13) at a given activation threshold, $\mu$. Normalizing this by the sample size gives us an estimate of the information, $\mathcal{D}(\mu)$ via Eq. (12) at the chosen $\mu$; to plot the $\mu$ dependence of $\mathcal{D}$, we repeat this calculation at various thresholds.

**B. Lower bound on information.** Information measure $\mathcal{D} = \mathcal{D}(\mu)$ as well as other information measures introduced in follow-up sections are lower bounded by the combinatorics, set solely by number of crosstalking TFs we consider. We can calculate this "information-theoretic lower bound" analytically in a slightly altered version of the model, where the target phenotype is easier to characterize mathematically.

In our setup, there is always one cognate TF (denoted as $\text{TF}_1$), and $n$ noncognate TFs (denoted as $\text{TF}_2 \ldots \text{TF}_{n+1}$). In the altered version of our model, we make two key assumptions: (i) we classify a CRE as functional whenever $\text{TF}_1$ activates the gene, e.g. $\rho_1 \geq \mu$, but none of the other TFs individually do, $\rho_2 < \mu, \rho_3 < \mu \ldots \rho_{n+1} < \mu$, where the total regulatory input of TF $j$ is $\rho_j$. (Note that this is a necessary but not sufficient condition for a functional CRE in our original model, where the *sum* of noncogante contributions should stay below $\mu$.) (ii) We assume that the consensus sequences of the TFs are dissimilar enough such that we can think of $\rho_1, \ldots, \rho_{n+1}$ as independent and identically distributed across random CREs.

In this case, the probability of any TF $j$ binding the CRE successfully, e.g. $\rho_j \geq \mu$ is $q$. Then we can approximate the fraction of random CREs that are functional with $q(1-q)^n$ because $\rho_1 \geq \mu$ has probability $q$ and $\rho_2 < \mu, \rho_3 < \mu \ldots \rho_{n+1} < \mu$ all happen with probability $(1-q)$. This probability depends on $\mu$ such that $q = q(\mu)$.

The fraction of functional random CREs is maximized when information measure,

$$\mathcal{D} = -\log_2\left(q(1-q)^n\right) \quad [14]$$

is minimized. We can calculate the minimum information required by solving

$$\left.\frac{\partial \mathcal{D}}{\partial q}\right|_{q=q^*} = 0, \quad [15]$$

which yields

$$q^* = \frac{1}{1+n} \quad [16]$$

and the minimum information

$$\mathcal{D}^*_{\min}(n) = -\log_2\left[\frac{1}{n+1}\left(1-\frac{1}{n+1}\right)^n\right] = (n+1)\log_2(n+1) - n\log_2 n. \quad [17]$$

This provides a lower bound on information that does not depend on any biophysical parameters (such as $\mu$), but only on the number of crosstalking TFs involved. Loosely speaking, while various constraints, imperfections, and limitations due to the physical implementation of the regulation can increase the needed information above this bound (known as "overspecification"), as analyzed in the companion paper, reliable regulation that would require information *below* the bound is information-theoretically impossible. This is why we refer to any biophysically-realistic regulatory scheme that requires information $\mathcal{D} \approx \mathcal{D}_{\min}$ as IT-efficient – it is the best possible, in an absolute, information-theoretic sense. For example, our gene expression model approximately reaches this lower bound in the STLC limit whenever binding sites are longer than 7 bp and the mismatch penalty is higher than 2 $k_BT$ (see main text figures).

We can also write the lower bound in terms of the Bernoulli information:

$$\mathcal{D}^*_{\min} = (n+1)H\left(\frac{1}{n+1}\right). \quad [18]$$

**C. Fitness-level information on realistic GP maps.** In this section, we describe two ways to estimate information quantifying how much "work" the selection has done to displace the random CRE ensemble towards the functional ensemble at the fitness level, for maps where the threshold $\delta$ is not infinite.

The first method calculates the KL divergence between adapted and neutral fitness distributions as follows:

$$D_{\mathrm{fit}}|_\lambda = \sum_{f\in F} P(f,\lambda)\log_2\frac{P(f,\lambda)}{Q(f,\lambda)},$$

where $\lambda = \{n, \ell, e, R, c_{\max}, \mu, \delta, N, L\}$ are the regulatory parameters that determine the GP map properties on which the adaptation unfolds. While we leave out $\lambda$ from our notation in what follows, it is important to keep in mind that the fitness level information always depends on the particular GP map we are considering.

To estimate the neutral fitness distribution, $Q(f,\lambda)$, we generate a random ensemble of $n+1$ TF consensus and CRE sequence sets and calculate their fitness. We generate these sequence sets explicitly because we can not describe the distributions $Q$ analytically, due to the possibility that the binding sites in the sequences overlap. The natural approximation via a multivariate Gaussian would not work here because these distributions (i.e., distributions of regulatory input phenotypes $\rho$ or fitness $f$) are highly non-Gaussian.

We estimate the steady-state fitness distribution under selection, $P(f,\lambda)$, from the equilibrated parts of the simulated evolutionary trajectories. To sample the adapted fitness distribution, we sample every fitness value in the stochastic simulation across 100 replicate trajectories, after the simulated fitness first exceeds a threshold level of fitness, which we define as $F_{\mathrm{high}} \geq 0.99^\alpha$ (see below). In detail, we only record a sample if a proposed mutation has been accepted and thus the fitness value has changed; rigorously, we should weigh these stationary fitness values with the time spent at each particular fitness peak. To save the calculation time, we avoided the weighting, which biases our estimate slightly towards lower fitness values as trajectories tend to spend more time at higher fitness peaks. We did study this bias, however, to show that it is typically negligible, as shown in Fig. F2a.

Following this procedure, we can estimate the KL divergence, $D_{\mathrm{fit}}$, between adapted and neutral fitness distributions. The mechanics of the calculation are illustrated in Fig. F2b-c. This information is always going to be higher than the information estimated using a sharp activation threshold, $D_{\mathrm{fit}}(\delta = 4) > \mathcal{D} = D_{\mathrm{fit}}(\delta \to \infty)$. On a smooth GP map, selection needs to push the total occupancy of a sequence significantly above the midpoint of the sigmoid for the cognate factors and significantly below the midpoint for the noncognate factors; whereas with a sharp threshold, it is enough to push the sequence to be just slightly above (below) the midpoint. Extra information needed to support this effect is called "overspecification" (15).

The second method to estimate information on the fitness level is based solely on the neutral fitness distributions, and is closely related to the thresholding approach described in the previous section. We simply count how many sequences fall above a fitness threshold, $F_{\mathrm{thr}}$, in the neutral distribution and calculate information from the fraction of high-fitness sequences, $p_{F>F_{\mathrm{thr}}}$, by computing:

$$\tilde{D}_{\mathrm{fit}} = -\log_2 p_{F>F_{\mathrm{thr}}}. \quad [19]$$

This estimation is methodologically more problematic than the first one. While we avoid dealing with adapted fitness distributions and thus can eliminate time-consuming evolutionary stochastic simulations, we need to make an arbitrary choice about the fitness threshold. The choice of threshold is not obvious, because fitness trajectories fluctuate with different magnitudes in the stationary period, depending on the regulatory parameters. Given this caveat, we used $D_{\mathrm{fit}}$ as the fitness level information measure throughout the main text and SI. We show a comparison between $\tilde{D}_{\mathrm{fit}}$ and $D_{\mathrm{fit}}$ in Fig. F2d.

**D. Mismatch-level information.** We can also quantify information on the level of mismatch distributions. We "scan" a given TF consensus sequence across the entire CRE and accumulate the distribution of mismatches, $P_j(k)$, by considering TF binding within any position of the CRE for each TF $j$, across an ensemble of random or adapted sequences. We then calculate the KL divergence between the adapted and neutral mismatch distributions on a particular GP map. The neutral mismatch distribution can be approximated by a binomial distribution, assuming that contributing sites are independent. Because of the overlaps of binding sites, the independence assumption is broken, but in the case of a long enough random sequence, the binomial approximation can still be very reasonable. We also assume equal background frequencies over the 4 nucleotides, consistent with our simulation setup. The neutral mismatch distribution is

$$Q(k) = \mathrm{Binom}\left(\ell, \frac{3}{4}\right) = \binom{\ell}{k}\left(\frac{3}{4}\right)^k\left(1-\frac{3}{4}\right)^{\ell-k}, \quad [20]$$

where $k$ refers to the number of mismatches, as in the main paper. For calculating the adapted mismatch distribution, we take all CREs from the stationary segment of the simulation (after first exceeding $F = 0.99$) of the 100 replicate evolutionary trajectories. (Note that this gives us an aggregated dataset over millions of CREs for one cognate factor and $n$ noncognate

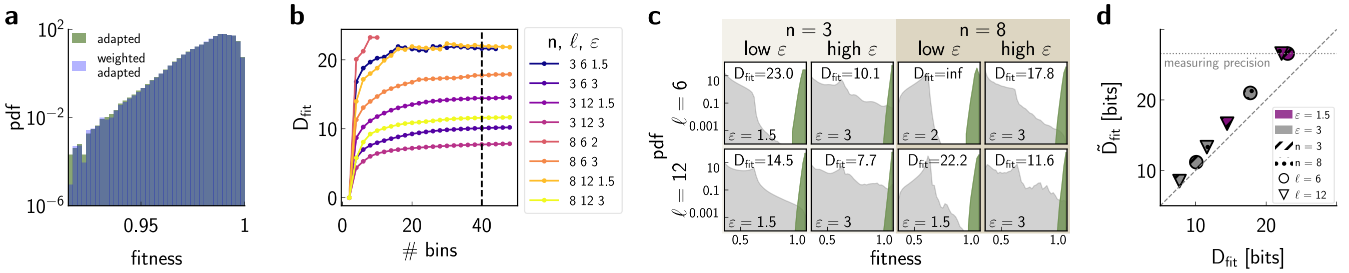

**Fig. F2. Estimating fitness level information on realistic GP maps. (a)** An example of an adapted fitness distribution calculated over the stationary segment of 100 replicate evolutionary trajectories using only substitution events (green) or weighted fitness values (blue). Weights are calculated according to the time spent at each fitness peak. Faster-to-compute estimate (green), which we use for $D_{\mathrm{fit}}$ calculation, is only slightly biased toward lower fitness values, and the difference between the distributions is negligible. **(b)** $D_{\mathrm{fit}}$ dependence on the choice of fitness binning (to estimate the distributions that enter the KL divergence calculation), at various $n, \ell, \varepsilon$ parameters. Horizontal axis shows the number of bins over the possible fitness interval $(e^{-1}, 1)$ at $\alpha = 1$. We use equally sized bins, and choose 40 bins to calculate the $D_{\mathrm{fit}}$ measure (indicated by the black dashed line), as the information estimates nearly saturate at that binning resolution; going significantly beyond 40 bins increases the risk of undersampling and biases in information estimates. **(c)** Neutral (gray) and adapted (green) fitness distributions at optimal $(\mu^*, c^*_{\mathrm{max}})$ values for various $n, \ell, \varepsilon$ parameters. $D_{\mathrm{fit}}$ is indicated in each subplot separately. For the neutral distributions, we sampled ensembles comprising $10^8$ random sequences. In the case of severe crosstalk, at $n = 8, \ell = 6, \varepsilon = 2$, the neutral distribution is undersampled, preventing us from estimating $D_{\mathrm{fit}}$ reliably. **(d)** $\tilde{D}_{\mathrm{fit}}$ at $F_{\mathrm{thr}} = 0.99$ slightly overestimates $D_{\mathrm{fit}}$, which is a theoretically more rigorous, and a more restrictive measure. Estimation of $\tilde{D}_{\mathrm{fit}}$ also breaks down at multiple parameter sets: $n, \ell, \varepsilon = \{8, 12, 1.5\}, \{3, 6, 1.5\}, \{8, 6, 2\}$. This is either because we do not sample fitness values above $F = 0.99$ or only sample one such value, risking large estimation errors. Parameters: $L = 256, R = 2, \delta = 4, \alpha = 1, N = 100$; for **(a)**: $n = 8, \ell = 6, \varepsilon = 3$

ones). We aggregate mismatch counts across all $n$ noncognate TFs (and later scale the results appropriately with $n$, wherever needed). This leaves us with an adapted cognate and noncognate mismatch distributions, $P_{\mathrm{cog.}}(k)$ and $P_{\mathrm{noncog.}}(k)$. We obtain KL divergences separately for the cognate and noncognate site distributions:

$$D^{\mathrm{site}}_{\mathrm{cog.}} = \sum_k P_{\mathrm{cog.}}(k) \log_2 \frac{P_{\mathrm{cog.}}(k)}{Q(k)}$$

$$D^{\mathrm{site}}_{\mathrm{noncog.}} = \sum_k P_{\mathrm{noncog.}}(k) \log_2 \frac{P_{\mathrm{noncog.}}(k)}{Q(k)}.$$

These KL divergences quantify how different the adapted and the neutral distributions are at the level of a binding site; we scale them up by the number of sites in a CRE to obtain the CRE level information:

$$D_{\mathrm{cog.}} = (L - \ell + 1) D^{\mathrm{site}}_{\mathrm{cog.}}$$

$$D_{\mathrm{noncog.}} = (L - \ell + 1) D^{\mathrm{site}}_{\mathrm{noncog.}}.$$

We define the information on the mismatch level as the additive contribution of cognate and scaled noncognate CRE level information:

$$D_{\mathrm{mm}} = D_{\mathrm{cog.}} + n D_{\mathrm{noncog.}}. \qquad [21]$$

When calculating adapted mismatch distributions, we can limit ourselves to using only one CRE sequence per replicate (the first sequence to exceed a fitness threshold $F > 0.99$), instead of sampling many sequences from the stationary segment of each single evolutionary trajectory; this approximation is $\tilde{D}_{\mathrm{mm}}$. Computing $\tilde{D}_{\mathrm{mm}}$ is computationally less intensive than computing $D_{\mathrm{mm}}$. We show in Fig. S7 that $\tilde{D}_{\mathrm{mm}}$ predicts very well the range of concentrations and activation thresholds where fastest adaptation of CREs takes place.

It is important to realize that $D_{\mathrm{mm}}$ treats binding sites as independent and does not account for correlations between them. There are multiple reasons why this assumption is violated in practice. First, because TFs can bind at every position in the CRE, the mismatch count at position $i$ is not fully independent of mismatch counts at positions $i'$ for which $|i - i'| < \ell$. Second, the presence of a location within the CRE with a small mismatch count (i.e., a "bona fide" binding site) changes the probability that another binding site will evolve in the same CRE. Third, the fact that a given position has a particular mismatch for TF $j$ is not independent of the mismatch it will have with TF $j'$. Nevertheless, the independent sites assumption, an thus $D_{\mathrm{mm}}$ calculation, can provide a good and very quick to compute approximation, as long as consensus sequences are not biased towards translationally invariant ones (e.g. `AATTAATT`) and the CRE is sufficiently long ($L \gg \ell$). In the main paper and SI figures we compare $D_{\mathrm{mm}}$ measure to alternative information measures to evaluate the effect of the independence assumption.

**Relationship between genetic and mismatch-level information** Genetic information, defined as the KL divergence of adapted ($\psi(\mathbf{s})$) and neutral ($\varphi(\mathbf{s})$) sequence distributions, $D_{\mathbf{s}} = \sum_{\mathbf{s}} \psi(\mathbf{s}) \log_2(\psi(\mathbf{s})/\varphi(\mathbf{s}))$, upper bounds the information extractable from our system. In this paragraph we elaborate on this statement and describe how mismatch distributions $P(k)$ (adapted) and $Q(k)$ (neutral) are derived as marginals of the respective genotype distributions, $\psi(\mathbf{s})$ and $\varphi(\mathbf{s})$.

First we can think of marginalizing the sequence distributions with respect to the mismatch vector $\mathbf{k} = (k_1, k_2, \ldots, k_{L-\ell+1})$, whose elements indicate mismatch counts at each sequence window $i \in \{1, 2, \ldots, L - \ell + 1\}$ along the CRE:

$$P(\mathbf{k}) = \sum_{\mathbf{s}:\mathbf{k}(\mathbf{s})=\mathbf{k}} \psi(\mathbf{s}),$$

where $\mathbf{k}$ is a mismatch vector computed with a particular TF's consensus sequence, and we picked the adapted distribution to illustrate the computation (which is identical for neutral distributions). We next marginalize $P(\mathbf{k})$ over positional windows $i$, writing the probability of finding $k$ mismatches at sequence window $i$:

$$P_i(k) = \sum_{\mathbf{k}:k_i=k} P(\mathbf{k}).$$

Then the global mismatch distribution is the average of such marginals:

$$P(k) = \frac{1}{L-\ell+1} \sum_{i=1}^{L-\ell+1} P_i(k).$$

This series of dimensionality reductions from the genotype distribution to the mismatch distribution shows that the mismatch distribution is a summary statistic of the full sequence ensemble. $P(k)$ can be obtained easily, albeit at the cost of losing information on the position of mismatches within a site, correlation between sites, and position of sites along the CRE. Our simple model of gene expression weighs all positions equally and all mismatches equally, so this reduction of dimensionality cannot not lose information about the phenotype for our model – even though it would, in reality, where these assumptions would not hold. The only discrepancy between mismatch-level information and genetic information, for our model, thus arises because of the site independence assumption.

KL divergences between distributions of summary statistics (e.g., marginal distributions) are less than or equal to the KL divergence between the respective original distributions. Thus, if we were concerned about a single cognate TF only (i.e., for $n = 0$), then $D_\mathbf{s} \geq D_\mathrm{mm}$ should hold strictly. For $n > 0$, a full account of mismatches would involve accumulating the joint distribution over mismatches for all $(n + 1)$ TFs, i.e., $P(k_1, k_2, \ldots, k_{n+1})$, and this could be used to compute the multi-TF generalization of the mismatch distribution. Unfortunately, even though it provides a drastic dimensionality reduction compared to the full genotype distributions, a joint distribution over mismatches still suffers from a curse of dimensionality to the extent that it makes it not tractable in practice.

Our way of dealing with the $n > 0$ case is to calculate $D_\mathrm{mm}$ as a sum over KL divergences of marginal distributions for cognate and non-cognate factors as in Eq. (21). The sum of KL divergences is still less than or equal to the KL divergence of genotype distributions if the summed mismatch statistics are independent (i.e., if the joint distribution over mismatches factorizes), and there is no overlap in the information these extract; otherwise the inequality between the mismatch-level information and genotypic information is not guaranteed. Nevertheless, due to the simplicity of computation and interpretation, and because for long CRE sequences and random consensus sequences it provides an informative measure, we use $D_\mathrm{mm}$ extensively in our work.

We note that even though simple in a simulational setting, applying $D_\mathrm{mm}$ to real DNA sequences would be non-trivial because we usually do not know all the TFs that are in play for a particular CRE, let alone can identify the TFs that should *not* bind a given CRE.

### 3. Detailed analyses in the sharp-threshold low-concentration (STLC) regime

STLC is an interesting theoretical limit of our model, with a sharp activation threshold ($\delta \to \infty$) and low concentrations ($c_\mathrm{max} \ll 1$). Examining evolutionary outcomes in this setup greatly helped us understand the effect of crosstalk and establish the connection between evolutionary dynamics and information theory.

In the $\delta \to \infty$ limit, the map between total regulatory input and phenotype is a step function, uniquely determined by a single parameter, the activation threshold $\mu$. Fitness correspondingly takes on a small, countable set of possible values (depending on the number of environments and non-cognate TFs), greatly simplifying the reasoning and calculations.

In the STLC limit, the gene expression of Eq. (3) reduces to:

$$g(\mathbf{c}) = \begin{cases} 1, & \rho - \mu \geq 0 \\ 0, & \rho - \mu < 0. \end{cases} \quad [22]$$

Fitness trajectories mirror the sharpness of the genotype-phenotype map, and evolutionary outcomes depend on the exact value of $\mu$. Large $\mu$ values necessitate the emergence of multiple strong binding sites for the cognate factor, which take a long time to evolve and thus result in low adaptation rates. Small $\mu$ values are too permissive and lead to erroneous expression under the activating input of noncognate TFs. Pruning away promiscuous binding of noncognate TFs across the entire CRE also takes a long time to evolve, and likewise leads to low adaptation rates. In the main paper, we report on the existence of optimal intermediate $\mu$ values, at which adaptation of CREs proceeds orders of magnitude faster than away from the optimum, at extremal $\mu$ values.

The total regulatory input of all TFs, $\rho$, is proportional to $c_{\max}$ in the limit of $c_{\max} \ll 1$. We can see this by first looking at the Taylor expansion of Eq. (1), i.e., at the binding probability of TF $j$ at site $i$ around $c_j = 0$. This gives us:

$$\rho_{ij}(c_j = 0) \approx c_j e^{-\varepsilon k_{i,j}}, \quad [23]$$

which makes the total regulatory input of TF $j$ proportional to its concentration, $\rho_j = c_j \sum_{i=1}^{L-\ell+1} e^{-\varepsilon k_{i,j}}$. Regulatory input in the ON environment comes from the cognate TF at $c_{\max}$ and from the additive contribution of noncognate TFs at $c_{\min}$. $\rho_{\texttt{ON}}$ then takes the form:

$$\rho_{\texttt{ON}} = c_{\max} \underbrace{\sum_{i=1}^{L-\ell+1} e^{-\varepsilon k_{i,j}}}_{A} + c_{\min} \underbrace{\sum_{j=2}^{n+1} \sum_{i=1}^{L-\ell+1} e^{-\varepsilon k_{i,j}}}_{B} = c_{\max}(A + 10^{-R} B), \quad [24]$$

where we used the notation $A, B$ for the residual cognate and noncognate contributions respectively. Similarly, in the OFF environment:

$$\rho_{\texttt{OFF}} = c_{\min} \sum_{i=1}^{L-\ell+1} e^{-\varepsilon k_{i,j}} + c_{\max} \sum_{j=2}^{n+1} \sum_{i=1}^{L-\ell+1} e^{-\varepsilon k_{i,j}} = c_{\max}(10^{-R} A + B), \quad [25]$$

which makes the total occupancy of the CRE in both environments proportional to $c_{\max}$. We can further approximate $\rho_{\texttt{ON}}$ and $\rho_{\texttt{OFF}}$ by only considering the main contributing factors in each environment. Assuming that $10^{-2} n c_{\max} \approx 0$, since $c_{\max} \ll 1$:

$$\rho_{\texttt{ON}} = A c_{\max},$$
$$\rho_{\texttt{OFF}} = B c_{\max}.$$

Note that this approximation relies on the number of non-cognate TFs, $n$, to be sufficiently small in Eq. (24), since the neglected term also scales with $n$. This is true for the parameters we explored in this study.

Combining these results with Eq. (22), we conclude that the CRE is functional if the following holds:

$$\rho_{\texttt{ON}} = A c_{\max} \geq \mu \quad \text{and} \quad \rho_{\texttt{OFF}} = B c_{\max} < \mu.$$

In the limit of $\delta \to \infty$ for any $0 < \mu < 1$, there exists a concentration that makes adaptive evolution of the CRE fastest, and regulation takes place via one evolved cognate binding site, as shown in Fig. F5. Low concentration means that site occupancies are very low as well. Regulation at vanishingly low concentration can only work in the $\delta \to \infty$ limit, because we can optimize activation thresholds $\mu$ such that one site with the highest cognate occupancy matters for expression, *even when its occupancy is very small in an absolute sense, i.e., compared to* 1*, which is full occupancy of a BS by a TF.* In this optimal regime, the occupancy of the cognate site is comparable to $\mu$, but other (noncognate) contributions from sites with even lower occupancies are silenced. This one site is therefore "strong enough" even though its occupancy might be low in absolute terms, on the available occupancy scale of $(0, 1)$.

It is not surprising that only one cognate site emerges in the CRE as an optimal solution in this regime. At the optimal $\mu$, there is a single type of mutation that has a significant fitness effect: the mutational step that changes a given site into a strong-enough cognate site (beneficial mutation) or a strong-enough noncognate site (deleterious mutation). That means the evolutionary search needs to stumble upon such a "lucky" mutation blindly (while avoiding "unlucky" ones), in the absence of any selection gradients. Outside of the optimal regime, the CRE adaptation dynamics get much slower, as multiple fixations are needed either to get rid of noncognate binding sites and/or to create multiple cognate sites.

Site occupancy by a TF depends on the TF concentration, mismatch penalty, and motif length. Large $\varepsilon$ and longer motifs both decrease occupancy, therefore optimal $\mu$ values are smaller for long sites with a high mismatch penalty compared to short sites and low $\varepsilon$, at the same concentration. Higher concentrations increase site occupancy, thus necessitating larger optimal $\mu$ values.

**A. Neutral sequence distributions in regulatory input space.** In the STLC limit, sequences can be conveniently embedded into the total regulatory input space, i.e., the 2D space where each sequence is represented by a pair of values $(\rho_{\texttt{ON}}, \rho_{\texttt{OFF}})$. As a result, sequence distributions can be visualized as point clouds on a plane, where they are seen to have a peculiar discrete structure, characterized by "clouds" of high probability densities around certain $\rho$ values, as shown in Fig. F3. Here, we look into how distributions with such a peculiar shape arise.

In the limit of $c_{\max} \ll 1$ we can neglect the contributions from factors at concentration $c_{\min}$, so long as $n 10^{-R} \ll 1$. We also use the Taylor expansion of Eq. (1) around $c_{\max} = 0$ and approximate the contribution of a site in mismatch class $k$ as $c_{\max} e^{-\varepsilon k}$. A sufficiently high mismatch penalty $\varepsilon \geq 2$ ensures that most of the sites in the CRE have negligible occupancy. Therefore, we can approximate the total regulatory input of the cognate TF as the additive contribution of the $m_{k^*}$ number of strongest (lowest mismatched) sites with $k^*$ mismatches,

$$\rho_{\texttt{ON}} = m_{k^*} c_{\max} e^{-\varepsilon k^*} + \sum_{k > k^*} m_k c_{\max} e^{-\varepsilon k} \approx m_{k^*} c_{\max} e^{-\varepsilon k^*}, \quad [26]$$

where $m_k$ denotes the number of binding sites in mismatch class $k$, and $k^*$ is the lowest mismatch number found within the CRE.

According to Eq. (26), CREs that harbor $m_k^* = 1$ binding sites with $k^*$ mismatches cluster together and clusters corresponding to various $k^* = 0, 1, 2, \ldots$ are exponentially spaced in the regulatory input space, as we show in Fig. F3. As the binding site mutates into a higher mismatch class, the total occupancy of the CRE changes by a factor of $(1 - e^{-\varepsilon})$. This explains the large-scale "cloud" structure in the sequence distributions. Smaller clusters within these larger ones arise due to sequences that have more than one site with $k^*$ mismatches, also shown in Fig. F3.

We can describe the interaction between each noncognate TF and the CRE within the same approximation. However, clusters along the $\rho_{\mathtt{OFF}}$ axis can now arise from the combinatorial contribution of multiple noncognate factors. For example, sequences with one strong-enough site with $k^*$ mismatches for one of the noncognate TFs cluster around $\rho_{\mathtt{OFF}} \approx c_{\max} e^{-k^*}$. And sequences having 3 such sites for one noncognate TF or 1 such site for each of the 3 noncognate TFs clusters together around $\rho_{\mathtt{OFF}} \approx 3 c_{\max} e^{-k^*}$.

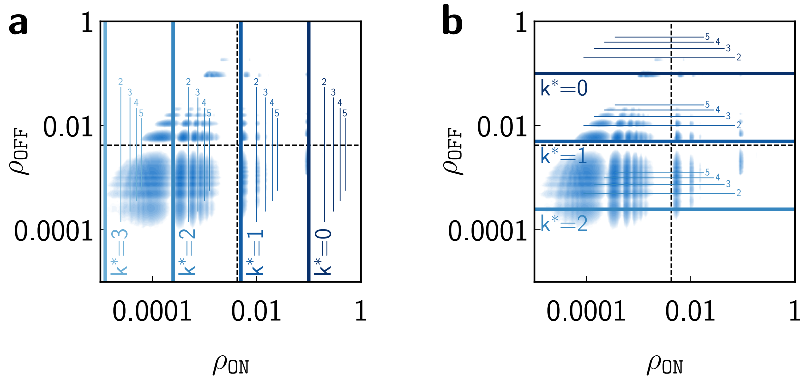

**Fig. F3. Neutral sequence distributions depicted in the space of total regulatory inputs.** Both plots show the same random sequence distribution in the $(\rho_{\mathtt{ON}}, \rho_{\mathtt{OFF}})$ space. In **(a)** we highlight the structure via approximation (see text) for $\rho_{\mathtt{ON}}$; in **(b)** we show the same for $\rho_{\mathtt{OFF}}$. The total occupancy $\rho_{\mathtt{ON}}$ is approximated by the dominant contribution of individual cognate sites, whereas $\rho_{\mathtt{OFF}}$ structure emerges from the combinatorial contribution of multiple noncognate sites. Thick lines represent the total occupancies arising from one site with $k^*$ mismatches. Thin lines show occupancies of multiples $(2, 3, 4, \ldots)$ of such sites. Underlying probability distributions (blue dots) are the true random (neutral) sequence distributions. Dashed black lines show the optimal activation threshold obtained by the information-theoretic calculation, explained in Section 2.A.1. At small $\rho$ (around $k^* \geq 2$ for $\rho_{\mathtt{ON}}$ and $k^* \geq 1$ for $\rho_{\mathtt{OFF}}$), the approximation is slightly off, with the thick lines falling into the gaps in the distribution, rather than onto the peaks. There, the background contribution from the entire CRE is comparable to the contribution from the dominant site with $k^*$ mismatches, yet sequences belonging to the next peak still most likely have a site with $k^*$ mismatches. This effect is more prominent – that is, happening at lower mismatch classes – for $\rho_{\mathtt{OFF}}$, since the background contribution of noncognate factors is typically $n$ times higher than for the cognate TF. The example distribution shown here is sampled at the following parameter values: $\ell = 8, \varepsilon = 3, L = 256, n = 3, R = 2, c_{\max} = 0.1, \delta = 10^7$.

**B. Optimally adapted CRE architecture in the STLC limit.** High mismatch penalty, $\varepsilon \geq 2$, ensures that the contribution of one "strong-enough" binding site will typically be greater than the contribution of any number of weak sites. In this case, we will always be able to find an activation threshold $(0 < \mu < 1)$ that allows for activation through a single strong-enough binding site but silences the gene when the CRE is bound only via weak sites. On such a GP map, CREs with one strong-enough cognate binding site and only weak noncognate sites are functional. Even the *cumulative* contribution of noncognate TFs will be close to zero on such maps, and so, naturally, will be the individual noncognate occupancies; importantly, all noncognate occupancies will be below the activation threshold individually, $\rho_2 < \mu, \rho_3 < \mu, \ldots \rho_{n+1} < \mu$, closely resembling the altered version of our model that we introduced for information-theoretic calculations in Section 2.B. Therefore, a threshold that allows for reliable gene activation through a single cognate binding site will be the optimal threshold, and the solution will correspond to the IT-efficient solution operating at the information lower bound, $\mathcal{D}_{\min}$.

How strong does the strong-enough single cognate site needs to be, i.e., what is the maximum number of mismatches that it can have? Its cognate occupancy has to be higher than the cumulative noncognate occupancy (i.e., total noncognate regulatory input) of the entire CRE. We can treat the CRE, as seen through the lens of noncognate factors, as a neutral (random) sequence, and approximate noncognate occupancies using a binomial distribution for mismatch counts along the CRE. Then, the mismatch count $k^*$ of the functional strong-enough site should fulfill:

$$c_{\max} e^{-\varepsilon k^*} > n(L - \ell + 1) c_{\max} \sum_k \mathrm{Binom}(\ell, k, 3/4) e^{-\varepsilon k}, \quad [27]$$

where we use equal background frequencies for each nucleotide, hence the 3/4 probability of finding a mismatch. We have $(L - \ell + 1)$ contributing site along the CRE for each of the $n$ noncognate TFs. We used Eq. (26) to describe the cognate total regulatory input of the CRE on the left-hand side.

The specific regulatory input only needs to be slightly higher than the total nonspecific regulatory input, since a sharp threshold with $\delta \to \infty$ can classify expression states even based on subtle differences. Therefore, from Eq. (27), the mismatch number of a strong-enough binding site is at most:

$$k^* \approx -\frac{1}{\varepsilon} \ln \left( n(L - \ell + 1) \rho_{\mathrm{neutral}} \right), \quad [28]$$

where we used the notation $\rho_{\text{neutral}} = \sum_k \text{Binom}(\ell, k, 3/4)e^{-\varepsilon k}$. We see that the predicted $k^*$ from this approximation coincides well with simulation results whenever $\varepsilon \gtrsim 2$, see Fig. F4.

Eq. (27) suggests that there is a maximum number of crosstalking TFs, $n_{\max}$, that a functional CRE can withstand in the one-strong-enough-site regulatory limit. We approximate $n_{\max}$ by assuming that the evolved functional strong-enough site actually has zero mismatches, $k^* = 0$,

$$n_{\max} < \frac{1}{(L - \ell + 1)\rho_{\text{neutral}}}. \qquad [29]$$

There exists a regime in which $n_{\max} < 1$, as shown in Fig. F4, meaning that the CRE cannot function via a single strong-enough binding site. CREs in this regime could still adapt by evolving multiple sites, but that happens at a higher information cost and thus takes much longer.

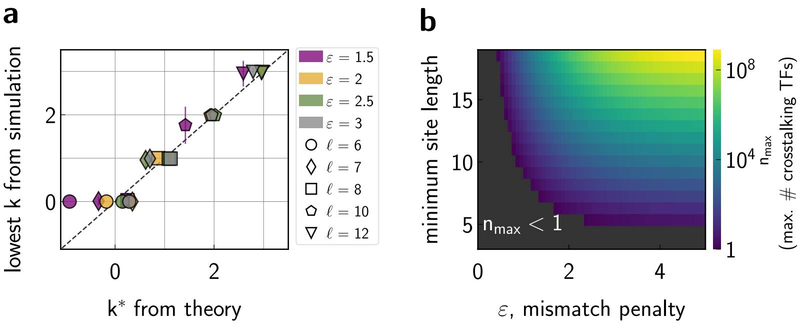

**Fig. F4. Optimally adapted CRE architecture and the limits of the IT-efficient regime.** **(a)** Minimal strength of the "strong-enough" binding site on the optimal genotype-phenotype map in the STLC limit, where reliable regulation can take place through a single cognate binding site. Shown is the strongest evolved site in the CRE in the dynamical simulations vs. the predicted mismatch from Eq. (28), for various $\varepsilon$ and $\ell$ parameters (legend). In cases where our prediction breaks down, at low $\varepsilon$ and whenever $k^* < 0$, adapted CREs are regulated through multiple cognate binding sites. Plotted lowest mismatch counts are averaged over 100 replicate simulations and are extracted when evolutionary trajectories first exceed the high fitness threshold ($F > 0.99$); Error bars are standard deviations. **(b)** Maximum number of crosstalking TFs that the CRE can effectively withstand in the efficient regime where regulation proceeds via one strong-enough cognate binding site. This is calculated for various BS lengths, $\ell$, and mismatch penalties, $\varepsilon$, at a fixed CRE length of $L = 256$ bp. Black region shows $(\ell, \varepsilon)$ parameter combinations that do not give rise to functional CRE in the single strong-enough site limit. As site length and mismatch penalty increase, $n_{\max}$ rapidly increases as well. Longer sites are less similar among themselves, and high mismatch penalties are an efficient way to suppress weak binding sites. Both of these effects act to decrease crosstalk (10). Parameters: (a-b) $L = 256, R = 2, c_{\max} = 0.1, \delta = 10^7$, (a) $n = 3, N = 100, \alpha = 1$.

We can also ask what happens outside of the STLC limit, when we allow concentrations to be high(er), but maintain a sharp, $\delta \to \infty$ , threshold. Here too, optimally adapted CREs implement regulation via one strong enough cognate binding site, as long as $\mu < 1$ holds. In this regime, the concentration at which the CRE with a single cognate site is functional is

$$c_{\max} > \frac{\mu}{1-\mu} e^{\varepsilon k^*}, \qquad [30]$$

where $k^*$ is the mismatch count of the site, and we assume that the strongest cognate site captures well the total occupancy in the ON environment, such that $\rho_{\texttt{ON}} = \dfrac{c_{\max}}{c_{\max} + e^{\varepsilon k^*}}$ and $\rho_{\texttt{ON}} > \mu$ holds for inducing expression in the $\delta \to \infty$ limit.

At the same time, the concentration must not be too high, because in that case the cumulative contribution of noncognate weak sites could induce erroneous expression in the OFF environment. The system can stay in the single-strong-enough-site regime only if the highest contributing noncognate sites have $k^* + 1$ mismatches, otherwise, there would be erroneous expression. We assume that the contribution of these weaker sites captures the total occupancy, $\rho_{\texttt{OFF}} = n \dfrac{c_{\max}}{c_{\max} + e^{\varepsilon(k^*+1)}}$, and that there is no ectopic expression, $\rho_{\texttt{OFF}} < \mu$. Then the concentration limit that must be satisfied reads:

$$c_{\max} < \frac{\mu}{n-\mu} e^{\varepsilon(k^*+1)}. \qquad [31]$$

The resulting concentration regime borders the regime where optimal solutions emerge in the simulations, as shown in Fig. F5.

**C. Adaptation rate estimates from information theory.** In the $\delta \to \infty$ limit, the genotype space consists of only two disjoint sets of functional and non-functional sequences. This makes it possible to estimate adaptation rates based on the fraction of functional sequences, $p(\mu)$, in this space. As $p(\mu)$ depends on the activation threshold, adaptation rates also depend on $\mu$. For clarity, we will use the notation $p = p(\mu)$ while keeping in mind the threshold dependency.

First, we calculate the average time it takes for a functional genotype to arise in the population. The population experiences incoming mutations at a rate of $NLu$, and the expected waiting time between two successive mutations is $1/(NLu)$. We assume that explored genotypes are independent and the probability of finding a functional genotype is $p$. Then, we can model the process with the geometric distribution, which gives us $1/p$ as the expected number of proposals until a functional genotype. Taken together, a functional genotype arises on average after $1/(pNLu)$ generations.

Second, we need to account for the fact that not every proposed mutation will be fixed in the population. In the $\delta \to \infty$ limit, where the fitness landscape is essentially flat, evolution is neutral. Under neutrality, only 1 out of $N$ mutations fix as

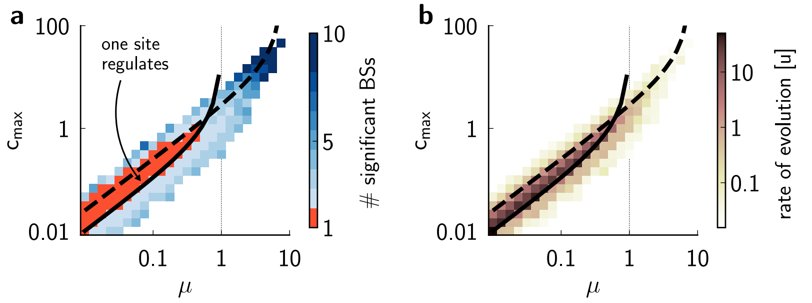

**Fig. F5. Optimal regulation regime in the $\delta \to \infty$ limit at higher concentrations. (a)** Average number of significant binding sites for the cognate factor in the adapted CRE (heatmap). Significant binding sites are defined as the strongest sites that cumulatively account for more than 90% of the total regulatory input, $\rho_{\mathtt{ON}}$. Solid black line shows $c_{\max}$ as a function of $\mu$, according to Eq. (30) at $k^* = 0$. This is the minimal concentration that permits the optimal regulation regime with a single cognate site. This curve diverges at $\mu = 1$, as the upper limit on the site contribution is 1, and at higher $\mu > 1$ values, one site is never enough to regulate even at concentrations when it is fully occupied. Concentrations that are too low lead to a suboptimal regime, in which CREs function through multiple cognate sites. Dashed black line shows $c_{\max}$ as a function of $\mu$, according to Eq. (31) at $k^* = 0$. This is the maximum concentration at which noncognate binding is still sufficiently low to permit reliable regulation. **(b)** Adaptation rates (heatmap) averaged over 100 evolutionary simulation replicates, as a function of $(c_{\max}, \mu)$ plane. Two limits from (a) are replotted in solid and dashed black lines. The limits delineate the optimal regime at which adaptive evolution is fastest and where evolved solutions feature a single significant cognate binding site. Here, the motif length is $\ell = 6$ bp and adapted CREs harbor a cognate site with, typically, 0 or 1 mismatches. Parameters: $L = 256, n = 8, \varepsilon = 3, \ell = 6, R = 2, c_{\max} = 0.1, \delta = 10^7$.

mutations spread through the population due to genetic drift rather than under selective pressure. Then it takes on average $N/p$ proposed mutations or $1/(pLu)$ generations to fix a functional genotype in the population. Therefore, the rate of adaptation is $pLu$, or, in terms of the information measure, $2^{-\mathcal{D}}Lu$. This matches very well the adaptation rates observed in the dynamical simulations, see Fig. S3.

Importantly, adaptation rates in this limit do not depend on population size or selection strength, $\alpha$; we expect and show (in the main paper as well as below) that this independence breaks down when we transition out of the $\delta \to \infty$ limit.

### 4. Detailed analyses in the realistic (smooth activation threshold) regime

In this section, we relax the sharp activation threshold assumption and use a finite value for $\delta$ instead. In the main paper, we focused our analyses on a particular pick of $\delta = 4$. Figs. F8, F9, F10 extend these analyses to other $\delta$ values, as summarized in SI Fig. S8.

A smooth threshold allows for a nonzero level of expression induced by the CRE, even if no factor is binding, which is often called "leaky" expression. This is the expression level induced at zero total occupancy. Leaky expression puts a limit on optimizing $\mu$, since we can no longer choose arbitrarily low threshold values if we look for fit CRE sequences, since the baseline expression will become too high in the `OFF` environment, rendering the CRE non-functional.

Similarly to the $\delta \to \infty$ limit, concentrations that are too high also lead to nonfunctional CREs, as they increase the crosstalk pressure by allowing high occupancies at weaker sites for noncognate factors. For $\mu$ values that are too high, as before, the CRE must still evolve multiple strong sites to induce expression, which is slow. Therefore, in this regime with finite $\delta$, a new, non-trivial optimum emerges at intermediate $\mu$ values, with a matching optimal concentration level, where both optimal parameters depend on the site length $\ell$ and the mismatch penalty $\varepsilon$.

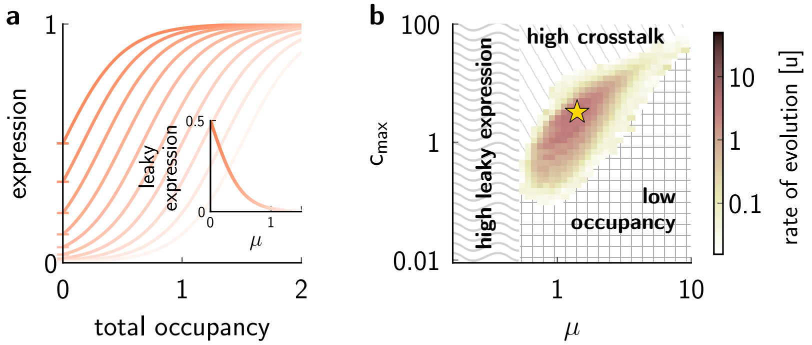

**Fig. F6. Parameter regimes where adaptation is impossible or proceeds exceedingly slowly. (a)** Genotype-phenotype maps at various $\mu$ parameters (orange shade). Small $\mu$ values (darker orange color) lead to an increased level of leaky expression (marks on the y-axis, inset). **(b)** Adaptation rate (colorbar) as a function of $c_{\max}$ and $\mu$. Optimal parameter combination that maximizes the adaptation rate is marked with a star. Adaptation is impossible in the "high leaky expression" regime. In the "high crosstalk" regime, adaptation is hindered by the noncognate factors that induce erroneous expression at high TF concentrations; selecting against such sites strongly slows down adaptation. In the "low occupancy" regime, the total CRE occupancy is insufficient to induce expression in the `ON` environment: here, concentrations are either too low (resulting in low site occupancy) or concentrations are high, but the activation threshold is too high, necessitating regulation through many strong sites, which do not emerge during the simulated evolution time. Parameters: $n = 8, \ell = 6, \varepsilon = 3, N = 100, \alpha = 1, \delta = 4, R = 2$.

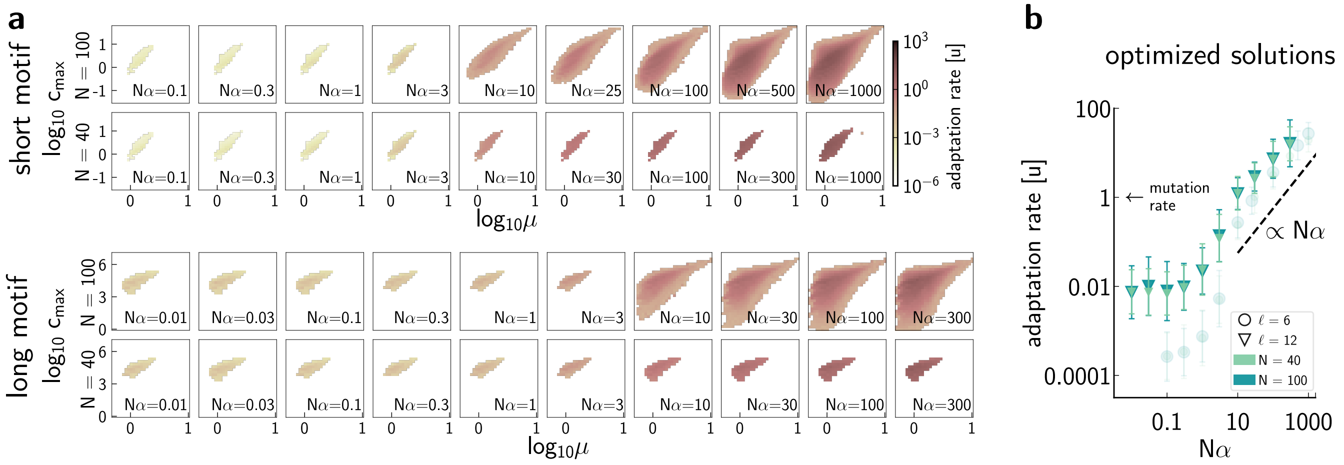

**Fig. F7. Adaptation rates with a smooth activation threshold depend on selection strength. (a)** Adaptation rates (colorbar) as a function of $(\mu, c_{\max})$, at various selection strengths $N\alpha$ (indicated in the subplots), for two motif lengths (upper row, $\ell = 6$; lower row, $\ell = 12$). Simulations at low $N\alpha$ ($N\alpha \leq 3$ for a short motif and $N\alpha \leq 1$ for a long motif) take a long time to finish. To save time, we only scan $(\mu, c_{\max})$ values around the expected optimum whenever possible; therefore, the subplots with smaller scanned regions do not report on the full space of possible evolutionary outcomes. Nevertheless, we ensure that even within the small scans, optimal solutions are locally surrounded by suboptimal ones. At least at the optimum, all replicates finish successfully, i.e., all 100 simulation replicates exceed the high fitness threshold, $F_{\text{thr}}(\alpha) = 0.99^\alpha$ (see text). To ensure this, simulations must run for a long time, up to $T_{\text{sim}} = 10000\ \frac{1}{u}$, which is typically $\sim 256$ million Monte Carlo steps. **(b)** Each plot symbol represents the adaptation rate at optimal $c_{\max}$ and $\mu$ in a system with long binding sites. Solid triangles are for $\ell = 12$; transparent circles for $\ell = 6$. Error bars show standard deviations over 100 replicates. In the regime where the fitness landscape is essentially flat (low $N\alpha$), adaptation is faster for a longer motif than for a shorter one. Optimal CREs with long binding sites have functional sites that contain a few mismatches (which is favorable so long as concentrations can be adjusted, as here, and take very high values, so that even with mismatches, the binding sites can be strongly occupied). In contrast, CREs regulated through short sites contain consensus or close to consensus sites at the optimum. As a consequence, the sequence space of possible functional long sites (which can contain mismatches) is larger the functional space of short sites that cannot contain mismatches, i.e., both scenarios require a similar total number of matches for functional regulation, but the long site scenario is entropically favored. Therefore, and perhaps counter-intuitively, adaptation is generally faster for a system that utilizes long binding sites. Note that this argument only holds when the maximal TF concentrations can be evolutionarily adjusted, as here, without bound. Parameters: $\delta = 4$, $L = 256$, $R = 2$, $\ell = 6, 12$.

In the smooth activation threshold regime, we simulated CRE adaptation in a fixed evolutionary setting with strong selection (Section 4.A), as well as in various evolutionary settings over a range of selection strengths (Section 4.B).

**A. Evolutionary outcomes with strong selection.** First, we fix evolutionary parameters such that selection is strong, e.g., $N\alpha \gg 1$; we typically use $N = 100, \alpha = 1$. We vary regulatory parameters $n, \ell, \varepsilon$ by choosing a representative pair of low and high values for each parameter, i.e., $\ell = \{6, 12\}, \varepsilon = \{1.5 \text{ or } 2, 3\}, n = \{3, 8\}$. Then, for each of these 8 parameter combinations, we run evolutionary simulations densely sampling the entire $(\mu, c_{\max})$ plane. From 100 replicate simulation trajectories at each $(\mu, c_{\max})$ combination, we assess which parameter set results in the fastest adaptation, thus identifying the optimal GP map.

At the identified optima, we then collect the adapted CRE sequences and examine their evolved structure, in terms of the number and strength of the binding sites they harbor. We find that optimal GP maps naturally lead to functional CREs that now contain not a single, but multiple cognate BSs. The number of sites depends on $n, \ell, \varepsilon$, as explored in detail in Figs. S4 and S5.

**B. Selection strength influences adaptation rates.** In the strong selection regime, the rate of evolution is set by selection, whose strength scales with the product $N\alpha$. In contrast, the population size sets the strength of evolutionary drift. Because the fixation probability of neutral mutations is $1/N$, higher $N$ results in stronger selection against neutral and deleterious mutations. Parameter $\alpha$ determines the steepness of the fitness function: higher $\alpha$ increases the fitness effects of small mutations, thereby increasing sensitivity for deleterious and beneficial mutations, making selection more effective. If $N\alpha$ is small, selection becomes weak and drift is the dominant evolutionary force on an essentially flat fitness landscape. However, the flatness here occurs at high fitness values, making this regime different from the STLC limit, where the flat fitness landscape occurs at low fitness values.

To assess functional CREs adapted at various strengths of selection ($\alpha$ values), we define an $\alpha$-dependent high fitness threshold, $F_{\text{thr}}(\alpha) = 0.99^\alpha$. This ensures that CREs exceeding this threshold reach the same precision in gene expression. Our reasoning is as follows: we have already arbitrarily chosen to use $F_{\text{thr}}(\alpha = 1) = 0.99$ at $\alpha = 1$. This corresponds to a particular precision, $x^*$, for the underlying expression pattern:

$$F_{\text{thr}}(\alpha = 1) = e^{-\frac{1}{2}\left((1-g_{\text{ON}})^2 + (0-g_{\text{OFF}})^2\right)} = e^{-x^*}$$
$$x^* = -\ln F_{\text{thr}}(\alpha = 1). \qquad [32]$$

We require the same precision regardless of the specific choice of $\alpha$, and write:

$$F_{\text{thr}}(\alpha) = e^{-\alpha x^*} = F_{\text{thr}}(\alpha = 1)^\alpha. \qquad [33]$$

Evolutionary outcomes on fitness landscapes at different $N\alpha$ values are shown in Fig. F7.

**C. Interplay between the GP map and the phenotype-fitness map.** As a last exploration of the more realistic model, we run evolutionary simulations as well as information calculations to gauge the effect of the steepness of the GP map (set by the regulatory parameter $\delta$) and the steepness of the fitness map (set by the evolutionary parameter $\alpha$).

First, we focus on the GP map effect and explore four values $\delta = \{1, 2, 3, 4\}$ while fixing $\alpha = 1$. We expect a higher $\delta$ to facilitate faster adaptation, since on steeper GP maps the effective selection coefficient for an identical mutation would be higher, thereby speeding up adaptive evolution. We validate this intuition in Figs. F8 and F9. Simultaneously, because overspecification presents a greater issue on smoother GP maps, adaptation will have to accumulate more bits of information to counteract its effect at low $\delta$. We show this effect on the fitness-level information in Fig. F10 and on the mismatch-level information in Fig. S8.

Second, we assess how the steepness of the fitness function influences our results. We run evolutionary simulations over a wide range of $\delta$ and $N\alpha$ parameters, depicted in Fig. F11. For each of these regulatory / evolutionary parameter combinations, we optimize over the $(\mu, c_{\max})$ plane to obtain the fastest-adapting solutions. We expect that higher $\alpha$ also facilitates faster adaptation, for the qualitatively similar reasons that led us to expect faster adaptation with higher $\delta$. We report on these results in Fig. S8. One salient observation is that the speed-up due to higher $\alpha$ is higher at lower $\delta$: this is likely because at high $\delta$, more of the sequence space is dominated by drift due to the vanishing selection gradient there.

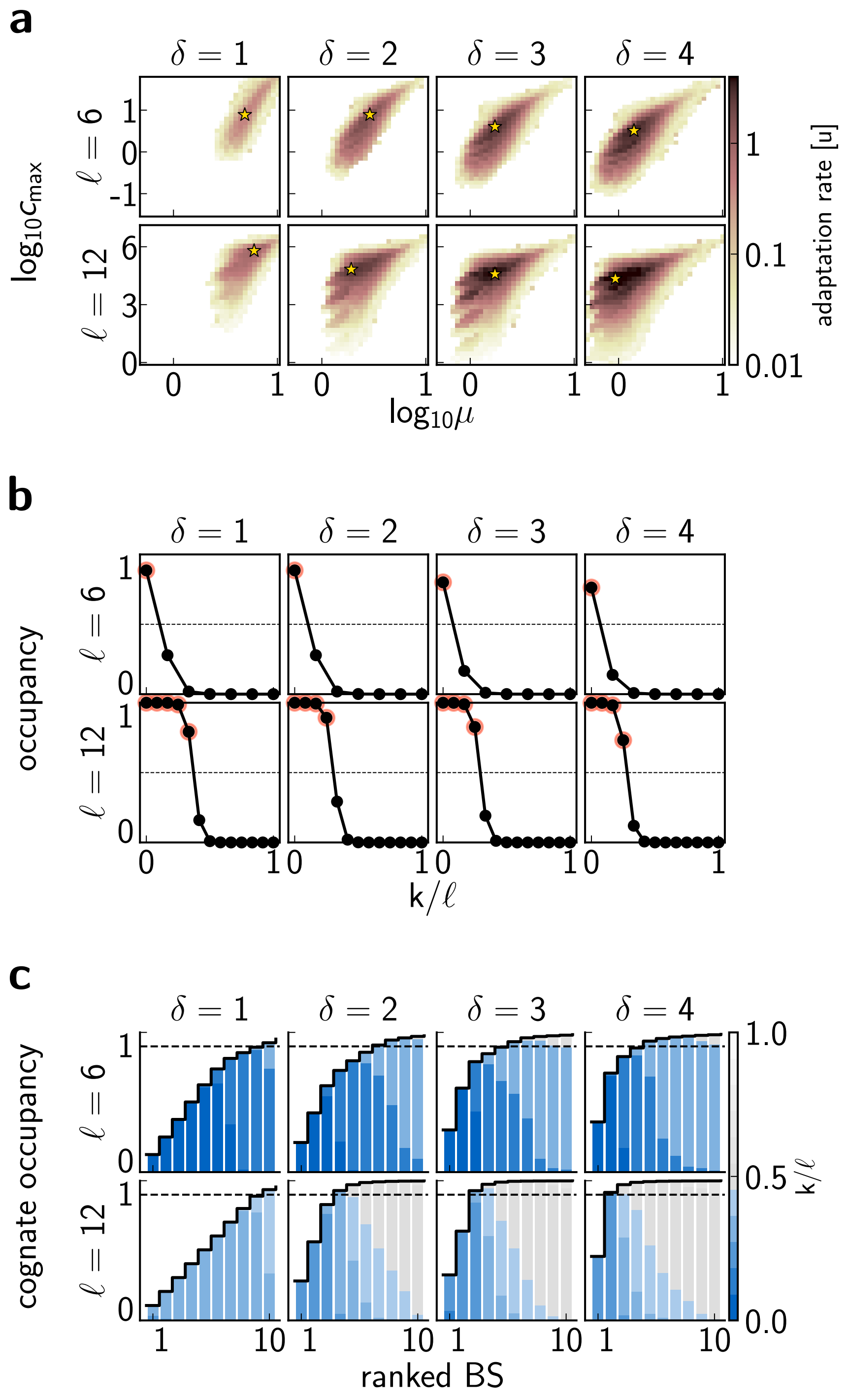

**Fig. F8.** **Evolutionary outcomes depend on activation threshold steepness $\delta$.** We analyzed this regime by comparing four GP maps that differ in their steepness. We display results for $\delta = 1, 2, 3, 4$ (columns) for two motif lengths, $\ell = \{6 \text{ vs. } 12\}$ bp (rows). 100 replicate simulations were run at every $(l, \delta, \mu, c_{\max})$ combination shown in **a**. **(a)** CRE adaptation rates (colorbar) as a function of $(\mu, c_{\max})$ for different parameter combinations. Adaptation proceeds faster on steeper GP maps (higher $\delta$), evolvable solutions cover a wider parameter range, and are slightly shifted across different $\delta$ values. Optimal $(\mu, c_{\max})$ parameters are denoted with a star. **(b)** Occupancy curves as a function of normalized mismatch class, $k/\ell$, plotted at the optimal value of regulatory parameters (denoted by a star in a). Red circles highlight strong sites that have $> 0.5$ occupancy at optimal concentration $c^*_{\max}$. The dashed line shows $0.5$ occupancy for ease of visualization. The curves are qualitatively similar across $\delta$ for short sites, with only the consensus site reaching high occupancy; in contrast, for long binding sites, the occupancy curves saturate. **(c)** Fractional contributions to the cognate regulatory input, $\rho_s$, at the optimum $(\mu^*, c^*_{\max})$. Evolved BSs are ranked by their contribution (x-axis), color (bar at right) shows the normalized match to the TF consensus (dark blue = full match, gray = complete mismatch), thick black line = cumulative input, dashed black = 90% threshold. A smoother GP (lower $\delta$) necessitates the emergence of multiple strong sites, as CREs on these maps need to be highly specified to reach high fitness. Parameters: $L = 256, n = 8, \varepsilon = 3, R = 2, \alpha = 1, N = 100$.

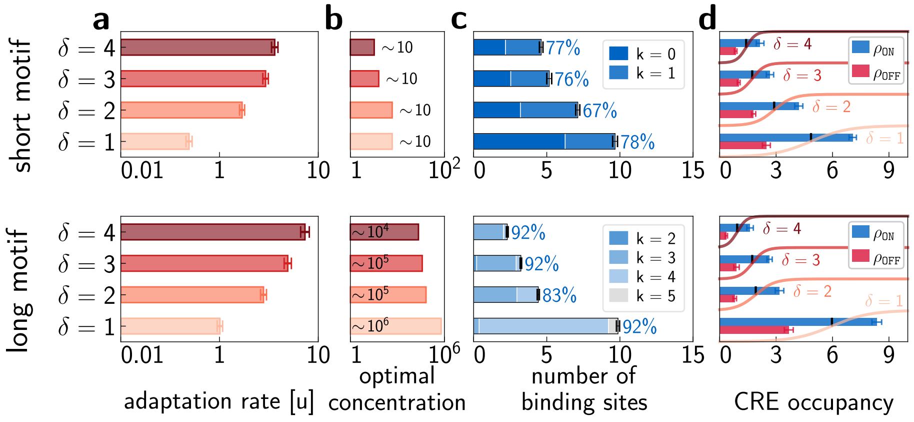

**Fig. F9. Optimal evolutionary outcomes for various activation threshold steepness values $\delta$.** Optimal solutions with short ($\ell = 6$ bp, top row) and long ($\ell = 12$ bp, bottom row) binding sites are shown on four different GP maps with varying steepness, $\delta = \{1, 2, 3, 4\}$. **(a)** CRE adaptation rates at the optimal $(\mu^*, c^*_{\max})$ parameters. Steeper GP maps (higher $\delta$) and longer motifs foster faster adaptation. **(b)** Optimal TF concentrations, $c^*_{\max}$. CREs with longer binding sites emerge more quickly but require much higher concentrations so as to saturate their binding sites even with mismatches, as shown in Fig. F8. **(c)** Structural details of the evolved CREs for different parameters. Bars show the number of BSs contributing significantly to total regulatory input, broken down by the number of mismatches (bluish hue; legend). Percentages show the fraction of total regulatory input accounted for by BSs with occupancy $> 0.5$ at the optimally chosen $c_{\max}$. Adapted CREs are enriched in strong sites over all GP maps explored here, and CREs adapted on smoother GP maps contain more of these. **(d)** Absolute values for the total CRE occupancy (regulatory input) in the `ON` and `OFF` environment, $\rho_{\texttt{ON}}$ (blue) and $\rho_{\texttt{OFF}}$ (red). Small black markers indicate the optimal activation threshold, $\mu^*$. We indicate the optimal activation nonlinearity (sigmoid) for each $\delta$ over the bars, showing that regulatory input $\rho_{\texttt{OFF}}$ and $\rho_{\texttt{ON}}$ result in the correct gene expression pattern, close to $\{0, 1\}$. $\delta = 1$ requires the highest optimal $\mu^*$ to avoid leaky expression; it also necessitates high CRE occupancy achieved by multiple strong sites, ensuring that expression in the `ON` environment is well specified. Bars show averages, and error bars represent standard error of the mean, over 100 replicates on **[a,c,d]**. Parameters: $L = 256, n = 8, \varepsilon = 3, R = 2, \alpha = 1, N = 100$.

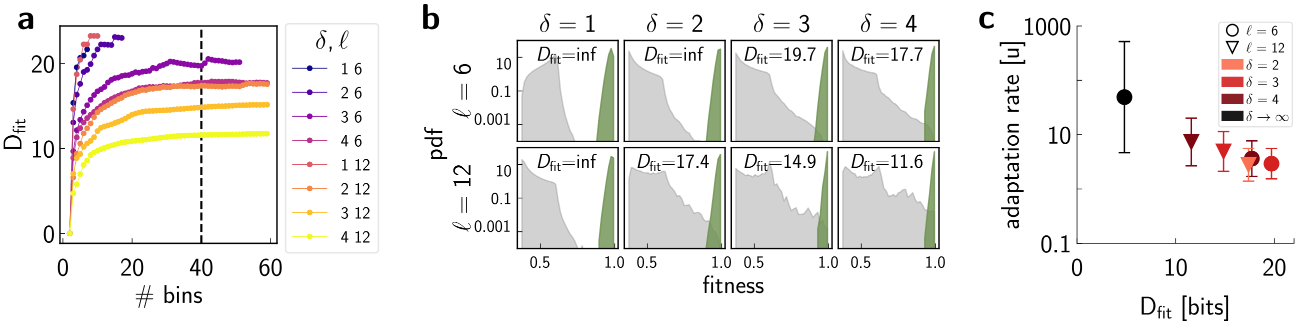

**Fig. F10. Fitness level information for optimal solutions with different activation threshold steepness values $\delta$.** **(a)** $D_{\text{fit}}$ dependence on the choice of binning used for information-theoretic estimations. $\delta, \ell$ parameter combinations are denoted by color (legend). On the x-axis, we show the number of bins over the possible fitness interval $(e^{-1}, 1)$ at $\alpha = 1$. We use equally sized bins, and choose 40 bins to calculate the $D_{\text{fit}}$ measure (indicated by the black dashed line). **(b)** Neutral (gray) and adapted (green) fitness distributions at optimal $(\mu^*, c^*_{\max})$ values for various $\delta, \ell$ parameters. $D_{\text{fit}}$ is indicated in each subplot separately. For the neutral distributions, we sampled $10^8$ random sequences. In three cases, the neutral distribution is undersampled, and we could not obtain $D_{\text{fit}}$, indicated by the "inf" in the corresponding plots. **(c)** Adaptation rate is inversely correlated with the fitness level information across various $\delta, \ell$ parameters. Black dot indicates the IT efficient solution in the STLC limit at $\delta \to \infty$ and $\ell = 6$. Adaptation is faster if the information that must be accumulated by selection is lower, and that is facilitated by steeper GP maps (higher $\delta$ leading to less overspecification) and longer sites. Parameters: $L = 256, n = 8, \varepsilon = 3, R = 2, \alpha = 1, N = 100$.

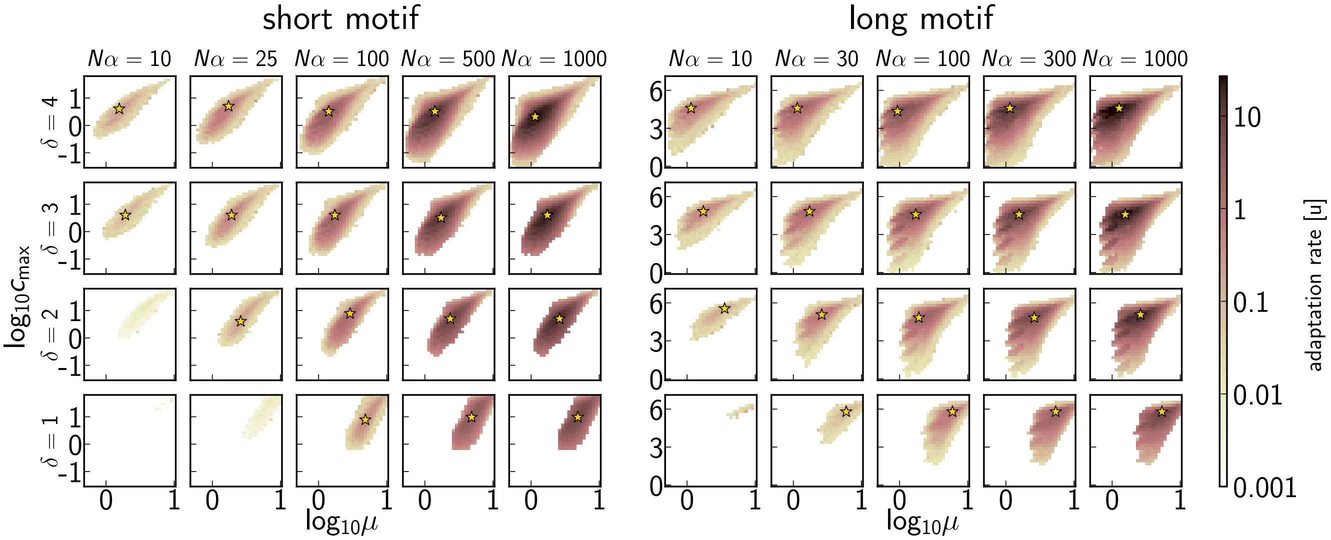

**Fig. F11. Adaptation rates depend on selection strength and on the activation threshold steepness.** CRE adaptation rates (color) as a function of $(\mu, c_{\max})$, for various values of evolutionary parameter $N\alpha$ that sets the selection strength (columns), and different regulatory parameters ($\delta$, rows; motif length $\ell = (6$ vs. $12)$, left vs. right subpanel, respectively). Simulations at low $\delta$ and insufficient $N\alpha$ take a long time to finish, even though we only sampled relatively high $N\alpha$ values. Optimal solutions are indicated with a star whenever all 100 simulation replicates finished successfully by reaching the high fitness threshold $F_{\text{thr}}(\alpha) = 0.99^{\alpha}$. We typically let the simulation run up to $T_{\text{sim}} = 100\ \frac{1}{u}$, except for the cases at $(\ell, \delta, N\alpha) = (6, 1, 10), (6, 1, 25), (6, 2, 10)$ where we simulated up to $T_{\text{sim}} = 1000\ \frac{1}{u}$ yet never reached 100% success rate at the optimum (we report $3, 82, 95$ successful replicates out of 100 for these cases, respectively). We still report on the last two scenarios at $(\ell, \delta, N\alpha) = (6, 1, 25)$ and $(6, 2, 10)$ in Fig. S8, with a caveat that their adaptation rates are slightly overestimated. In the case of $(\ell, \delta, N\alpha) = (12, 1, 10)$, trajectories also did not finish successfully; as we only have 8% success rate at the optimum, we omit this parameter combination from other analyses. Parameters: $L = 256, n = 8, \varepsilon = 3, R = 2, \ell = 6, 12$.

### References

1. U Gerland, JD Moroz, T Hwa, Physical constraints and functional characteristics of transcription factor–DNA interaction. *Proc. Natl. Acad. Sci.* **99**, 12015–12020 (2002) Publisher: Proceedings of the National Academy of Sciences.
2. JB Kinney, A Murugan, CG Callan, EC Cox, Using deep sequencing to characterize the biophysical mechanism of a transcriptional regulatory sequence. *Proc. Natl. Acad. Sci.* **107**, 9158–9163 (2010) Publisher: Proceedings of the National Academy of Sciences.
3. J Berg, S Willmann, M Lässig, Adaptive evolution of transcription factor binding sites. *BMC Evol. Biol.* **4**, 42 (2004).
4. M Lässig, From biophysics to evolutionary genetics: statistical aspects of gene regulation. *BMC Bioinforma.* **8**, S7 (2007).
5. OG Berg, PH von Hippel, Selection of DNA binding sites by regulatory proteins: Statistical-mechanical theory and application to operators and promoters. *J. Mol. Biol.* **193**, 723–743 (1987).
6. L Bintu, et al., Transcriptional regulation by the numbers: models. *Curr. Opin. Genet. & Dev.* **15**, 116–124 (2005).
7. L Bintu, et al., Transcriptional regulation by the numbers: applications. *Curr. Opin. Genet. & Dev.* **15**, 125–135 (2005).
8. R Grah, B Zoller, G Tkačik, , et al., Nonequilibrium models of optimal enhancer function. *Proc. Natl. Acad. Sci.* **117**, 31614–31622 (2020).
9. B Zoller, T Gregor, G Tkačik, Eukaryotic gene regulation at equilibrium, or non? *Curr. Opin. Syst. Biol.* **31**, 100435 (2022).
10. T Friedlander, R Prizak, CC Guet, NH Barton, G Tkačik, Intrinsic limits to gene regulation by global crosstalk. *Nat Commun* **7**, 12307 (2016) Number: 1 Publisher: Nature Publishing Group.
11. ML Perkins, J Crocker, G Tkačik, Chromatin enables precise and scalable gene regulation with factors of limited specificity. *Proc. Natl. Acad. Sci.* **122**, e2411887121 (2025).
12. M Kimura, On the Probability of Fixation of Mutant Genes in a Population. *Genetics* **47**, 713–719 (1962).
13. DT Gillespie, Exact stochastic simulation of coupled chemical reactions. *The journal physical chemistry* **81**, 2340–2361 (1977).
14. T Friedlander, R Prizak, NH Barton, G Tkačik, Evolution of new regulatory functions on biophysically realistic fitness landscapes. *Nat Commun* **8**, 216 (2017) Bandiera_abtest: a Cc_license_type: cc_by Cg_type: Nature Research Journals Number: 1 Primary_atype: Research Publisher: Nature Publishing Group Subject_term: Biological physics;Evolutionary theory;Molecular evolution;Regulatory networks Subject_term_id: biological-physics;evolutionary-theory;molecular-evolution;regulatory-networks.
15. M Hledík, N Barton, G Tkačik, Accumulation and maintenance of information in evolution. *Proc. Natl. Acad. Sci.* **119**, e2123152119 (2022) Publisher: Proceedings of the National Academy of Sciences.
16. A Wagner, Information theory, evolutionary innovations and evolvability. *Philos. Transactions Royal Soc. B: Biol. Sci.* **372**, 20160416 (2017) Publisher: Royal Society.

### 5. Supplementary figures

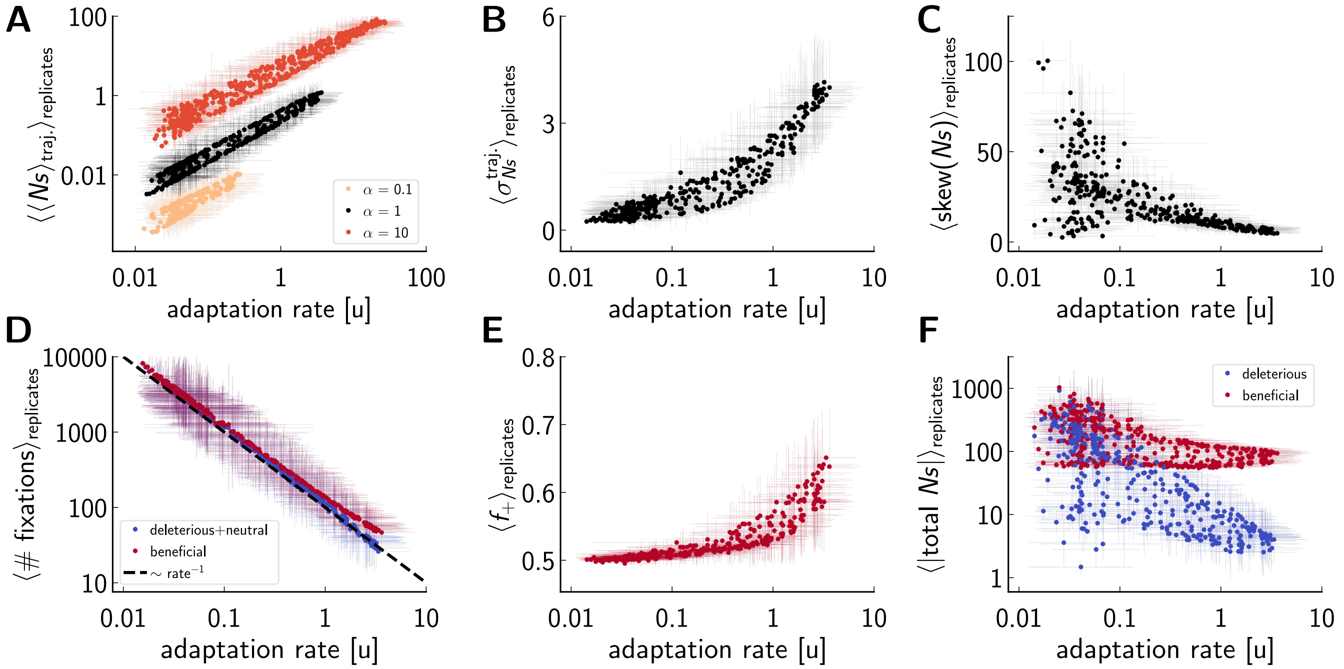

**Fig. S1.** **Selection strength and other fixation statistics for CRE evolution on GP maps with smooth activation threshold.** Effective selection strength of fixed mutations, $Ns$, across adaptation trajectories and replicates at specific regulatory parameters: $n = 8, \ell = 6, \varepsilon = 3$. We measure $Ns = N(F_{\mathrm{new}} - F_{\mathrm{old}})/F_{\mathrm{old}}$ only when a fixation/substitution event happens; therefore, this quantity is not the traditionally used $Ns$ of population genetics, where $s$ stands for the fitness advantage of each arising mutation. Each dot represents an entire GP map defined by a combination of regulatory parameters $(\mu, c_{\mathrm{max}})$ (with other parameters held fixed to values reported in this caption). Adaptation on these various GP maps unfolds at different rates (see the adaptation rate landscape on Fig. S4A, upper right corner) shown on the horizontal axis; different panels report on various quantities (vertical axes) as a function of adaptation rate. For each GP map, we run 100 replicates used for statistics here. On a small fraction of highly suboptimal maps, some of the replicates do not reach the fitness threshold during the simulation time; such replicates are omitted from the statistics. **(A)** Selection strength of fixed mutations, $Ns$ averaged over the evolutionary trajectory duration and over replicates, as a function of the adaptation rate. Different $\alpha$ (color) directly modulates $s$ in our model and thus systematically vertically shifts the points. Optimal GP maps that support higher adaptation rates feature mutations with higher fitness differences. **(B)** Average (across replicates) of the standard deviation (across trajectories) of $Ns$. On optimal GP maps, $Ns$ has a wider distribution, compared to the suboptimal GP maps (low adaptation rate), where many $Ns$ values are close to zero, resulting in a tighter distribution. **(C)** Average (across replicates) of the skewness (across trajectories) of $Ns$. **(D)** Average (across replicates) of the number of fixations per CRE adaptation trajectory. Blue dots show deleterious and neutral fixations ($s \leq 0$), red dots show beneficial or beneficial fixations ($s > 0$). Faster adaptation happens on landscapes that require a smaller number of fixations and where there is an excess of beneficial mutations. **(E)** Average (over replicates) fraction of beneficial fixations. For the worst GP maps, evolution proceeds through roughly an equal number of deleterious and beneficial fixations; for optimal GP maps, $\sim 65\%$ of all fixations are beneficial. **(F)** Absolute value of accumulated fitness effects, $Ns$, separated into beneficial (red) and deleterious (blue), per trajectory, averaged over replicates. Combined with panels (**D, E**), we see that optimal GP maps feature CRE evolution via fewer fixations that are more effective. Baseline parameters: $L = 256, n = 8, \ell = 6, \varepsilon = 3, R = 2, \delta = 4, \alpha = 1, N = 100$. Error bars show the standard deviations over replicates. On log-scaled axes, we report on means and standard deviations over $\log_{10}(\mathrm{quantity})$.

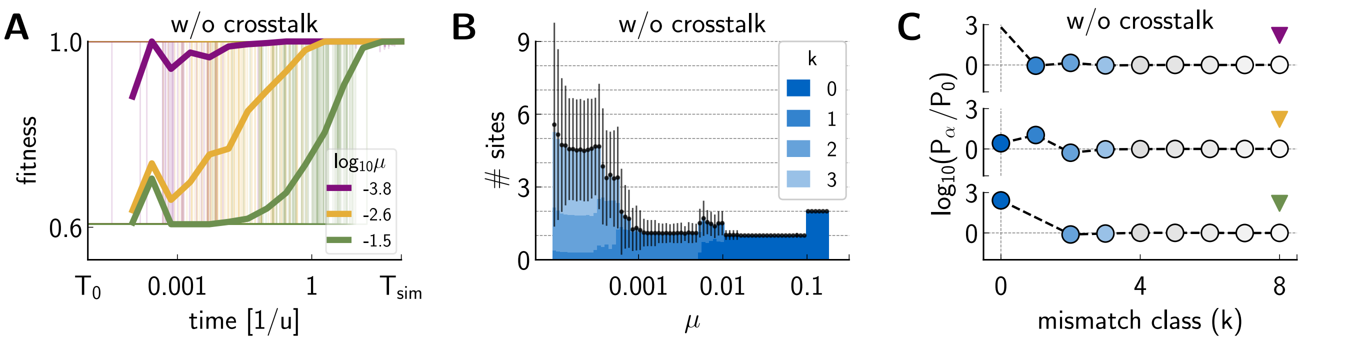

**Fig. S2. Control case results without crosstalk in the STLC limit. (A)** Replicate simulation fitness trajectories (thin lines, 100 replicates per $\mu$) and their mean (thick lines) at three activation threshold values $\mu$ (legend). Adaptation is fastest at low $\mu$ (purple). On average, as $\mu$ increases, the search on the flat fitness landscape takes longer. Simulations start at $T_0 = 10^{-5}$ and terminate at $T_{\text{sim}} = 80\frac{1}{u}$ with characteristic time scale of $1/(NL)$. **(B)** BS composition of evolved CREs as a function of $\mu$. Color (legend) indicates the average mismatch composition of BS that together contribute at least $90\%$ of the total cognate regulatory input; shown are average $\pm$ SD over replicates. As expected, adapted CREs differ from the crosstalk case in the low $\mu$ limit, where crosstalk has the highest impact; in this regime, fewer BSs with fewer mismatches evolve compared to the same scenario with crosstalk. **(C)** Ratio between adapted $(P_\alpha)$ and neutral $(P_0)$ mismatch distribution across the entire CRE for different $\mu$ (triangle color as in A), for cognate (blue) TFs. Positive values indicate enrichment of the corresponding BSs in evolved CREs. Missing markers at small $k$ indicate that no corresponding CRE sequence evolved across $100$ simulated replicates. Statistics are extracted when a replicate simulation run first exceeds fitness $F > 0.99$. Parameters: $L = 256, \ell = 8, n = 3, R = 2, c_{\max} = 0.1, \delta = 10^7, \varepsilon = 3, \alpha = 1, N = 100$.

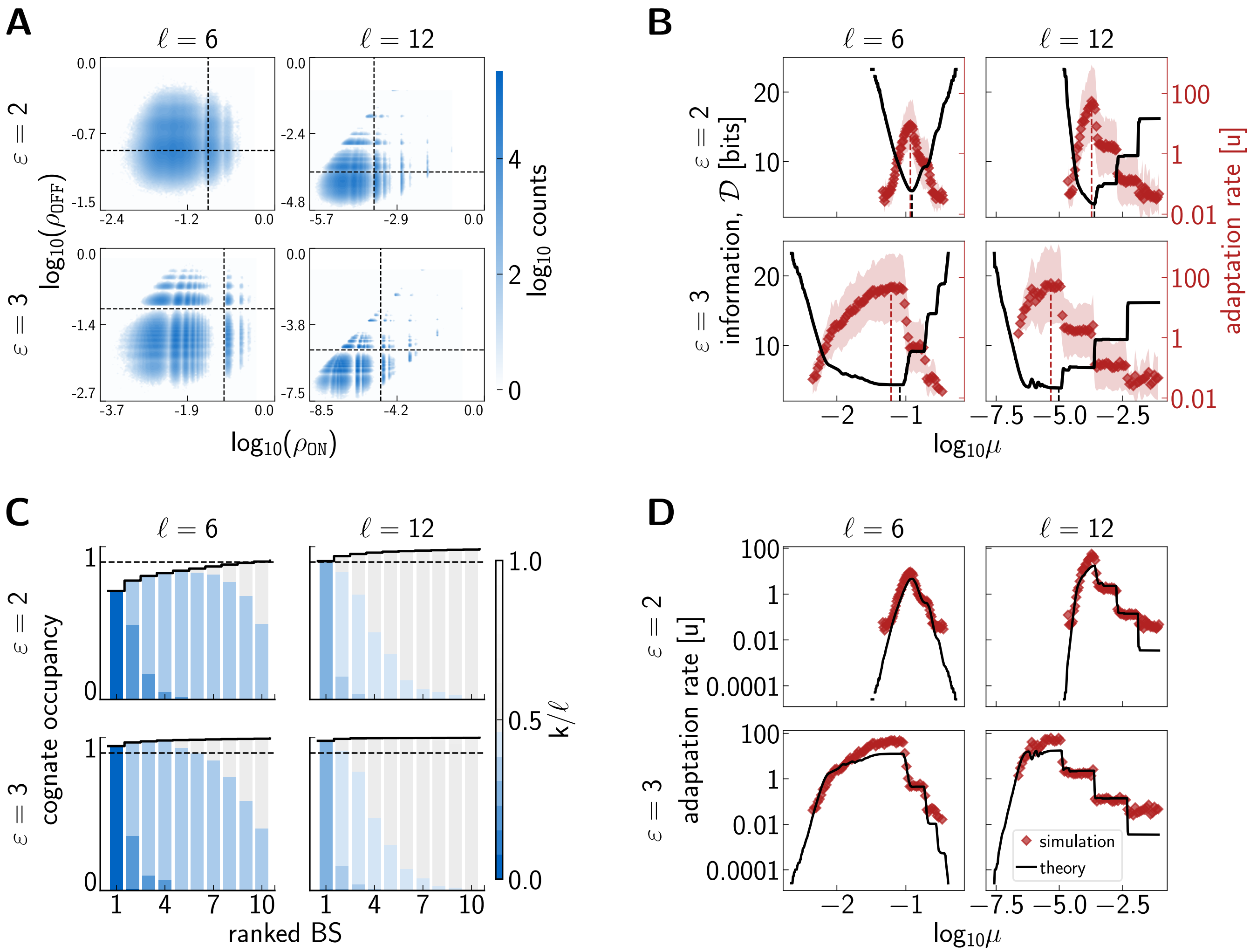

**Fig. S3.** **Detailed analysis of the STLC limit including information theoretic predictions.** We analyzed this regime over a wide range of parameters, for each combination of mismatch penalty and motif length from the sets of $\varepsilon = \{1.5, 2, 2.5, 3\}\ k_BT$ and $\ell = \{6, 7, 8, 10, 12\}$ bp. Here we report on a subset of these results. In each subpanel, we highlight results with short vs. long motifs ($\ell = 6$ bp vs. 12 bp, in columns) and a lower and higher mismatch penalty value ($\varepsilon = 2\,k_BT$ vs. $3\,k_BT$, in rows). **(A)** Monte Carlo sampling of random sequences depicted in the $(\rho_{\mathrm{ON}}, \rho_{\mathrm{OFF}})$ plane. Dashed black lines indicate optimal activation threshold, $\mu^*$; sequences falling to the lower right corner of the distributions (bounded by the dashed lines) are functional with respect to $\mu^*$. **(B)** Overlay of information curve, $\mathcal{D}$ (black, left axis, Monte Carlo over random sequences) and adaptation rate estimated via "optimize-to-adapt" approach (red, right axis, evolutionary dynamics simulations). Activation threshold that minimizes $\mathcal{D}$ (black dashed line) nearly coincides with activation threshold that maximizes adaptation rate (red dashed line). **(C)** Fractional contributions to cognate regulatory input, $\rho_s$, at optimal activation threshold (indicated by red dashed lines on **B**). Evolved BSs are ranked by contribution (x-axis), color (bar at right) shows the normalized match to TF consensus (dark blue = full match, gray = complete mismatch), thick black line = cumulative input, dashed black = 90% threshold. At $\varepsilon = 2$ multiple sites contribute at $\ell = 6$ in contrast to $\ell = 12$, however, at higher mismatch penalty ($\varepsilon = 3$), regulation goes through only one binding site for both long and short mismatches. **(D)** Adaptation rates from "optimize-to-adapt" simulations recapitulate rates derived from IT calculations (rate $\sim 2^{-\mathcal{D}}Lu$) over a wide range of activation thresholds. Baseline parameters: $L = 256, n = 3, R = 2, c_{\max} = 0.1, \delta = 10^7, \alpha = 1, N = 100.$

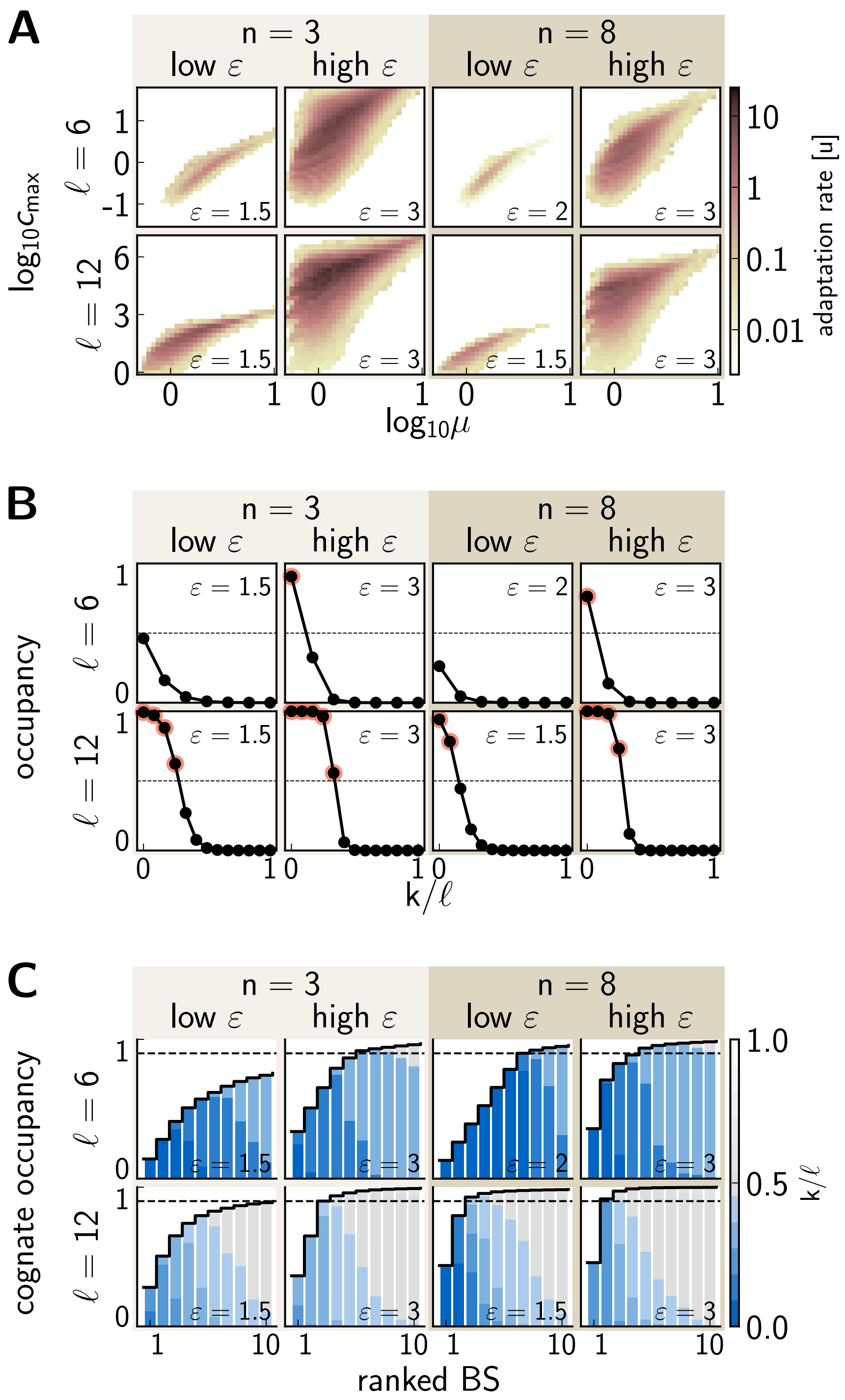

**Fig. S4.** **Evolutionary outcomes on GP maps with a smooth activation threshold.** Smooth activation threshold ($\delta = 4$) regime was explored by comparing and contrasting low vs high parameter values as follows: motif length, $\ell = \{6 \text{ vs. } 12\}$ bp (in rows); number of crosstalking TFs, $n = \{3 \text{ vs. } 8\}$ (light vs. dark background); mismatch penalty, $\varepsilon = \{1.5 \text{ or } 2 \text{ vs. } 3\}$ (in columns). Results for specific $(n, l, \varepsilon)$ parameter combinations are shown in each subfigure as averages over 100 replicates. With parameters $\ell = 6, n = 8, \varepsilon = 1.5$, evolutionary simulations have taken too long to complete, therefore, we report here on results with a slightly higher mismatch penalty value, $\varepsilon = 2$. **(A)** CRE adaptation rates (colorbar) as a function of $(\mu, c_{\text{max}})$, for different regulatory parameter combinations. High mismatch penalty and low number of noncognate factors cause minimal crosstalk, extending the feasible solution space in the $(\mu, c_{\text{max}})$ plane toward the upper left corner, parametrized by high concentrations and low threshold values (second column). In contrast, low $\varepsilon$ increases the crosstalk pressure, resulting in a much narrower solution space regardless of $n$ (columns 1 and 3). The last column shows the resulting solution space when high $n$ yields an increased crosstalk pressure, but it is counteracted by the crosstalk-reducing effect of high mismatch penalty $\varepsilon$. **(B)** TF occupancy curves for a site as a function of normalized mismatch class, $k/\ell$, computed for the optimal $c^*_{\text{max}}$ identified in (A). Red circles highlight strong sites that have $> 0.5$ occupancy (dashed horizontal line). At low mismatch penalty and short binding sites, even perfect (zero-mismatch) binding sites are not fully saturated and their occupancy falls below $0.5$. Even though they can contribute to gene activation, these sites are "weak" in the occupancy sense. **(C)** Fractional contributions to cognate regulatory input, $\rho_s$, at the optimum $(\mu^*, c^*_{\text{max}})$. Evolved BSs are ranked by contribution (x-axis), color (bar at right) shows the normalized match to TF consensus (dark blue = full match, gray = complete mismatch), thick black line = cumulative input, dashed black = 90% threshold. Short motif length facilitates the emergence of multiple significant binding sites, whereas a lower mismatch penalty results in sites with fewer mismatches. Increased crosstalk pressure from a higher number of noncognate TFs results in fewer but better specified binding sites. Baseline parameters: $L = 256, R = 2, \delta = 4, \alpha = 1, N = 100$.

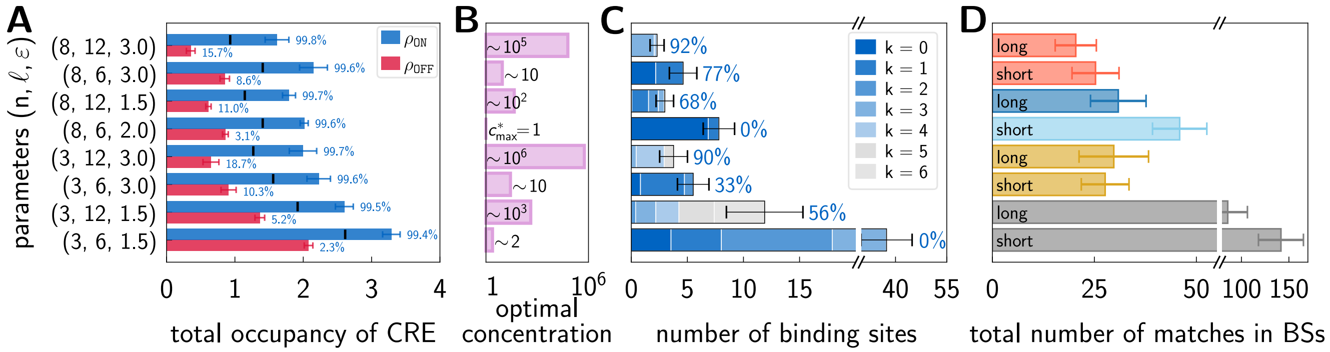

**Fig. S5. Properties of optimal solutions on realistic (smooth activation threshold) GP maps.** We show results for the same parameters $(n, \ell, \varepsilon)$ as in Fig. S4. **(A)** Absolute values of total occupancy in the ON and OFF environment, $\rho_{\mathtt{ON}}$ (blue) and $\rho_{\mathtt{OFF}}$ (red). Percentages show the fraction of the total contribution coming from the cognate TF. $\rho_{\mathtt{ON}}$ comes almost entirely from the specific contribution of the cognate TF. Cognate contribution is not negligible in the OFF environment, especially when regulation is implemented through long binding sites ($\ell = 12$ bp), since optimal concentrations must be high in these cases for rapid adaptation to take place. Little black markers indicate the optimal activation threshold, $\mu^*$. **(B)** Optimal maximal TF concentrations, $c^*_{\max}$ normalized to the reference case of $n = 8, \ell = 6, \varepsilon = 2$. **(C)** Structural details of adapted CREs for different regulatory parameters. Bars show the number of BSs contributing significantly to the total regulatory input, broken down by the number of mismatches (bluish hue; legend). Percentages show the fraction of total regulatory input accounted for by BSs with occupancy $> 0.5$ at the optimally chosen $c^*_{\max}$ for each scenario. In scenarios with short binding sites and low mismatch penalty, consensus sites do evolve, but they are only weakly occupied (indicated by 0%). **(D)** The total number of matches in significant BSs (contributing $\sim 90\%$ of regulatory input) averaged over replicates. Same colors represent the same $n, \ell, \varepsilon$ values. At the optimum, a similar number of loci needs to be specified across the CRE through both short and long binding sites at $\varepsilon = 3$, colored red and yellow. This leads to more short binding sites (see **C**) that are better matched to the cognate consensus, whereas mismatches are easily permitted in longer binding sites. Baseline parameters: $L = 256, R = 2, \delta = 4, \alpha = 1, N = 100$. On (**A, C, D**), we report on average values over 100 replicates, and error bars show standard deviations.

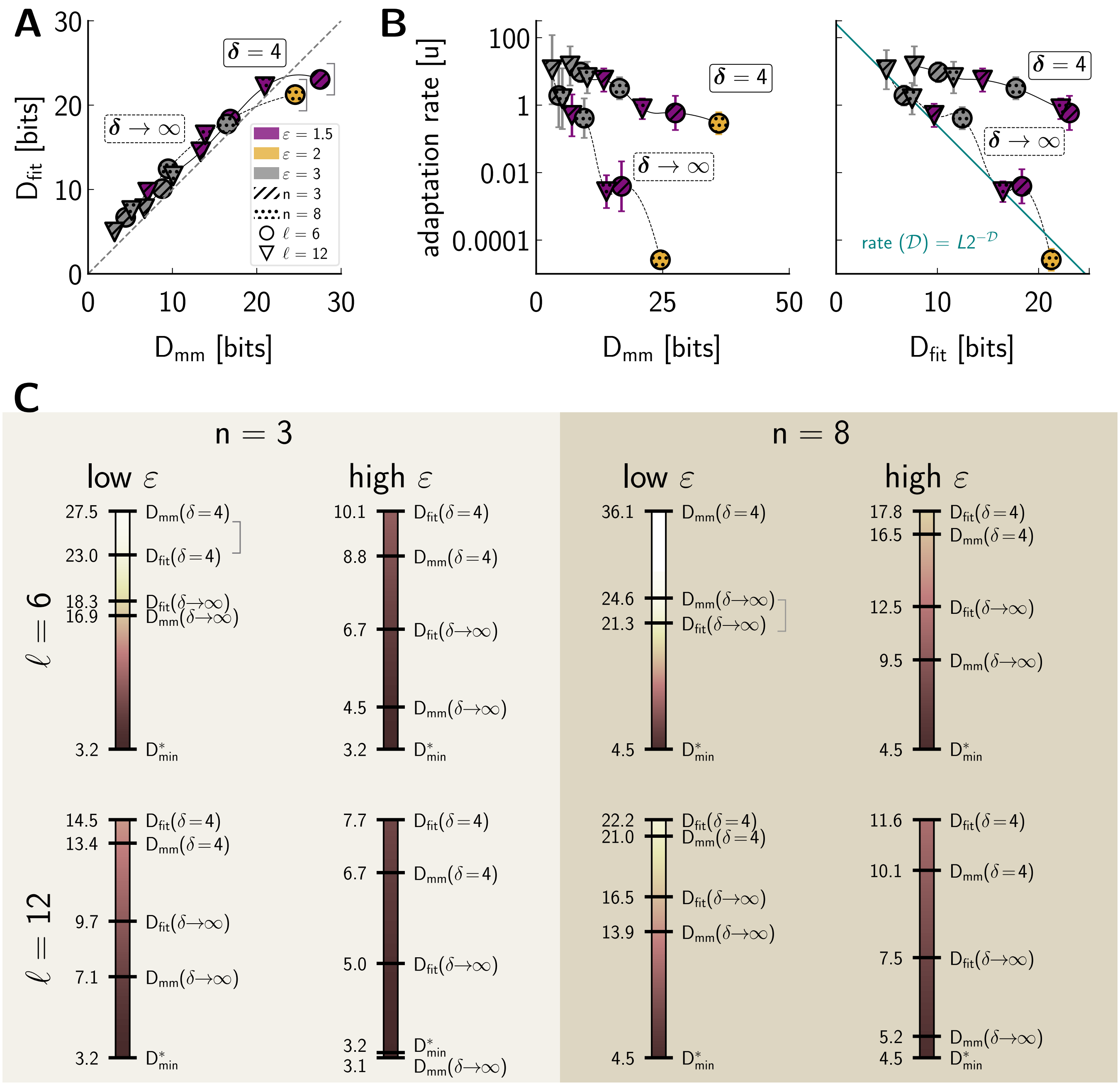

**Fig. S6. Various measures of accumulated information in adapted CRE ensembles.** We show results over the same parameters $(n, \ell, \varepsilon)$ as in Fig. S4. **(A)** Information on the fitness level is only slightly higher than information calculated on the mismatch level. Dashed line denotes $D_{\rm fit} = D_{\rm mm}$. Information values reported for the sharp threshold, $\delta \to \infty$ case (dotted line), are evaluated at $(\mu, c_{\max})$ parameters that are optimal for the $\delta = 4$ case; consequently, these are *not* the IT-optimal solutions for $\delta \to \infty$ and STLC regime. This has been done to isolate the effect on information of varying just $\delta$, the activation threshold steepness. Solutions indicated by the gray brackets deviate from the trend, likely because their neutral fitness distributions are undersampled even at $10^8$ samples. **(B)** CRE adaptation rate as a function of $D_{\rm mm}$ and $D_{\rm fit}$. Adaptation is faster, as hypothesized, when selection needs to shift the neutral distributions by fewer bits to favor fit solutions. Adaptation rates estimated from $\mathcal{D}$ (cyan line; right) still capture the trend in the $\delta \to \infty$ case even though here the adaptation happens at relatively high concentrations. Error bars show the standard error of the mean. **(C)** Ordering of the information measures (bits, numbers at left of the bars) for various sets of regulatory parameters. Gray brackets indicate the same values as in **A**. Lighter colors on the colorscale indicate higher information values overall; for example, outcomes with a high mismatch penalty $\varepsilon$ utilize fewer bits than with low $\varepsilon$ (compare low and high $\varepsilon$ columns at the same $(n, \ell)$ pair). Similarly, more crosstalking TFs necessitate higher information (compare results in light vs. dark background at the same $(\ell, \varepsilon)$ pair). Baseline parameters: $L = 256, R = 2, \delta = 4, \alpha = 1, N = 100$.

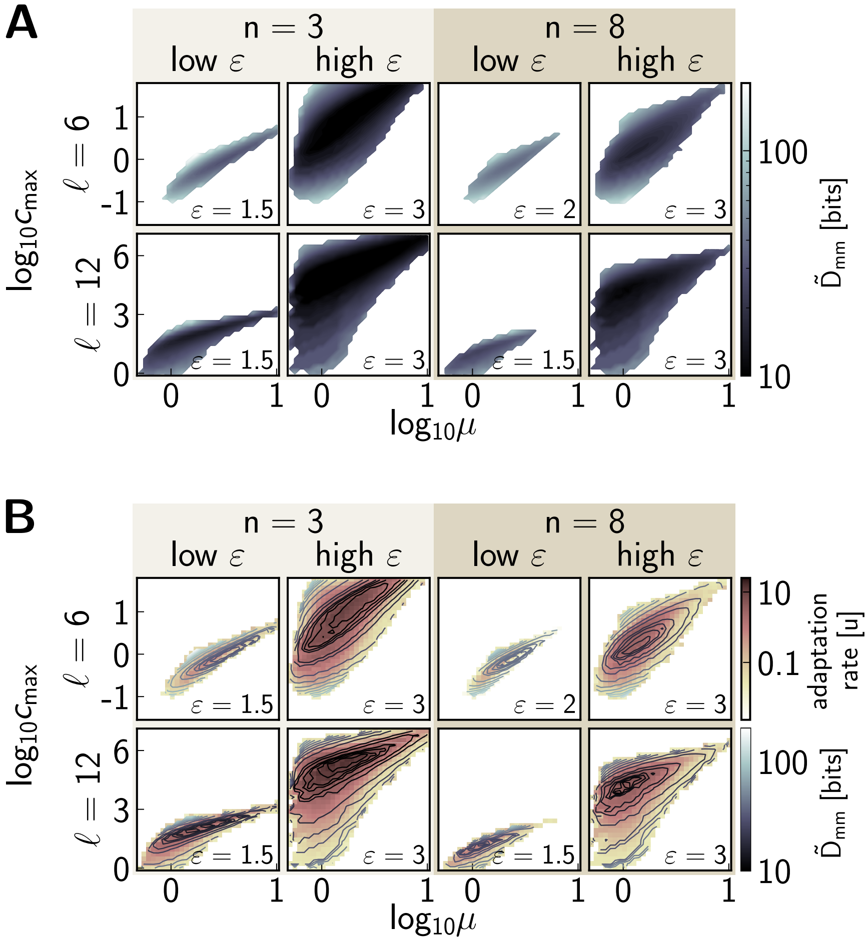

**Fig. S7. Information on the mismatch level predicts CRE evolutionary rates at optimal and suboptimal regulatory parameter combinations.** We show results over the same parameters $(n, \ell, \varepsilon)$ as in Fig. S4. **(A)** Mismatch level information (colorbar) as a function of regulatory parameters $(\mu, c_{\max})$. $\tilde{D}_{\mathrm{mm}}$ is based on the KL divergence between adapted and neutral mismatch distributions across an ensemble of CREs; mismatch distribution is estimated from one evolved CRE across (typically) 100 simulation replicates. This quick-to-compute measure that we can estimate across the entire $(\mu, c_{\max})$ plane approximates a more rigorous estimate, $D_{\mathrm{mm}}$, which is computationally more intensive and which we use everywhere else for the optimal solution (SI Appendix Section 2). **(B)** Adaptation rate (brownish colorbar) as a function of $(\mu, c_{\max})$, overlaid by the contours representing mismatch level information, $\tilde{D}_{\mathrm{mm}}$ (grayscale colorbar). The landscapes closely resemble each other, and the information measure correctly captures fast adaptation rates even on a smooth GP map. Baseline parameters: $L = 256, R = 2, \delta = 4, \alpha = 1, N = 100$.

 Réka Borbély, Michal Hledík and Gašper Tkačik

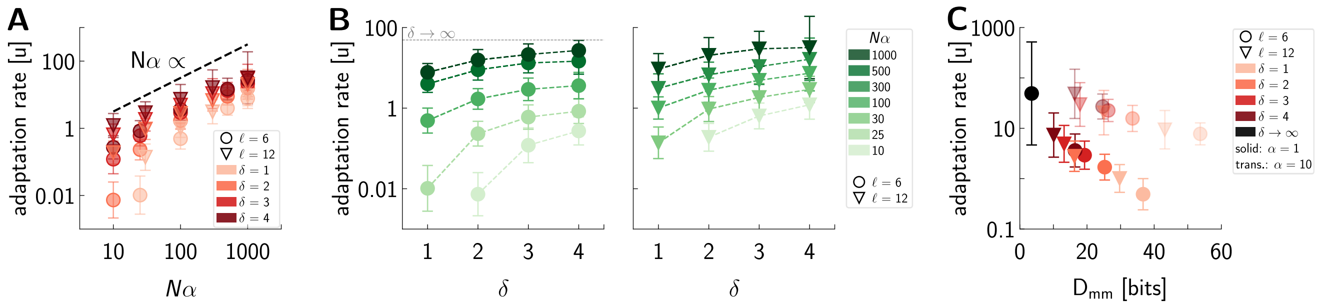

**Fig. S8. Adaptation rates at optimized solutions over a wide range of evolutionary parameters and activation nonlinearity steepness $\delta$. (A)** Optimal CRE adaptation rates as a function of effective selection strength, $N\alpha$, at various $\delta$ values (shade of red) for $\ell = 6$ (circles) and $12$ bp (triangles) long sites. Except for two parameter combinations at $(\ell, \delta, N\alpha) = (6, 1, 25)$ and $(6, 2, 10)$, we sampled the regime where adaptation rates scale with the effective selection strength, $N\alpha$. **(B)** Optimal CRE adaptation rates increase with $\delta$ at different effective selection strength values, $N\alpha$ (green, shaded), for $\ell = 6$ (circles, left) and $\ell = 12$ (triangles, right). Higher $N\alpha$ and higher $\delta$ facilitate faster adaptation; however, $N\alpha$ has a larger effect for smoother activation thresholds (low $\delta$), presumably because selection gradient is larger across a larger swathe of the genotype space. Increasing $N\alpha$ at low $\delta$ results in a higher jump in adaptation rates than increasing it at high $\delta$. For example, the adaptation rate increases $\sim 3$ orders of magnitude at $\ell = 6, \delta = 1$ when going from $N\alpha = 25$ to $1000$, but the same change in $N\alpha$ only results in $\sim 1$ order of magnitude rate change at $\ell = 6, \delta = 4$. As we increase the steepness of the activation threshold $\delta$, solutions approach the STLC regime (with $\delta \to \infty$ ), where the fitness landscape is piecewise flat, IT-efficient solutions exist for certain parameter combinations, overspecification is minimal, and adaptation rates become independent of $N\alpha$. The adaptation rate of this limit is indicated with the dotted gray line at $l = 6$. **(C)** Adaptation rate plotted against the mismatch level information $D_{\mathrm{mm}}$. Adaptation is faster when selection needs to accumulate less bits by "shifting" the neutral distribution towards the adapted one. As $\delta$ increases (shade of red) and the GP map becomes steeper, overspecification is reduced, fewer bits are needed and the adaptation proceeds faster. Black marker is the IT-efficient solution at $\delta \to \infty$ and $\ell = 6$. Transparent markers indicate optimized solutions at a higher $\alpha$ value (legend). Even though more information is accumulated at higher $\alpha$, adaptation nevertheless proceeds faster because selection is more efficient. Parameters: $L = 256, R = 2, n = 8, \varepsilon = 3, N = 100$. Markers represent averages, and error bars indicate standard deviations, over 100 replicate simulations. Exceptions: $(\ell, \delta, N\alpha) = (6, 1, 25)$ and $(6, 2, 10)$, where we only had $82$ and $92$ successfully adapted trajectories, in which case the adaptation rates may be slightly overestimated.

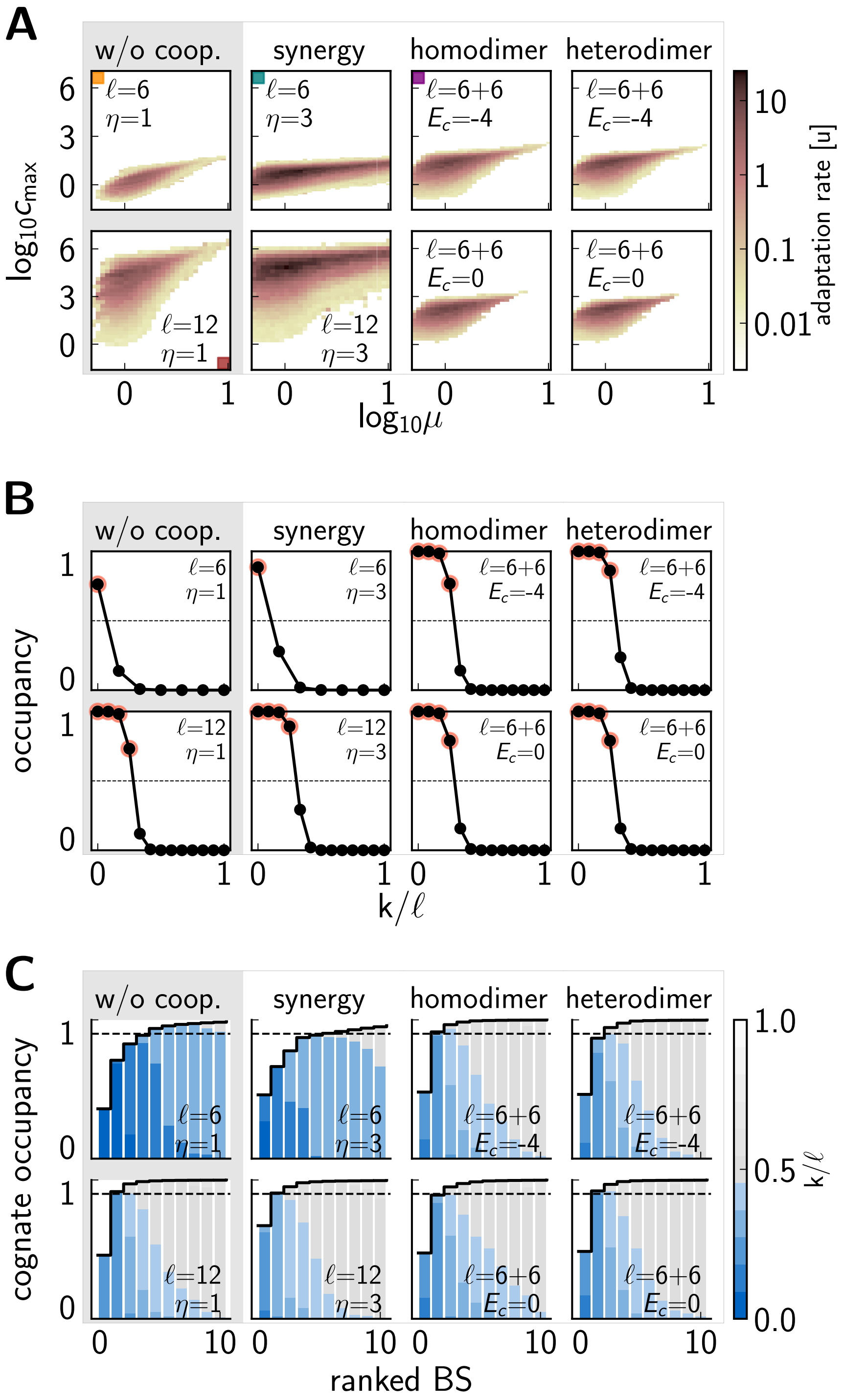

**Fig. S9.** **Evolutionary outcomes on GP maps with and without cooperative interactions.** We analyzed four different GP maps: (i) TFs bind as short ($\ell = 6$ bp) or long ($\ell = 12$ bp) monomers; (ii) TFs bind as monomers, ($\ell = 6, 12$ bp), and additionally the map features synergistic interactions among the same type of TFs ($\eta = 3$); (iii) TFs bind as homodimers ($\ell = 6 + 6$ bp) with the consensus being twice the same 6 bp motif either dimerizing on the CRE ($E_c = -4k_BT$) or already in the solution ($E_c = 0k_BT$); (iv) TFs bind as heterodimers ($\ell = 6 + 6$ bp) with different consensus for the two 6 bp binding domains, either dimerizing on the CRE ($E_c = -4k_BT$) or in the solution ($E_c = 0k_BT$). Each of these GP maps has a smooth threshold ($\delta = 4$) and, as shown in **A**, we optimize over the regulatory parameters $(\mu, c_{\max})$ separately for each scenario (i-iv). We display optimal solutions in **B-C** and in Fig. S10. **(A)** CRE adaptation rates as a function of regulatory parameters $(\mu, c_{\max})$, for different scenarios. CREs utilizing short binding sites optimally adapt at lower concentrations than CREs harboring long binding sites; this is the case for both monomer binding and synergistic binding. In contrast, dimerization enables optimal solutions at lower concentrations compared to equally long monomer scenario. Colored square markers indicate GP maps that we report on in the main text and main Fig. 6. **(B)** TF occupancy curves as a function of the normalized mismatch class, $k/\ell$, computed for the optimal $c^*_{\max}$ for each scenario. Red circles show strong sites that have $> 0.5$ occupancy (horizontal dashed line). **(C)** Fractional contributions to the cognate regulatory input, $\rho_s$, at the optimal regulatory parameters $(\mu^*, c^*_{\max})$. Evolved BSs are ranked by contribution (x-axis), color (bar at right) shows the normalized match to TF consensus (dark blue = full match, gray = complete mismatch), thick black line = cumulative input, dashed black = 90% threshold. Short motif length facilitates the emergence of multiple significant binding sites, whereas CREs with $\ell = 12$ or $\ell = 6 + 6$ function through significant binding sites. Baseline parameters: $L = 256$, $R = 2$, $\delta = 4$, $\alpha = 1$, $N = 100$.

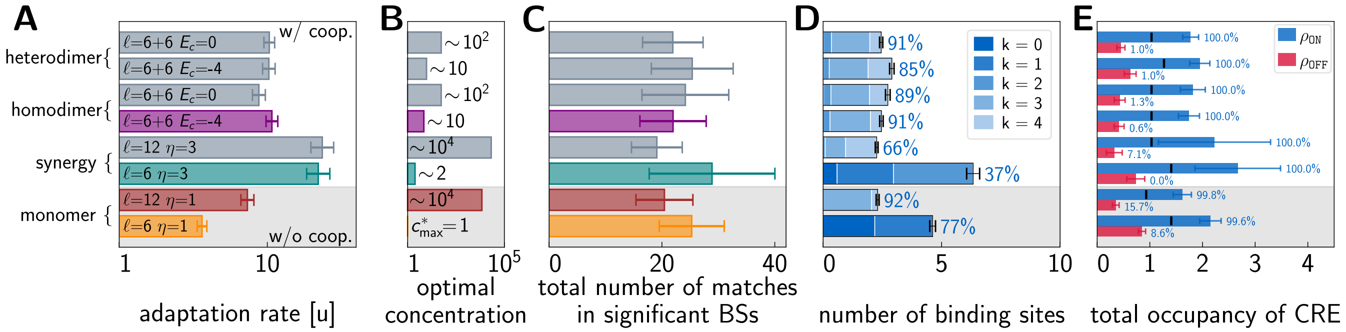

**Fig. S10.** **Properties of optimal solutions on realistic GP maps with and without cooperativity.** We show results over the same regulatory parameters as in Fig. S9. **(A)** CRE adaptation rate for six regulatory architectures with cooperativity and two architectures without, with regulatory parameters $(\mu, c_{\max})$ optimized separately for each architecture. Colored bars show the same results that we report on in the main text and main Fig. 6. **(B)** Optimal maximal TF concentrations, $c^*_{\max}$ normalized to the reference case of noncooperative monomer binding with $n = 8, \ell = 6, \varepsilon = 3$. Optimal solutions in architectures featuring dimerization occur at lower concentrations than solutions featuring the same length binding sites with synergy or without cooperativity. **(C)** The total number of matches in significant BSs (contributing $\sim 90\%$ of regulation) averaged over 100 replicates. To implement optimal regulation, different architectures (all at same $\epsilon = 3$ used here) all must fix approximately the same number of matches. **(D)** Structural details of evolved CREs for different parameters. Bars show the number of BSs contributing significantly to total regulatory input, broken down by the number of mismatches (bluish hue; legend). Percentages show the fraction of total regulatory input accounted for by BSs with occupancy $> 0.5$ at optimally chosen $c_{\max}$. CREs harbor either 2-3 long sites or 5-7 short sites. Synergistic interactions facilitate the emergence of multiple weaker sites. **(E)** Absolute values of total occupancy in the ON and OFF environment, $\rho_{\texttt{ON}}$ (blue) and $\rho_{\texttt{OFF}}$ (red). Percentages show the fraction of total regulatory input due to the binding of the cognate TF. Little black markers indicate the optimal activation threshold $\mu^*$. Baseline parameters: $L = 256, \varepsilon = 3, R = 2, \delta = 4, \alpha = 1, N = 100$. In (**A, C, D, E**) we report on averages over 100 replicates. Error bars in (**A, D**) are standard errors of the mean; in (**C, E**) the standard deviation.

### 6. List of tables

**Table S1. Parameter values on four different genotype-phenotype maps (color; legend at right).**

| Type | Parameter | Value |
|---|---|---|
| Combinatorial | CRE length, $L$ | 256 bp |
| | | 256 bp |
| | | 256 bp |
| | | 256 bp |
| | motif length, $\ell$ | 6,7,8,10,12 bp |
| | | 6 vs. 12 bp |
| | | 6+6 bp |
| | | 6 vs. 12 bp |
| | # noncognate TFs, $n$ | 3 |
| | | 3 vs. 8 |
| | | 8 |
| | | 8 |
| Biophysical | mismatch penalty, $\varepsilon$ | 1.5, 2, 2.5, 3 $k_BT$ |
| | | 1.5/2 vs. 3 $k_BT$ |
| | | 3 $k_BT$ |
| | | 3 $k_BT$ |
| | concentration, $c_{max}$ | 0.1 |
| | | optimized |
| | | optimized |
| | | optimized |
| | activation threshold, $\mu$ | optimized |
| | | optimized |
| | | optimized |
| | | optimized |
| | slope of threshold, $\delta$ | $\infty$ |
| | | 1, 2, 3, 4 |
| | | 4 |
| | | 4 |
| | cooperative coefficient | - |
| | | - |
| | | $E_c$ = 0 vs. -4 $k_BT$ |
| | | $\eta$ = 3 |
| Evolutionary | population size, $N$ | 100 |
| | | 40 vs. 100 |
| | | 100 |
| | | 100 |
| | selection strength, $\alpha$ | 0.25, 1, 5 |
| | | 1; (0.0001, . . . , 25) |
| | | 1 |
| | | 1 |

| GP map features |
|---|
| Monomer binding in the STLC limit ($\delta \to \infty; c_{max} \ll 1$) |
| Monomer binding and smooth threshold |
| Dimerization |
| Synergistic interactions |
